# P53 deficiency triggers hypertranscription, inducing nucleotide insufficiency causing replication stress, and genomic instability

**DOI:** 10.1101/2025.04.17.649416

**Authors:** Wisam Zaatra, George Philippos, Petr Smirnov, Shira Milo, Jan Otonicar, Michelle Chan, Michal Harel, Frauke Devens, Tchelet Goldberg, Alexandra Eliassaf, Karen Grimes, Michal Irony-Tur Sinai, Gianluca Sigismondo, Tamar Geiger, Jan O. Korbel, Jeroen Krijgsveld, Ori Shalev, Batsheva Kerem, Aurelie Ernst

## Abstract

P53 prevents DNA damage by inducing repair processes, cell cycle arrest or apoptosis. P53 loss leads to replication stress and genomic instability, yet the mechanisms underlying these effects and their contribution to catastrophic genomic events such as chromothripsis remain poorly understood. Using patient-derived fibroblasts with germline p53 variants, that spontaneously undergo chromothripsis, and p53-downregulated fibroblasts, we discovered that p53 loss leads to aberrant transcriptional upregulation, increasing nucleotide consumption while simultaneously decreasing nucleotide biosynthesis. This imbalance in production and consumption results in insufficient nucleotide pools, leading to replication stress and genomic instability, which are rescued by nucleoside supplementation or transcription normalization. The replication stress triggers telomere dysfunction, micronuclei formation, and ultimately chromothripsis. Emerging dominant chromothriptic clones exhibit normal DNA replication, telomere stabilization, and ecDNA, highlighting critical features for clonal selection. Hence, p53 coordinates transcription and nucleotide pools, crucial for maintaining genomic stability and preventing early cancer development.

## Introduction

Genome instability is both a hallmark and a driving force of cancer development, increasing the likelihood of accumulating point mutations and structural variants. The latter include rearrangements such as deletions, amplifications, translocations and insertions, which collectively generate the majority of cancer drivers^1^. Next-generation sequencing technologies and their applications in cancer genomics have enabled the discovery of a complex form of genome instability called chromothripsis^2,3^. In contrast with the classical view of multi-step tumor development, this type of catastrophic event generates massive chromosomal rearrangements on one or a few chromosomes^2,3^, which frequently play a causative role in tumor development^4^.

Modeling chromothripsis in cell culture systems has shed light on putative mechanisms underlying chromothripsis^5–8^. A widely used cellular model involves the induction of micronuclei formation, a known marker for complex chromosomal rearrangements, including chromothripsis^6,7^. However, the molecular basis underlying spontaneous chromothripsis events in human cells remain largely unknown. It remains unclear to what extent mechanisms observed in artificially induced chromothripsis *in vitro* accurately mirror chromothripsis events in human tissues. Capturing the early stages of cancer evolution in humans is challenging, and the sequence of events leading to chromothripsis, as well as how these events confer selective advantages, remains elusive. Thus, our current understanding of how chromothripsis is triggered comes indirectly from genomic data analysis of fully developed tumors, long after the chromothriptic events have occurred, and from artificially induced chromosome segregation errors in cell lines. These studies revealed a strong link between the loss of the tumor suppressor p53 and chromothripsis occurrence^6,9–12^. In particular, the prevalence for chromothripsis was shown to reach almost 100% in a number of tumor types that develop in Li Fraumeni Syndrome (LFS) patients, which carry p53 germline variants^12,13^. Mechanistic studies on externally induced chromothripsis were performed in cells lacking functional p53^6,7,14^, raising the possibility that the loss of p53 function contributes to the induction of chromothripsis. The p53 tumor suppressor is activated by cellular stresses such as DNA damage, oncogene activation or hypoxia^15–17^. Mutations or deletions of p53 occur in over 50% of human cancers^18^, while in many other cases, p53 function is inactivated through other mechanisms^19^, showing that the functional inactivation of the p53 pathway is a critical step in tumor development. The high risk of early cancer onset in LFS patients, together with findings from studies in p53⁺/⁻ mice and cell lines, indicate that a single wild-type (WT) p53 allele is not sufficient to preserve normal cellular functions^20–22^.

One proposed mechanism for the induction of genomic instability following p53 loss involves perturbed DNA replication dynamics^23,24^. Defects in DNA replication dynamics are characterized by slowed fork progression and dormant origin activation (increased numbers of active origins)^25–27^. The latter is a compensatory mechanism enabling the completion of DNA replication during S-phase. Previous studies in cancer cell lines have demonstrated that p53 plays a role in regulating dormant origin firing, restarting of stalled replication forks and preventing replication–transcription collisions^23,24^. However, the molecular basis underlying the replication stress following p53 loss, as well as the role of p53 loss in inducing chromothripsis, is not well understood.

In this study, we reveal that loss of WT p53 function induces hypertranscription, which induces nucleotide insufficiency by enhancing transcription-driven nucleotide consumption, which together with reduced nucleotide production, leads to replication stress. This stress drives DNA damage, telomere shortening, micronuclei formation and chromothripsis. We further demonstrate that nucleotide supply or transcription normalization rescues replication stress and genomic instability, highlighting the critical role of nucleotide availability in genome maintenance upon p53 loss. We further show the relevance of this mechanism for cancer development driven by genomic instability, in particular in the context of chromothripsis. By directly dissecting the mechanisms underlying spontaneous chromothripsis, we reveal that early events include mild replication stress, characterized by reduced replication rates, activation of dormant origins, DNA double-strand breaks, and micronuclei formation, along with high numbers and highly diverse chromothriptic events, all of which are intensified following p53 loss close to the growth crisis. We also identify clonal selection processes that support cell survival after chromothripsis, including telomere stabilization and amplification of extrachromosomal DNA (ecDNA). Altogether, our findings uncover a novel role for p53 in coordinating transcription and nucleotide pools, which are crucial for maintaining genomic stability and preventing early cancer development.

## RESULTS

### DNA damage foci and nuclear atypia in early-passage LFS cells, with further aggravation during crisis

To investigate cellular processes underlying genomic instability, and in particular chromothripsis occurrence, we used primary fibroblasts from LFS patients, which spontaneously enter a growth crisis characterized by slower growth, from which a subset of chromothriptic cells escapes and becomes immortalized (**Fig. 1a, Supplementary Information, Fig. S1a-c**)^28^. First, we performed a longitudinal profiling of genomic instability and phenotypic features associated with chromothripsis, such as DNA double-strand breaks^5^, chromatin bridges^8,29^, mitotic defects^5,7^ and micronuclei^5,6^ (i.e., nuclear atypia encapsulating chromosomal fragments, or up to a few chromosomes). Quantification of γH2AX foci showed many DNA double-strand breaks at early passages, which significantly increased at crisis passages, but decreased at late passages to significantly lower levels relative to the early passages (**Fig. 1b**). Chromatin bridge quantification showed similar kinetics, with high levels detected at early passages, increased levels during the crisis, but significantly lower levels at late passages (**Fig. 1c, d**, patient LFS041 and **Supplementary Information, Fig. S1d**, patient LFS087). In line with this, strand-specific single-cell DNA sequencing (strand-seq)^30^ showed fold-back inversions followed by terminal deletions characteristic of breakage-fusion-bridge cycles in a high fraction of cells (up to 70% on chromosome 5p) already at early passages (**Fig. 1e**).

**Figure 1.**
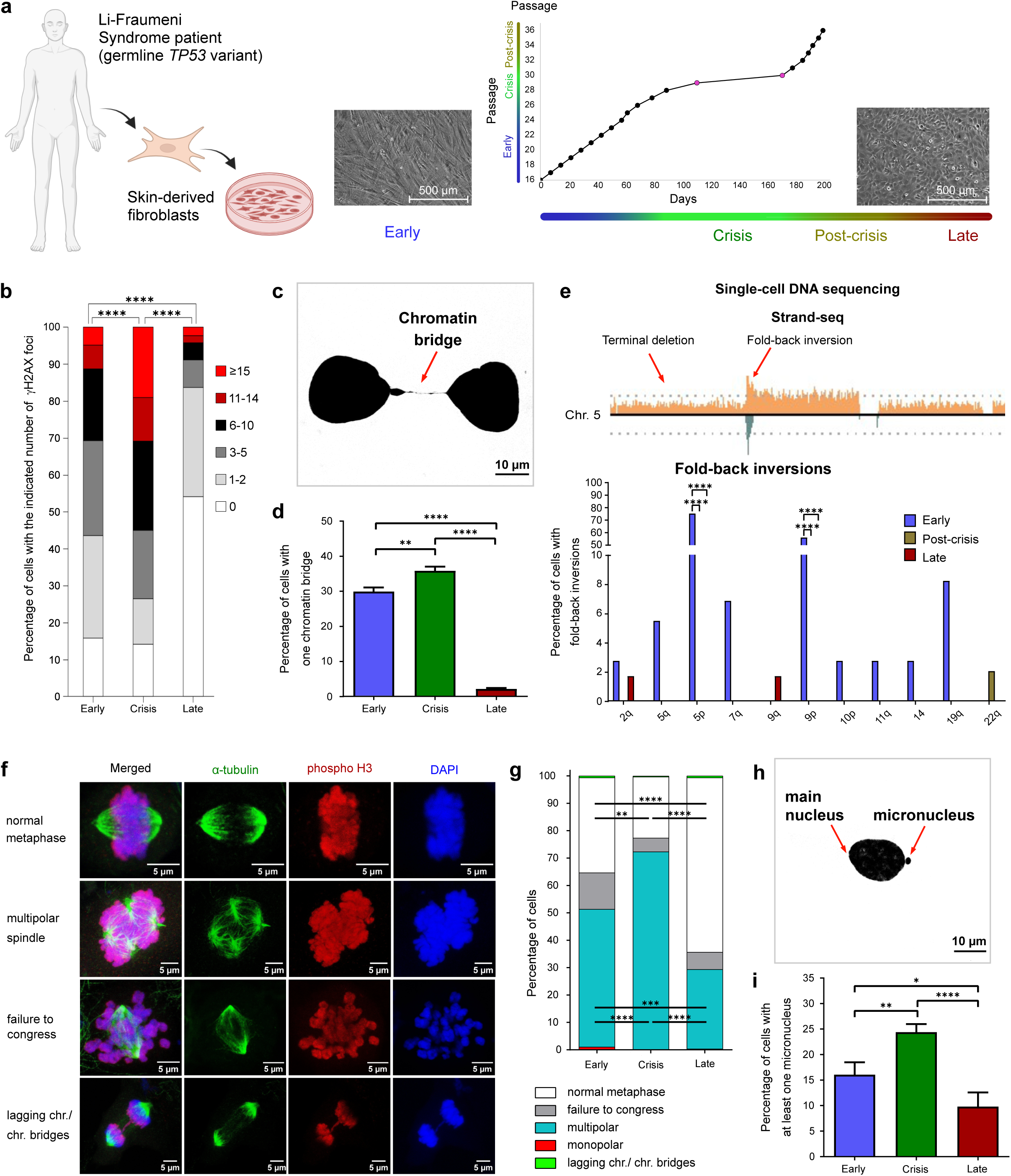
Phenotypic profiling identifies chromatin bridges, multipolar spindles and micronuclei already at early passages. **a.** Scheme describing the LFS model system. The growth curve depicts the time required for cells to reach confluency (x-axis) from passage 16 (early passage), passing through the growth crisis, and continuing to passage 36 (post-crisis) (y-axis) for patient LFS041. Black circles: splitting ratio 1:3; purple circles: splitting ratio 1:2. Representative images showing the morphology of LFS patient fibroblasts at early and late passages are shown. Images were taken using a 5x magnification objective. Scale bar: 500 µm. BioRender.com was used to create this panel. **b.** Percentage of nuclei with γH2AX foci. The quantification was done in early-passage cells (mean foci number: 4.7 ±0.3; n=410), crisis cells (8.8 ±0.5; n=415) and late-passage cells (1.7 ±0.2; n=622). The data show a summary of three independent experiments. Statistical analysis was performed using Mann Whitney rank-sum test done on the distribution. **c.** Representative picture of a chromatin bridge between two cells. Scale bar: 10 µm. **d.** Quantification of chromatin bridges. Four independent biological replicates were performed for each condition: early (mean: 29.7 ±1.1), crisis (mean: 35.6 ±1.2), and late (mean: 2 ±0.4). For each replicate, 750 cells were quantified. **e.** Upper panel, representative example of a fold-back inversion followed by a terminal deletion on chromosome 5, which is characteristic of Breakage-Fusion-Bridge (BFB) cycles (Strand-seq data). Orange and green represent Watson (forward/plus) and Crick (reverse/minus) strands, respectively. Lower panel, quantification of fold-back-inversions in early, post-crisis and late passages from strand-seq data. **f.** Representative immunofluorescence pictures of a normal metaphase and different types of mitotic defects. Scale bar: 5 µm. **g.** Quantification of mitotic defects. Three independent biological replicates were performed for each condition. For each replicate, 100 cells were quantified. Lagging chr./chr. bridges: lagging chromosomes/ chromatin bridges. **h.** Representative picture of a micronucleus. Scale bar: 10 µm. **i.** Quantification of micronuclei. Four independent biological replicates were performed for each condition: early (mean: 16.1 ±1.2), crisis (mean: 24.4 ±0.8), and late (mean: 9.8 ±1.4). For each replicate, 750 cells were quantified. In **e**, Fisher’s exact test was performed for pairwise comparisons between each of the three passages. P-values were adjusted for multiple comparison tests. In **g,** data are presented as mean values. Statistical significance was assessed using repeated-measures two-way ANOVA followed by uncorrected Fisher’s LSD for multiple comparisons. In **d** and **i,** data are presented as mean ± SEM. Statistical significance was assessed using a one-way ANOVA followed by Tukey’s multiple comparisons test. P-values less than 0.05 were considered statistically significant. (*p < 0.05, **p < 0.01, ***p < 0.001, ****p < 0.0001).

In inducible model systems, broken bridge chromosomes were reported to undergo mitotic DNA damage and frequent mis-segregation to form micronuclei^5^. To explore the role of mitotic errors as a source of spontaneous chromothriptic events, we quantified mitotic defects using immunofluorescence analysis of phosphorylated histone H3 and acetylated tubulin in fibroblasts from patients LFS041 and LFS087 (**Fig. 1f** and **Supplementary Information, Fig. S1e**). We detected high levels of mitotic defects at early passages and during the crisis, and in particular a large number of multipolar spindles (**Fig. 1g** and **Supplementary Information, Fig. S1f)**, which can lead to chromosomal instability^31^. Importantly, multipolar spindles cause lagging chromosomes and mis-segregation and are a source of micronuclei^31^. In line with this, we found high numbers of micronuclei at early passages and an increase during the crisis (**Fig. 1h, i** and **Supplementary Information, Fig. S1g**). The fraction of cells with mitotic defects and micronuclei decreased at late passages (**Fig. 1f-i)**. Altogether, phenotypic profiling of canonical hallmarks linked with chromothripsis indicates that the LFS cellular system is well-suited to study the mechanistic basis of spontaneous chromothripsis and this profiling also identified critical time points for dissecting clonal evolution using single-cell genomics.

### High numbers and high diversity of chromothriptic events already at early passages and later selection of dominant clones

Next, we used strand-seq to investigate the timing of structural variant formation and how clones with different types of alterations evolve over time. Even at early passages, cells exhibited a high number and broad diversity of structural rearrangements (averaging 20 per cell, predominantly deletions), indicating that they had already undergone cellular stress by this stage (**Fig. 2a** and **Supplementary Information, Fig. S2a**). In contrast, post-crisis cells (39 structural variants per cell on average) and late-passage cells (53 structural variants per cell on average) exhibited more clonal events, suggesting that they have undergone a selection bottleneck (**Fig. 2a**). The decreasing diversity from early to post-crisis and the appearance of dominant clones was supported by lineage tree reconstruction (**Fig. 2b** and **Supplementary Information, Fig. S2b**). The distribution of pairwise distances between cells (**Supplementary Information, Fig. S2c**) and phylogenetic diversity estimates - quantified by adding the tree branch lengths in each group - showed that post-crisis cells (p63) were genetically closely-related to each other, whereas early-passage cells (p22) showed more diversity (phylogenetic diversity: 5072 at P22, 2791 at P63) (**Fig. 2b**). The increase in diversity in (very) late-passage cells (p349) potentially reflects the accumulation of alterations after long-term culturing (phylogenetic diversity: 5199 at P349).

**Figure 2.**
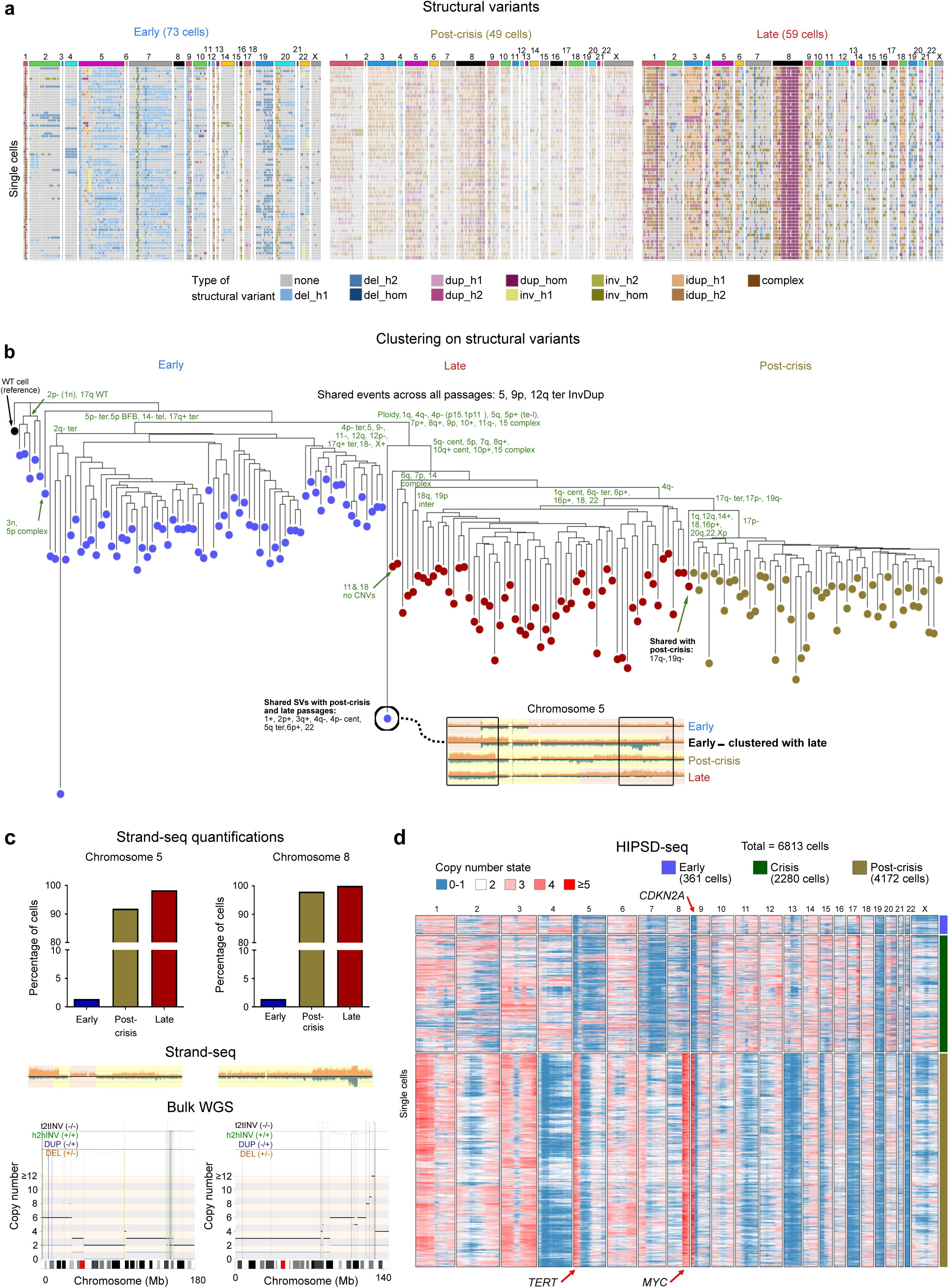
Single-cell DNA sequencing shows a high number and high diversity of chromothriptic events at early passages and later selection of dominant clones. **a.** Heatmaps showing structural variants per chromosome occurring in at least 10% of the cells within each passage (strand-seq data). Each row shows one cell. Deletions are depicted in blue, amplifications and complex rearrangements in orange and brown, respectively. Del: deletion, dup: duplication, inv: inversion, idup: inverted duplication, h1: homolog 1, h2: homolog 2, hom: homologous. **b.** Clustering on structural variants detected in strand-seq data, using an event-based modified Hamming distance and neighbour-joining **(Methods)**. Structural variants separating the main cell clusters are shown on the nodes. Shared events are annotated in black, while differences between the main clusters are annotated in green. Bottom, chromosome 5 from representative early-passage, post-crisis and late-passage cells are shown. Abbreviations: -: loss, +: gain, cent: centromeric, tel: telomeric, term: terminal. The distance between cells reflects the number of events. **c. Upper panels:** Quantifications of the frequency of complex rearrangements on chromosomes 5 and 8 across passages from LFS041 based on strand-seq data. **Middle panels:** Representative examples of complex rearrangements in the strand-seq data for chromosomes 5 and 8. **Lower panels:** Copy-number plots for chromosomes 5 and 8 from bulk WGS (LFS041 p.63, post-crisis passage). t2tINV: tail-to-tail inversion (black), h2hINV: head-to-head inversion (green), DUP: duplication (blue) and DEL: deletion (orange). **d.** HIPSD-seq heatmaps (bin size of 1000 kb) of passage p.22 (early, n=361 cells), p.27 (crisis, n=2280 cells) and p.62 (post-crisis, n=4172 cells) showing copy-number variation. Each row represents one cell. Copy number profiles of post-crisis cells were corrected for ploidy.

The vast majority of the complex rearrangements detected at early passages appeared not to have strong selective advantages. Interestingly, we detected one early-passage cell that clustered with the late-passage cells, as a putative descendant of the same precursor cell which gave rise to the dominant subclone seen in the late-passage cells (**Fig. 2b**). Consistent with this, complex rearrangement events detected as clonal in late-passage cells and detectable by bulk whole genome sequencing (WGS) were observed only in minor subpopulations of early-passage cells (**Fig. 2c** and **Supplementary Information, Fig. S2d, e**). This implies that fewer than 5% (**Fig. 2c**) of early-passage cells give rise to the dominant clones observed at later stages. Common events between early and late-passage cells included alterations on chromosome 5, loss on chromosome 9p and terminal inverted duplications on chromosome 12q (**Fig. 2b** and **Supplementary Information, Fig. S2c**).

Chromothripsis has been previously linked with genome doubling^13,14^. However, the order by which chromothripsis and polyploidization occur, inferred from cancer genomes, can vary. In this patient-derived fibroblast model, only 1 out of 73 early-passage cells was polyploid (ploidy inference from strand-seq, see **Methods**), while almost a third (24/73) had at least one complex rearrangement. At late passages, all analyzed cells were polyploid (49/49, passage 63), as confirmed by strand-seq, metaphase spread and WGS analysis (in LFS041 and LFS087 cells) (**Supplementary Information, Fig. S3**). Hence, in this model system, chromothripsis seems to precede genome doubling.

To analyze copy-number variants using a high-throughput method capable of detecting rare events and also applicable to crisis passage cells (the low proliferation rate of the cells during the crisis was not compatible with the requirement for BrdU incorporation for strand-seq analysis), we applied HIgh-throughPut Single-cell DNA-sequencing (HIPSD) on patient LFS041 cells (early, crisis and late passage) (**Fig. 2d**) and patient LFS087 cells (late passage) (**Supplementary Information, Fig. S2f**)^32^. The copy-number profiles inferred from HIPSD-seq were largely consistent with the strand-seq data, showing a high number of rearrangements and high genetic diversity in early-passage and crisis cells but more clonal events at later passages (**Fig. 2d**). Importantly, the high clonal diversity in early-passage cells compromised the ability of bulk WGS to detect copy-number variants, as alterations from distinct clones masked one another, resulting in the appearance of seemingly balanced regions (**Supplementary Information, Fig. S4a–d**) (see also bulk WGS of LFS041 early-passage cells in our previous work^28^). Clonal events detected already at early passages and potentially facilitating chromothripsis (and/or the survival of chromothriptic cells) included the loss of chromosome 9p containing the *CDKN2A* locus, encoding for an essential regulator of cell growth, cell division and cell death (**Fig. 2d**). In post-crisis passage cells, clonal focal gains likely explaining the selective advantage and clonal expansion included the locus of the oncogene *MYC* (**Fig. 2d**). In addition, the locus encoding for telomere reverse transcriptase (*TERT*), located on chromosome 5p was clonally gained in post-crisis cells, even though a loss was detected at earlier passages (**Fig. 2d**), suggesting that BFB cycles on chromosome 5p may promote chromothripsis, followed by stabilization via *TERT* gain. Taken together, genomic profiling identified potential drivers that may contribute to clonal selection.

### Telomere stabilization occurs near the crisis and promotes the emergence of dominant clones

We set out to investigate how dominant clones at late passages avoid senescence and apoptosis, achieve telomere stabilization and acquire specific drivers. Telomere stabilization via reactivation of TERT or by Alternative Lengthening of Telomeres (ALT) appeared as a requirement in dominant clones. Whereas early-passage LFS041 cells only rarely showed gains of *TERT*, we detected cells with high copy number of *TERT* after the crisis and at late passages, as shown by *TERT* FISH across passages (**Fig. 3a, b**). In fibroblasts from patient LFS087, which acquired telomere stabilization through ALT, ALT-positive clones also emerged from the crisis onwards (**Fig. 3c, d**). Early-passage cells in LFS patients already have shorter telomeres as compared to healthy donors, promoting genomic instability, and telomere stabilization potentially prevents telomere fusions and further shattering of the genome in such cells^31^. Clonal telomere stabilization after the crisis likely explains the reduction in the number of chromosome bridges that we observed in late-passage cells (see **Fig. 1c, d**) and appears to be a critical driver for chromothriptic clones to become dominant.

**Figure 3.**
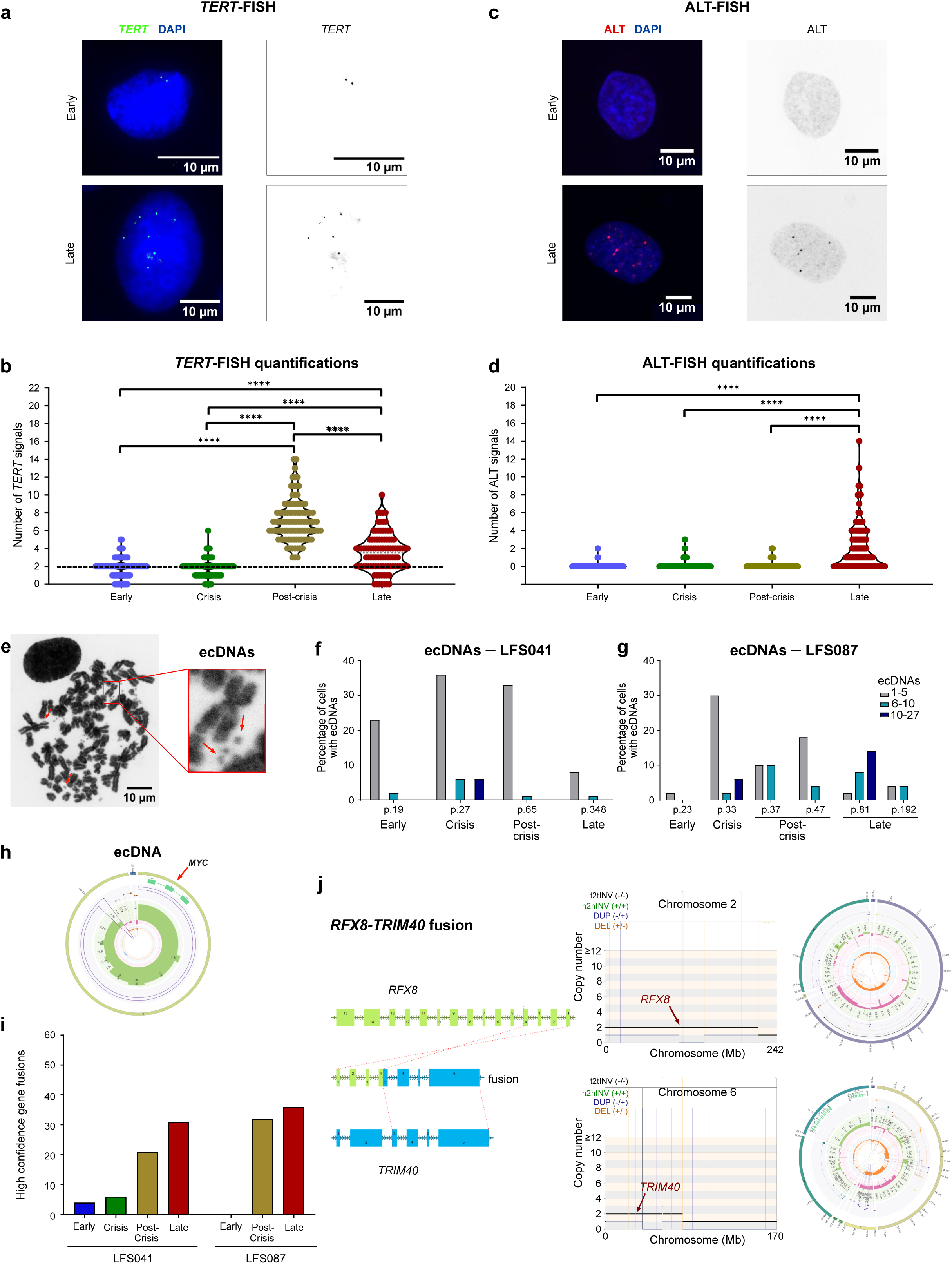
Genomic features of the dominant clones. **a.** Representative pictures of *TERT*-FISH from early and late passages. Scale bar: 10 µm. **b.** Quantification of *TERT*-FISH signals per nucleus in patient LFS041: early (mean ± SEM: 1.8 ±0.1), crisis (mean ± SEM: 1.8 ±0.1), post-crisis (mean ± SEM: 7.1 ±0.2), late (mean ± SEM: 3.5 ±0.2). 100 nuclei per passage were quantified. Dashed line represents diploid *TERT* levels (2 copies per cell). **c.** Representative pictures of ALT-FISH from early and late passages. Scale bar: 10 µm. **d.** Quantification of ALT-FISH signals per nucleus in patient LFS087: early (mean: 0.03 ±0.01), crisis (mean: 0.10 ±0.03), post-crisis (mean: 0.11 ±0.03), and late (mean: 2.00 ±0.17). Two independent biological replicates were performed for each condition. For each replicate, 100 cells were quantified. **e.** Representative picture of LFS087 p.81 (post-crisis) metaphase spreads showing high numbers of ecDNAs. Scale bar: 10 µm. **f, g.** Quantification of the percentage of cells with ecDNAs. 50 cells per passage were quantified for each patient. **h.** Example of ecDNA structure detected in LFS041 (bulk WGS) containing the oncogene *MYC*. ecDNAs were identified using OncoAnalyser and verified using AmpliconSuite. **i.** Number of high-confidence gene fusions identified from bulk RNA sequencing. **j.** Representative example of the *RFX8-TRIM40* gene fusion in LFS087 p.195 (late passage). Left panel, breakpoint sites of both genes involved in the fusion, namely *RFX8* (chromosome 2) and *TRIM40* (chromosome 6). Middle panels: ReConPlot of Chromosomes 2 and 6 from bulk WGS of LFS087 p.195 (late passage) highlighting *RFX8* and *TRIM40I*, respectively, and showing structural variants on both chromosomes. Color coding indicates sites of deletions (orange), duplications (blue), head-to-head inversions (green) and tail-to-tail inversions (black). t2tINV: tail-to-tail inversion, h2hINV: head-to-head inversion, DUP: duplication and DEL: deletion. Right panels: Circos plots from bulk WGS of LFS087 (p.195) showing structural variations supporting the gene fusion detected by RNA sequencing. In **b** and **d,** statistical significance was assessed using a nonparametric Kruskal-Wallis test, followed by Dunn’s multiple comparisons test. P-values below 0.05 were considered statistically significant (*p < 0.05, **p < 0.01, ***p < 0.001, ****p < 0.0001).

### Formation of extrachromosomal circular DNAs and gene fusions contributes to the emergence of dominant clones

Next, we investigated whether extrachromosomal circular DNAs (ecDNAs) and/or gene fusions generated by chromothripsis contribute to the selective advantages in dominant clones. We detected low numbers of ecDNAs at early passages, which were further increased at crisis and post-crisis passages as compared to early passages, indicating more ecDNAs in the dominant clones (**Fig. 3e-g**). Assembly from bulk WGS data confirmed the presence of ecDNAs harboring driver genes (e.g. *MYC*) at late passages (**Fig. 3h** and **Supplementary Information, Fig. S5a-d**). At very late passages (after p150), the decreasing number of detected ecDNA structures may potentially be explained by re-integration of ecDNAs into linear chromosomes, as reported previously^33^. Altogether, ecDNAs appear to play a critical role to help cells overcome the crisis.

In addition to ecDNAs, chromothripsis was shown by us and by others to generate cancer drivers via gene fusions^3,34^. Using gene fusion detection from bulk RNA sequencing (see **Methods**), we identified very low numbers of gene fusions (less than 10 high-confidence fusions per patient) at early passages, as expected due to the absence of clonal chromothriptic event at this stage. In contrast, we detected an increased number of gene fusions from the crisis onwards (**Fig. 3i, j** and **Supplementary Information, Fig. S5e-g**). For instance, chromothripsis resulted in the formation of a *TRIM40-RFX8* fusion, leading to a 100-fold overexpression of the *TRIM40* gene—normally barely expressed in healthy individuals but described as a pathogenic driver in epithelial cells^35^.

### Loss of WT p53 function precedes the crisis stage in LFS cells

To better understand the role of p53 in chromothripsis occurrence, we examined the p53 status at the crisis stage for both LFS patients (LFS041 and LFS087), at both protein level by Western blot using the DO1 p53-specific antibody, and DNA level by DNA sequencing. In LFS041 cells, DNA sequencing analysis identified heterozygosity for the WT and the germline variant. However, protein analysis revealed the loss of WT p53 protein at the crisis, coinciding with the emergence of a truncated p53 variant (**Supplementary Information, Fig. S6a**). We confirmed that this short p53 variant was the p53 β isoform (P53β) using specific primers detecting the inclusion of intron 9β into the mature p53 mRNA (**Supplementary Information, Fig. S6b**). In patient LFS087, a loss of the WT allele at the DNA level was detected at the crisis, leaving the germline missense allele that was translated to a p53 protein with the expected molecular weight (**Supplementary Information, Fig. S6c**). To confirm that crisis LFS cells from both patients lost the WT p53 function, we analyzed the expression of two canonical p53 target genes (*p21* and *RRM2B*) by real-time quantitative PCR following treatment with the MDM2 antagonist Nutlin-3a, which is expected to activate target genes in the context of WT p53. As expected, p53-proficient BJ cells treated with 10µM Nutlin-3a for 48hrs showed a strong increase in the expression of *p21* and *RRM2B* mRNA levels. In contrast, crisis LFS041 and LFS087 cells showed no detectable increase in the expression of p53 target genes following Nutlin-3a treatment (**Supplementary Information, Fig. S6d**), indicating that crisis LFS cells have already lost their WT p53 protein activity (**Supplementary Information, Fig. S6e**). Altogether, these analyses confirmed the loss of WT p53 at the crisis stage.

### Loss of p53 reduces nucleotide metabolism and enhances translational activity

We further aimed to understand the contribution of WT p53 loss to the observed genome instability, which peaks during the crisis stage following p53 loss. Using bulk RNA-sequencing analyses across the various passages of LFS cells, we observed that replication stress and hypertranscription increase in LFS cells during the crisis as compared to early passages (**Fig. 4a-c**, patient LFS041). In particular, gene set enrichment and gene ontology (GSEA GO) analysis of transcriptomic data identified increased ‘DNA replication initiation’ (GO:0006270) and ‘DNA-binding transcription activator activity, RNA polymerase II-specific’ (GO:0001228) as major processes activated in crisis cells as compared to early passages (**Fig. 4a-c**, **Supplementary Information, Fig. S7c**).

**Figure 4.**
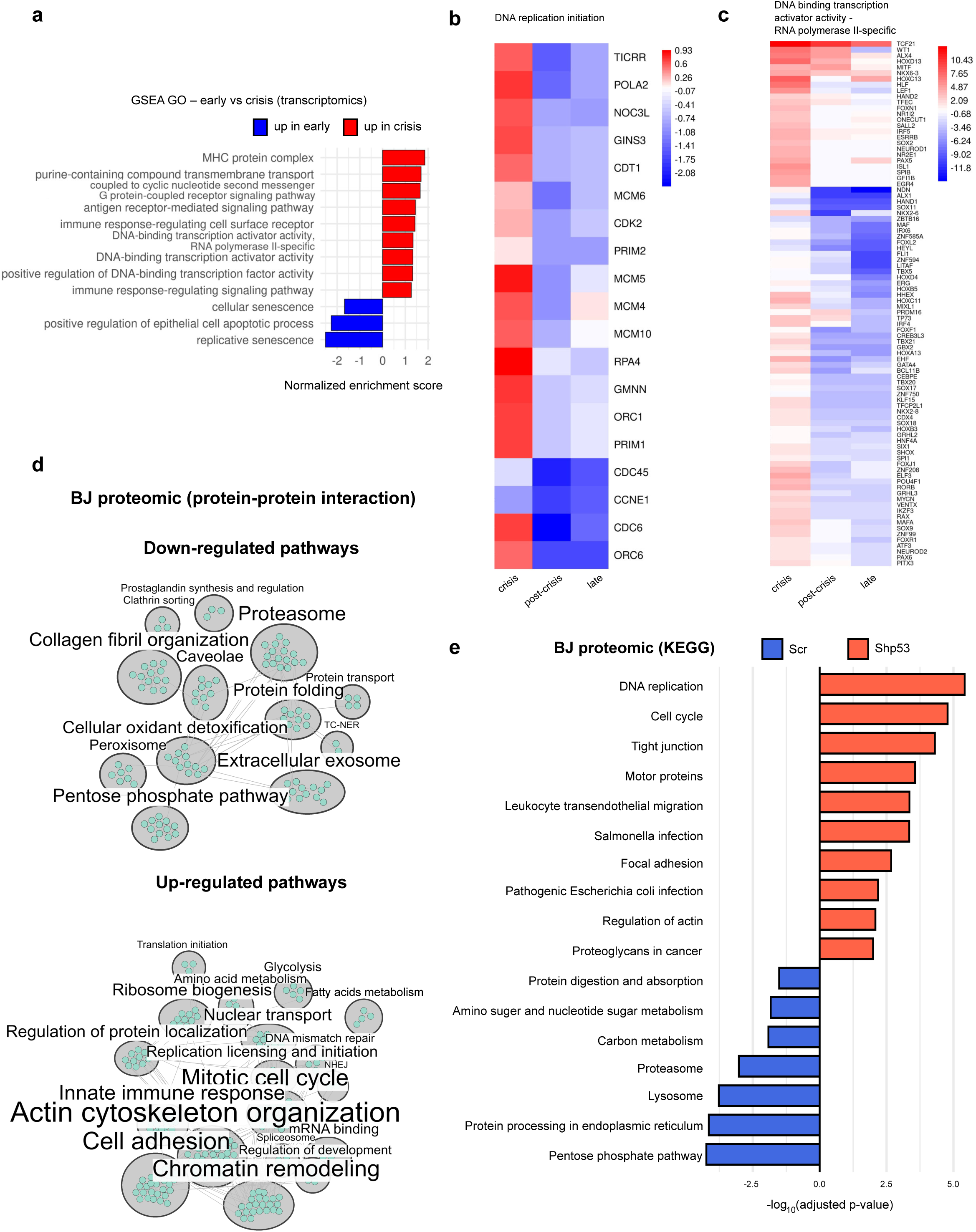
p53 loss dysregulates pathways related to nucleotide metabolism, translation, RNA processing and replication regulation. **a.** GSEA GO analysis of key significantly enriched processes in LFS041 fibroblasts, comparing early (p.19) and crisis (p.29) passages using bulk RNA sequencing. **b, c.** Differential gene expression analysis of DNA replication initiation **(b)** and DNA-binding transcription activator activity (RNA polymerase II-specific) **(c)** based on bulk RNAseq analysis in LFS041. Heatmaps display differentially expressed genes (DEGs) across early (p.19), post-crisis (p.65), and late (p.346) passages, with comparisons relative to early passage (p.19). The colour gradient represents the level of gene expression changes derived from NOIseq, with red indicating upregulation (in crisis, post-crisis, or late passages) and blue indicating downregulation as compared to early passage. **d.** Protein-protein interaction networks of significantly differentially expressed proteins (log2 fold change >1 or <-1, P-value <0.05) in BJ shp53 cells relative to control scr cells. Title text size corresponds to the number of proteins included in each sub-network. Downregulated pathways (**Upper panel**). Upregulated pathways (**Lower panel**). **e.** KEGG biochemical pathway enrichment of differentially expressed proteins following p53 downregulation. Enrichment was calculated by gProfiler using Fisher’s one-tailed test (P-value<0.05).

To gain a comprehensive molecular understanding of the consequences of p53 loss, we performed proteomics analyses in LFS cells and in a complementary cellular system, namely immortalized fibroblasts (BJ-hTERT, BJ) infected with a plasmid expressing a short hairpin against p53 (shp53). In the shp53 cells, p53 mRNA levels are 20-fold lower relative to control cells and no p53 protein can be detected by western blot (**Supplementary Information, Fig. S6f, g**). In BJ cells, proteomic profiling identified 556 differentially expressed proteins following p53 downregulation, including 323 upregulated and 233 downregulated proteins (Welch’s t-test, p<0.05). To interpret the functional impact of these changes, we performed protein-protein interaction (PPI) network analysis and functional enrichment analyses (cumulative hypergeometric test, g:SCS) using several databases including KEGG, GO, and REACTOME (Supplementary Information). PPI network analysis revealed distinct functional clusters among both up- and downregulated proteins; among downregulated proteins, a major cluster included the pentose phosphate pathway (PPP) – a critical supplier of ribose-5-phosphate and nicotine adenine dinucleotide phosphate (NADPH) required for nucleotide biosynthesis^36^. Specifically, seven PPP-related proteins (TALDO1, G6PD, PGD, PGM1, GPI, PFKM, and TKT) showed threefold reduction (**Fig. 4d**, top panel; supplementary information). Among upregulated proteins, PPI analysis revealed clusters involved in replication origin licensing, DNA repair, ribosome biogenesis, nucleoplasmic transport, and response to double-stranded RNA (dsRNA) – all of them are common markers of DNA replication stress and elevated transcriptional activity (**Fig. 4d**, bottom panel). Additional clusters included proteins involved in chromatin remodeling and actin cytoskeleton organization, indicating more general alterations in cellular structure and gene regulation. KEGG enrichment^37^ identified the PPP as the most significantly downregulated biochemical pathway (KEGG:00030, adjusted p-value=5.78x10^-5^; **Fig. 4e**), consistent with reduced nucleotide metabolism and statistically supporting the PPI results. GO^38,39^ and REACTOME^40^ enrichment further suggested activation of the ATR-dependent replication stress response (REAC:R-HSA-176187, adjusted p-value=3.95x10^-3^; **Supplementary Information**), enhanced DNA repair (GO:0006281, adjusted p-value=2.736x10^-5^; **Supplementary Information**) and replication initiation (GO:0006270, adjusted p-value=1.1x10^-3^; **Supplementary Information**), and reduced nucleotide biosynthesis (GO:0009117, adjusted p-value=0.014; **Supplementary Information**) following p53 downregulation. Based on the proteomic analysis in BJ cells, we revealed altered nucleotide metabolism upon p53 loss.

Importantly, these results were further supported by gene set enrichment and gene ontology (GSEA GO) analysis of proteome and RNA sequencing data performed on LFS cells at early, crisis and late passages (**Supplementary Information, Fig. S7a-e**). In particular, LFS cells during the crisis showed an increase in ribosome biogenesis and DNA duplex unwinding as compared to early-passage cells (**Supplementary Information, Fig. S7d-e**). Taken together, the proteome and transcriptome analyses in both BJ and LFS cells reveal that nucleotide metabolism is impaired following p53 loss, suggesting a role in the observed genomic instability.

### Mild replication stress in early-passage LFS cells is further enhanced in crisis cells

Classical replication stress, characterized by slowed fork progression and increased firing from dormant origins to compensate for the reduced replication rates, is a major cause of genome instability in precancerous and cancer cells^41–43^. As the proteomic analyses pointed to increased replication licensing and firing, as well as DNA repair in crisis and in shp53 cells, we further investigated whether replication stress is the molecular basis underlying genomic instability in LFS cells. We analyzed the DNA replication dynamics using DNA combing, a high-resolution technique enabling the detection of DNA replication dynamics in single DNA strands^44^. An example of a single combed DNA fiber is shown in **Fig 5a**. We quantified the replication fork rate progression and the fork distance, which represent dormant origin firing^44,45^. As compared to fibroblasts with WT p53, LFS041 cells showed slower replication rate in early-passage cells (1.1 kilobase per minutes (kb/min)), which was further significantly reduced in crisis LFS cells (0.8 kb/min, **Fig. 5b, left**). Fork distance was also significantly decreased in early-passage cells relative to normal fibroblasts and was further decreased in the crisis cells, indicating additional replication origins firing (**Fig. 5b, right**). The late-passage LFS cells that escaped from the crisis exhibited normalization of replication dynamics (**Fig. 5b**), consistent with the observed reduced DNA damage, chromatin bridges, multipolar spindles and micronuclei (**Fig. 1**). These results show that early-passage cells with heterozygous *TP53* already suffer from mild replication stress, which is further exacerbated in cells with a complete loss of WT p53. We further analyzed the replication dynamics and DNA damage in LFS087 cells, for which the earliest available passages for this analysis had already lost WT p53. The results showed a very slow replication rate and short fork distance as compared to normal fibroblasts (**Supplementary Information, Fig. S8a**) together with high DNA damage levels (**Supplementary Information, Fig. S8b**), further suggesting that total loss of WT p53 leads to replication stress and DNA damage.

**Figure 5.**
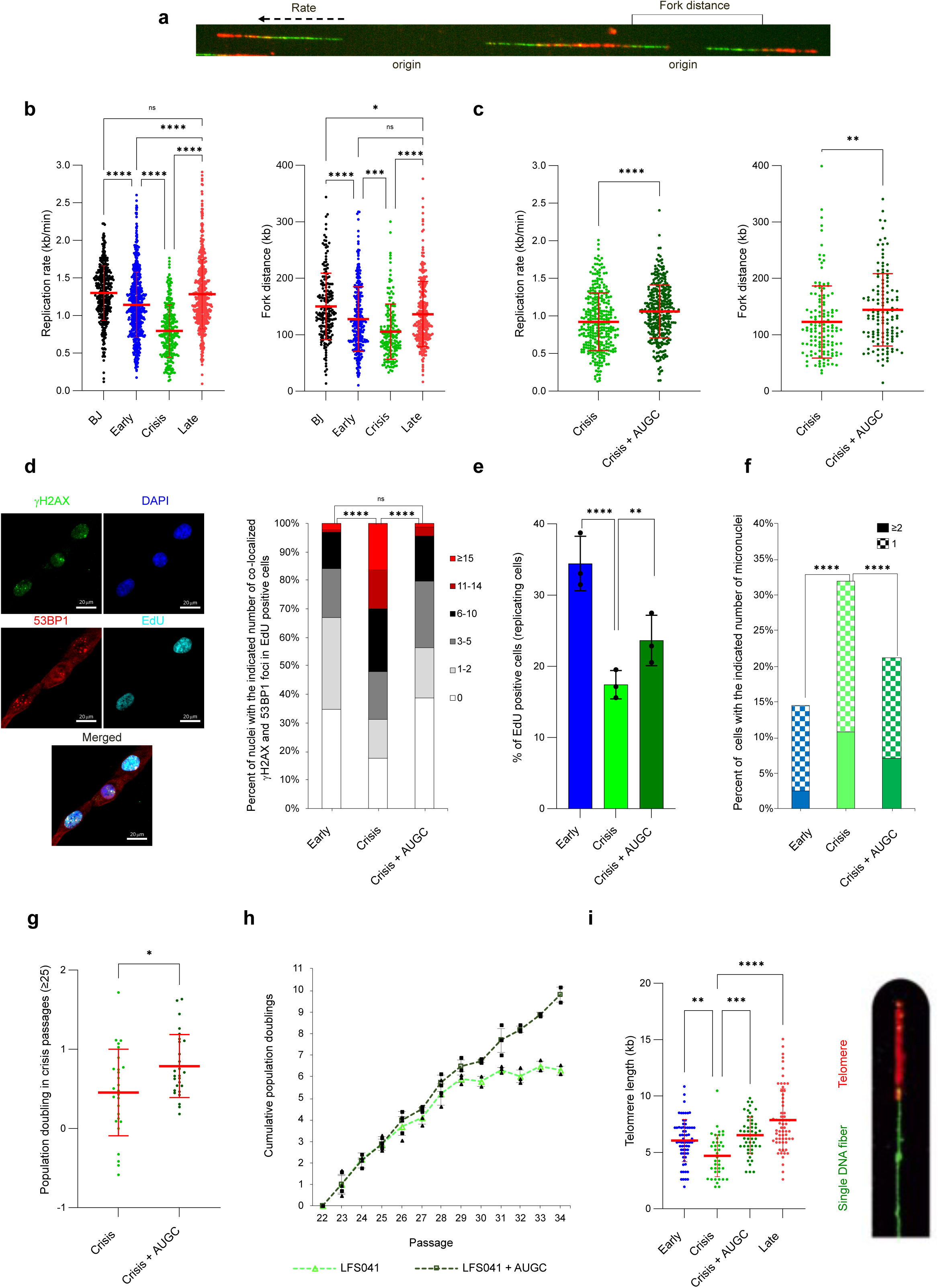
Replication stress due to insufficient nucleotides is the underlying mechanism leading to genomic instability and early growth crisis in LFS cells. **a.** Example of a single combed DNA fiber showing two adjacent out-going origins of replication with an in-going forks towards termination (within the middle of the red signal). **b. Left -** Fork rate (kb/min) distribution. Red lines indicating the mean±SD fork rate values in BJ control cells (1.3±0.4; n=400), early passage cells (1.1±0.4; n=624), crisis cells (0.8±0.4; n=255) and late passage cells (1.3±0.5; n=553). **Right -** Fork distance (kb) distribution. Red lines indicating the mean±SD fork distance values in BJ control cells (150±59; n=196), early passage cells (128±57; n=313), crisis cells (106±49; n=164) and late passage cells (137±58; n=276). The data are a summary of three independent experiments. Statistical analysis was performed using one-Way Anova test. **c. Left -** Fork rate (kb/min) distribution. Red lines indicating the mean±SD fork rate values in crisis cells (0.9±0.4; n=316), crisis cells supplemented with long-term nucleoside supply (1.1±0.4; n=341). **Right -** Fork distance (kb) distribution. Red lines indicating the mean fork distance values in crisis cells (123±64; n=133), crisis cells supplemented with long-term nucleoside supply (144±64; n=131). The data are representative of three independent experiments with similar results. Statistical analysis was performed using unpaired t-test. **d.** Co-localization of γH2AX (green) and 53BP1 (red) foci in EdU (cyan) positive cells for early passage cells (mean foci/cells 2.8; n=133), crisis cells (7.9; n=160) and crisis cells supplemented with long-term nucleoside supply (3; n=248). Representative images of co-localized γH2AX (green) and 53BP1 (red) foci in EdU positive nuclei **(left)**, and percentage of EdU positive cells with indicated number of co-localized γH2AX and 53BP1 foci/nucleus **(right)**. The data are representative of two independent experiments with similar results. Statistical analysis was performed using Mann Whitney rank-sum test done on the distribution. **e.** Percentage of EdU positive cells in early passage cells (mean 34%, n = 932), crisis cells (17%, n = 763), and crisis cells supplemented with long-term nucleoside supply (24%, n = 960). Bars represent the mean±SD percentage of EdU positive cells, and individual dots indicate the values from three independent experiments for each condition. A Chi-Square test of independence was performed on the three repeats combined to compare the proportion of EdU positive cells relative to the total number of cells between each two conditions. **f.** Percentage of cells with the indicated number of micronuclei in early passage cells (n=857), crisis cells (n=605) and crisis cells treated with long-term AUGC (n=829). Chi-Square test of Independence was performed to compare the distribution of cells with micronuclei between each condition. The data are a summary of three independent experiments. **g.** Population doubling in crisis passages for untreated cells (mean±SD 0.79±0.4; n=27) or treated cells with long term-nucleoside supply (0.45±0.55; n=27). **h.** Cumulative population doubling of untreated 041-cells from passage 22 till they reached the crisis, and of supplemented cells with long-term nucleoside supply. P-value calculated using t-test on the distribution of population doubling data from crisis passages (≥25) as shown in g. **i.** Telomere length distribution analyzed by Telosizer assay in early passage cells (mean 6.1 kb, n=68), crisis cells (4.7 kb, n=39), crisis cells treated with long term-nucleoside (6.6 kb, n=52) and late passage cells (7.9 kb, n=64). A representative image of a combed DNA fiber (green) and telomeric region detected by a FISH probe (red). Statistical analysis was performed using one-Way Anova test.*P<0.05, **P<0.01, ***P<0.001 and **** P<0.0001.

### Insufficient nucleotides lead to replication stress-induced genomic instability in crisis LFS cells

Several mechanisms were suggested to generate replication stress, including increased origin firing, transcription-replication conflicts, hypertranscription and nucleotide pool depletion^46–48^. The latter was shown to be a central mechanism underlying oncogene-induced replication stress^46,48,49^. Our proteomic analyses showed that total p53 loss dysregulated nucleotide metabolism and also downregulated the PPP (**Fig. 4**). In addition, our findings showed a significant increase in replication stress and genomic instability at the crisis cells compared to the early. This suggests that total p53 loss induced insufficient pool of nucleotides underlying enhanced replication stress and genomic instability. In order to test this possibility, we supplemented LFS041 with exogenous nucleosides (A,U,G and C) starting from pre-crisis passages. Nucleoside supply significantly increased the replication rate and fork distance in the supplemented crisis LFS041 cells, indicating a rescue of the replication stress (**Fig. 5c and Supplementary Information, Fig. S8c**). To investigate whether nucleoside insufficiency contributes to the DNA damage in crisis cells, we analyzed the effect of nucleoside supply on the DNA damage levels in S-phase cells, that reflects replication-induced DNA damage. For this, we performed immunofluorescence using antibodies against γH2AX and 53BP1 in LFS041 cells that incorporated EdU to label replicating cells (**Fig. 5d, left panel**). The early-passage replicating LFS041 cells already exhibited a high level of DNA damage (see **Fig. 5d, right panel**), which was further increased in the crisis of replicating cells. Exogenous nucleoside supply significantly decreased the DNA damage in the crisis replicating cells (**Fig. 5d, right panel**), and also significantly decreased the overall damage (**Supplementary Information, Fig. S8d)**. Notably, nucleoside supply increased the percentage of EdU-positive cells in crisis LFS cells (**Fig. 5e**), indicating that the nucleoside supply increased the percentage of proliferating cells. To directly investigate whether nucleotide insufficiency contributes to the emerging genomic instability, including chromothripsis, we quantified micronuclei formation (which is sufficient to induce genomic rearrangements^6^) under nucleoside supply. As expected, nucleoside supply significantly reduced the micronuclei formation (by 34%, p-value=1.1x10^-4^) in crisis LFS cells (**Fig. 5f**). We further studied the effects of nucleoside supply on replication dynamics, DNA damage and micronuclei in crisis cells from patient LFS087, in which WT p53 allele is lost (**Supplementary Information, Fig. S8e-g**). Nucleoside supply led to increased replication rate, reduced DNA damage and micronuclei formation, indicating that insufficient nucleotides is the basis for the replication stress and genomic instability in crisis LFS cells. Even though late-passage LFS cells completely lost the WT p53 protein (**Supplementary Information, Fig. S8h**), they rescued their replication stress-induced genomic instability (**Figure 1b** and **Figure 5a**), possibly via *MYC* amplification (**Fig. 2d**) leading to *MYC* overexpression, resulting in upregulation of its target genes involved in de-novo nucleotide biosynthesis (**Supplementary Information, Fig. S8i**).

### Insufficient nucleotide pool leads to growth crisis in LFS cells

As exogenous nucleoside supply increased the percentage of EdU-positive cells in crisis LFS cells (**Fig. 5e**), we hypothesized that insufficient nucleotide pool leads to the growth crisis. To test our hypothesis, we supplemented pre-crisis LFS041 cells with exogenous nucleosides and quantified cell proliferation. In the unsupplemented cells, the slowing proliferation started at passage 29, until cells reached a total growth crisis at passage 32 (**Fig. 5g, h**). Interestingly, cells supplemented with nucleosides continued to proliferate beyond passage 29 and for more than 3 additional population doublings (**Fig 5g, h, Supplementary Information, Fig. S8j**). To further investigate the possibility that nucleoside supply might prevent telomere shortening due to replication stress, which might be the cause of the growth crisis, we treated LFS041 cells with long-term exogenous nucleoside supply (starting from early passage) and quantified telomere length using the Telosizer assay^50^. Interestingly, nucleoside supply indeed prevented telomere shortening and crisis (**Fig. 5i**). In line with this, a recent study showed that human telomere length is controlled by thymidine nucleotide metabolism^51^. Altogether, we showed that nucleoside supply rescues the replication stress, leading to rescue of the DNA damage, micronuclei formation, telomere shortening and cell proliferation arrest. Our findings further indicate that nucleotide insufficiency plays a significant role in promoting genomically unstable cells, resulting in a growth crisis, from which chromothriptic cells emerge.

### Loss of p53 alone disrupts the nucleotide pool, leading to replication stress, DNA damage, and micronuclei formation

We further hypothesized that p53 downregulation by itself is sufficient to induce replication stress and genomic instability due to nucleotide insufficiency. Hence, we studied the effects of p53 downregulation and nucleoside supply on the replication dynamics using DNA combing. Analysis of the DNA replication dynamics revealed a significant decrease in the mean replication rate and replication fork distance in p53-downregulated BJ cells (**Fig. 6a, b**). Interestingly, exogenous nucleoside supply significantly increased the replication rate and fork distance (**Fig. 6a, b**), indicating rescue of the replication stress generated by p53 downregulation. In addition, p53 downregulation markedly increased DNA damage and micronuclei formation, both of which were rescued by nucleoside supplementation (**Fig 6c, e**). To further investigate the effects of p53 downregulation on the nucleotide pools, we performed untargeted metabolomic analysis using liquid chromatography-mass spectrometry (LC-MS) in BJ cells following p53 downregulation. Pathway enrichment and impact analyses (based on KEGG human metabolic pathways using the hypergeometric test) revealed PPP and the pyrimidine and purine metabolism pathways among the top downregulated pathways (**Fig. 6f** and **Supplementary Information, Fig. S9a, b**). The levels of metabolites such as Ribose 5-phosphate and all four nucleosides, Adenosine, Uridine, Guanosine and Cytosine, were significantly reduced following p53 downregulation (**Fig. 6g**). In addition, metabolomic analysis of LFS041 early-passage cells harboring heterozygous *TP53* showed a significant reduction in nucleotide metabolites compared to non-LFS fibroblasts (**Supplementary Information, Fig. S9c**). Taken together, our findings suggest that p53 loss limits the nucleotide pool available for DNA replication, leading to replication stress and genomic instability.

**Figure 6.**
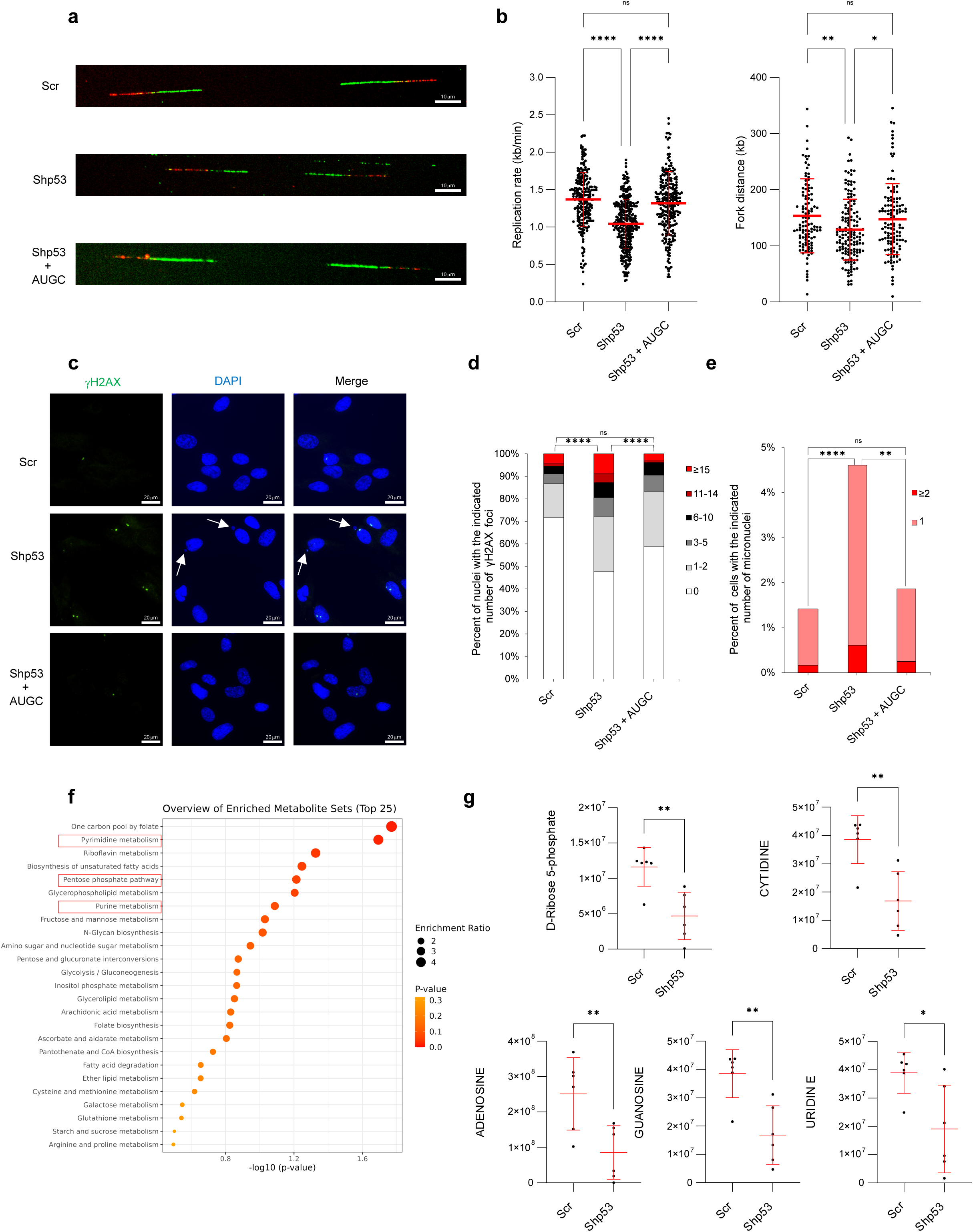
p53 loss by itself decreases the nucleotide pool leading to replication stress and genomic instability. **a.** Representatives combed DNA fibers from scr cells, shp53 cells and shp53 cells supplemented with AUGC for 48 hours. **b. Left -** Fork rate distribution (kb/min). Red lines indicating the mean±SD fork rate values in BJ cells (scr) (1.4±0.4; n=229), shp53 cells (1.0±0.3; n=308), shp53 cells supplemented with AUGC for 48 hours (1.3±0.4; n=252). **Right -** Fork distance (kb) distribution. Red lines indicating the mean±SD fork distance values in scr cells (154±66; n=111), shp53 cells (129±54; n=152), shp53 cells supplemented with AUGC for 48 hours (148±63; n=124). The data are a summary of three independent experiments. Statistical analysis was performed using one-Way Anova test. **c.** Representative images for γH2AX foci (green) in nuclei (blue) from scr cells, shp53 cells and shp53 cells supplemented with AUGC for 48 hours. White arrows point towards micronuclei. **d.** Percentage of nuclei with the indicated number of γH2AX foci for scr (mean foci/cells 1.9; n=1711), shp53 (4.1; n=1751) and shp53 + AUGC for 48 hours (1.9; n=1634) cells. The data are a summary of four independent experiments. Statistical analysis was performed using Mann Whitney rank-sum test done on the distribution. **e.** Percentage of cells with the indicated number of micronuclei for the same condition as shown in D. The data are a summary of three independent experiments. **f.** Pathway enrichment analysis based on the metabolite levels from LC-MS in scr and shp53 cells. The data include six biological repeats in each condition. Red boxes indicate important pathways related to nucleotide biosynthesis. **g.** Normalized metabolite levels of D-Ribose 5-phosphate and Cytidine, Adenosine, Guanosine and Uridine in scr and shp53 cells, measured using LC-MS based metabolomic analysis. Statistical analysis was performed using unpaired t-test. The data include six biological repeats in each condition.*P<0.05, **P<0.01, ***P<0.001 and **** P<0.0001.

### Loss of p53 induces hypertranscription, resulting in increased nucleotide consumption and genomic instability

In addition to DNA replication, transcription is a major source of nucleotide consumption, as RNA accounts for nearly 90% of total cellular nucleic acids^52^. Our proteomic data showed an increase of direct regulators of RNA transport and processing proteins (**Fig. 4d**) such as NCL, SSB, HNRNPD, HNRNPA3 and RNGTT. In addition, we found high ribosome biogenesis and increased response to dsRNA (including STING1, DDX58, EIF2AK2, OAS2 and OAS3). These results led us to hypothesize that p53 loss increased transcription and thus excessive nucleotide consumption, resulting in insufficient nucleotide availability for DNA replication. First, we analyzed nascent transcription levels following p53 loss in BJ cells, by quantifying EU incorporation into nascent RNA. The results showed that p53 loss indeed leads to hypertranscription (**Fig. 7a**). Next, we quantified the total RNA level in single cells, using acridine orange which emits red fluorescence upon binding to RNA. The results showed an increased fluorescent intensity in the shp53 cells, indicating a significant increase in the total cellular RNA levels (**Supplementary Information, Fig. S10a-c**). We then analyzed the possibility that this hypertranscription contributes to elevated levels of DNA damage induced by p53 downregulation. For this, we divided the EU intensity into tertiaries (high, mid and low) and quantified the DNA damage level in the same nuclei. In high-EU nuclei, the level of DNA damage was high, whereas in nuclei showing low EU, the level of damage was very low (**Fig. 7b**). We further analyzed the transcription levels in LFS041 compared to non-LFS primary fibroblasts as control. In early-passage LFS041 cells, with heterozygous p53 variant, an increased transcription relative to non-LFS cells was detected. The transcription was further increased in the crisis LFS cells, in which there is a total loss of functional p53 (**Supplementary Information, Fig. S5a-e).** Late-passage LFS041 cells exhibited high EU intensity compared to non-LFS fibroblasts, further supporting a link between p53 loss and hypertranscription (**Supplementary Information, Fig. S10d, e**). An association between increased RNA transcription and aneuploidy was recently shown^53^. Therefore, we assessed the aneuploidy status of p53-downregulated cells using low-pass whole-genome sequencing (lp-WGS) and found no detectable changes (**Supplementary Information, Fig. S10f**), suggesting that the hypertranscription induced by short-term p53 loss occurs independently of aneuploidy. Altogether, these results indicate that hypertranscription is a direct consequence of p53 loss and contributes to the accumulation of DNA damage.

**Figure 7.**
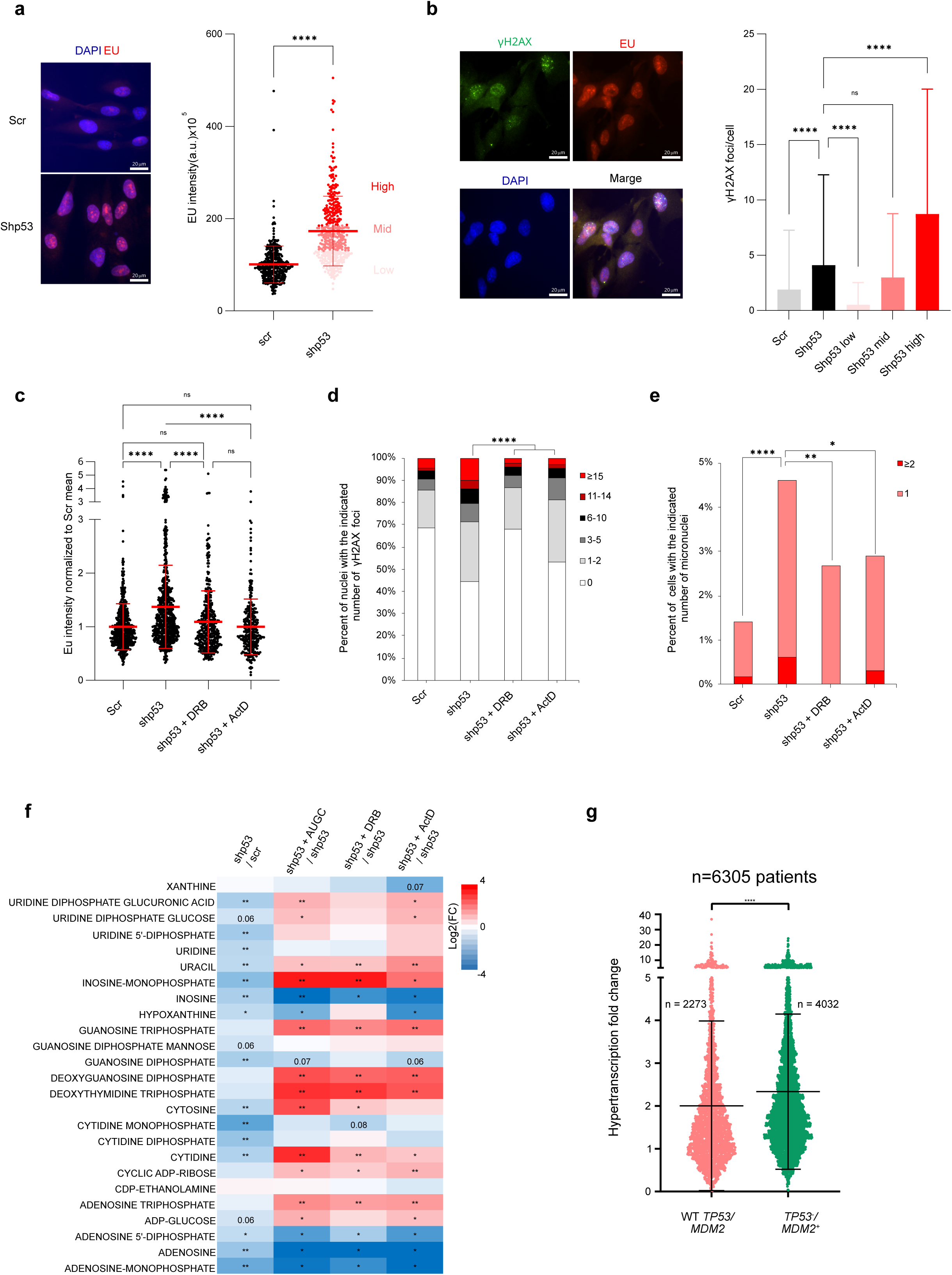
p53 loss by itself increases transcription levels resulting in nucleotide consumption and genomic instability. **a. Left -** Representative images of EU (red) and DAPI (Blue) staining, in scr and shp53 cells. Right - quantification of nuclear EU intensity per cell in Scr (101±40; n=383) and shp53 cells (173±76; n=398). EU distribution is divided to tertiaries. The data are representative of three independent experiments with similar results. Statistical analysis was performed using the student’s t-test. **b. Left -** Representative images of γH2AX (green), EU (red), nuclei (blue) in the same nucleus. **Right –** Quantification of γH2AX foci in scr (mean foci/cells 1.9; n=383), shp53 (4.1; n=398) and in each tertial of shp53 distribution as shown in A. The mean foci per nucleus in the low (0.5; n=132), mid (3.0; n=132), high (8.7; n=133) tertial. **c.** Transcription levels quantified using nuclear EU intensity and normalized to scr mean for scr cells (1±0.3; n=708), shp53 cells (1.37±0.3; n=731) and shp53 cells treated with 10-15 μM DRB for 48 hours (1.1±0.4; n=392) or treated with 10ng/ml ActD for 3 hours (1±0.1; n=304). The data are representative of two independent experiments with similar results. Statistical analysis was performed using one-Way Anova test. **d.** Percentage of nuclei with the indicated number of γH2AX foci for Scr (mean foci/cells 1.9; n=791), shp53 (4.4; n=831), shp53 + 10-15 μM DRB for 48 hours (1.6; n=696) and shp53 + 10ng/ml ActD for 3 hours (1.9; n=769). The data are a summary of two independent experiments. Statistical analysis was performed using Mann Whitney rank-sum test done on the distribution. **e.** Percentage of cells with the indicated number of micronuclei with the same conditions as d, number of analyzed cells per condition; Scr (n=1126), shp53 (n=1076), shp53 + 10-15 μM DRB for 48 hours (n=945) and shp53 + 10ng/ml ActD for 3 hours (n=934). The Chi-Square test of Independence was performed to compare the distribution of cells with micronuclei between each condition. **f.** Heatmap representation of log fold changes (log FC) in metabolite levels analyzed using LC-MC in scr, shp53 and shp53 cells treated with 50 μM AUGC for 48 hours, 15 μM DRB for 48 hours or 10ng/ml ActD for 3 hours. The comparisons are shown in the columns (appear at the top of the panel) and include six biological repeats for each condition. In all comparisons the same shp53 data was used. Green - increased metabolite levels compared to the indicated condition. Red - decreased metabolite levels compared to the indicated condition. P-value for each comparison calculated using one-sided t-test and indicated in each box (*p < 0.05, **p < 0.01, the value <0.1 is indicated). **g.** Hypertranscription fold-change (re-analysis of the TCGA cohort from the study by Zatzman et al.) shown for two groups: WT *TP53/MDM2* (mean ± SEM: 2 ± 0.04, n=2273) and *TP53-/MDM2+* (mean ± SEM: 2.33 ± 0.03, n=4032). Each circle represents a tumor sample, with the total number of samples per group indicated on the plot. Data are presented as mean ± SEM. Statistical significance was assessed using a non-parametric t-test (Mann-Whitney test) to compare two independent groups, as the data were not normally distributed. P-values less than 0.05 were considered statistically significant (*p < 0.05, **p < 0.01, ***p < 0.001, **p < 0.0001).

To investigate whether hypertranscription is the source of DNA damage following p53 downregulation, we reduced the nascent transcription levels to those observed in control using either the specific RNA Pol II inhibitor, DRB or the RNA Pol I inhibitor, Actinomycin D (for 3hrs, 10ng/ml that inhibits only the RNA Pol I) (**Fig. 7c**). The results showed a significant reduction in DNA damage and micronuclei formation, in cells treated with each of the transcription inhibitors (**Fig. 7d, e**). To further investigate the effect of nucleoside supply or transcription inhibitors on the nucleotide metabolites, we performed LC-MS in p53-downregulated BJ cells treated with AUGC for 48 hours, DRB for 48 hours or ActD for 3 hours. The analysis included 23 nucleotide metabolites that were identified in all conditions. Our results showed a significant increase in more than 50% of the analyzed metabolites following AUGC supply and in at least one of the transcription inhibitor treatments **(Fig. 7f)**. Furthermore, to investigate potential links between p53 dysfunction and hypertranscription in cancer, we compared hypertranscription scores calculated from RNA sequencing analyses between cancers with and without p53 alterations (**Fig. 7g**)^54^. Importantly, p53 inactivation was associated with higher hypertranscription scores both in pan-cancer context (**Fig. 7g**) and tumor entity specific analyses (**Supplementary Information, Fig. S10g-n**). The effect size observed for p53 inactivation was comparable to hypertranscription associated with amplifications of *MYC*, which is a known driver of hypertranscription^55,56^, and the difference in hypertranscription between p53-proficient and deficient tumors was significant irrespective of the *MYC* status (**Supplementary Information, Fig. S10o-p).** Taken together, our results indicate that high transcription driven by p53 downregulation leads to excessive nucleotide consumption, resulting in replication stress leading to genomic instability including complex rearrangements (**Fig. 8**).

**Figure 8.**
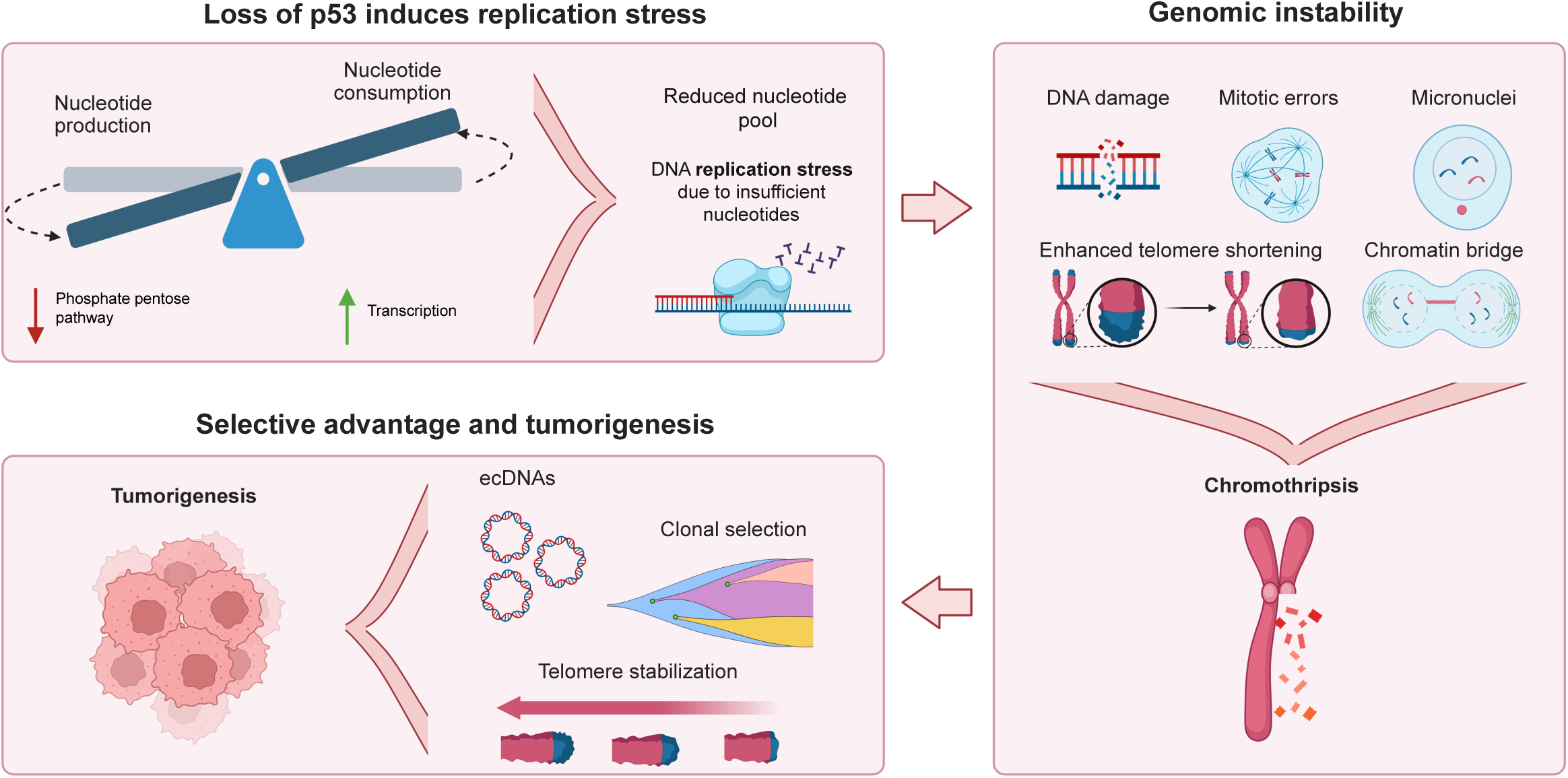
Schematic illustration of the effect of p53 loss leading to impaired nucleotide metabolism, replication stress and chromothripsis. Loss of WT p53 reduces the nucleotide pool by impairing the nucleotide regulation: increasing nucleotide consumption (due to hypertranscription) and decreasing nucleotide production (due to downregulation of the phosphate pentose pathway). This results in replication stress due to insufficient nucleotides, which in turn causes DNA damage, mitotic errors, enhanced telomere shortening and micronuclei formation. This leads to genomic instability, including chromothripsis. After chromothripsis, telomere stabilization and ecDNA amplification play significant roles in clonal selection, driving cancer development.

## DISCUSSION

In the current study, we conducted longitudinal analyses to understand the mechanistic role of WT p53 loss as a driver of genomic instability. We revealed that the loss of WT p53 function drives genomic instability by increasing transcription activity and hence nucleotide consumption, resulting in replication stress due to insufficient nucleotide pools for DNA replication. We also showed that the replication stress is sufficient to generate micronuclei and ecDNAs that fuel clonal selection (**Fig. 8**).

Transcription can pose a significant direct obstacle to the DNA replication process by either head-on collisions or co-directional conflicts^57^. Previous studies have shown that p53 regulates DNA replication by preventing transcription-replication collisions^24^. Inhibition of transcription by treatment with DRB, an RNA pol II transcription inhibitor, rescued the replication stress, through reduced conflicts between the replication and transcription machineries^24^. In our study, we revealed a novel indirect mechanism, by which transcription challenges the DNA replication. We demonstrated that p53 loss significantly elevates transcription levels (**Fig. 7a**), and since transcription is a major nucleotide-consuming process⁵¹, this increase leads to nucleotide depletion (**Fig. 6f, g** and **Supplementary Information, Fig. S9a-d**), promoting replication stress. In line with this, exogenous supply of AUGC effectively rescued the replication stress induced by p53 loss (**Fig. 6a-e**) while transcription normalization by RNA pol I or RNA pol II inhibitors (actinomycin D and DRB, respectively) (**Fig. 7c-e**) reduced p53-induced genomic instability and significantly increased several nucleotide metabolites (**Fig. 7f**). The fact that replication stress can be rescued by either inhibiting mRNA or rRNA transcription, or by supplementing nucleotides, strongly supports the conclusion that nucleotide insufficiency is the underlying cause of replication stress promoting R-loop formation and elevated transcription-replication collisions. The hypertranscription phenotype following p53 loss identified in our study together with the increased response to double-stranded RNA (**Fig. 4d**) are consistent with previous work showing that p53 loss leads to elevated immune response due to increased formation of double stranded RNA, defined as Transcription of Repeats Activates Interferon^58^. In addition, a recent study demonstrated that p53 loss promotes derepression of heterochromatin, with a massive loss of histone H3K9me3, resulting in increased transcription of microsatellites, leading to R-loop formation and genomic instability at those sites^59^. Altogether, these data support our hypothesis that global hypertranscription is a consequence of WT p53 loss.

LFS fibroblasts offer a complex model for p53 loss, harboring numerous copy-number alterations (**Fig. 2a**), high levels of micronuclei (**Fig. 1h**) and telomere shortening (**Fig. 1c-d** and **Fig. 5i**). Thus, we also studied the direct effects of p53 loss by utilizing the BJ cell system, a controlled model that allows p53 downregulation without introducing additional perturbations. Although BJ cells offer a model for p53 loss, rather than a p53 dysfunction as in LFS, both cell systems showed broadly consistent results. We demonstrated that p53 loss is sufficient to induce replication stress and DNA damage through nucleotide insufficiency, supported by the finding that nucleoside supplementation rescues replication stress, DNA damage, and micronuclei formation (**Fig. 6a-e**). Furthermore, we revealed that hypertranscription is a direct consequence of p53 loss and a major source of excessive nucleotide consumption, resulting in perturbed replication, as normalizing transcription raised nucleotide pools and rescued the DNA damage. Taken together, the p53 downregulation system provided direct evidence that p53 loss reduces nucleotide availability by increasing nucleotide consumption through hypertranscription.

It is important to note that the LFS system is highly relevant to cancer biology due to its spontaneous progression toward fast proliferating, spontaneously immortalized cells, paving the way to a deeper understanding of early cancer biology. LFS is a classical example for p53 haploinsufficiency^20–22^. Intriguingly, early-passage fibroblasts from LFS041, carrying a frameshift germline variant that has no effect on the second (WT) p53 allele (non-dominant negative mutation), suffer from replication stress, DNA damage and various forms of genomic instability (**Fig. 1, 2** and **5**). This raises the possibility that in cases of p53 heterozygosity, the replication is already perturbed, predisposing to DNA damage, which can result in genomic alterations and loss of heterozygosity. Our work shows that one copy of WT p53 is not sufficient to prevent cancer, as the cells still lose their proliferation barrier and suffer from replication stress and genomic instability. Furthermore, our findings demonstrate that insufficient nucleotide pools drive the spontaneous growth crisis in LFS cells by enhancing replication stress, telomere shortening, micronuclei formation and genomic instability. Hence, we link replication stress with complex rearrangements, including chromothripsis, in LFS cells. Notably, exogenous nucleoside supply not only delayed the growth crisis, but also prevented telomere shortening. This highlights the critical role of nucleotide availability in maintaining telomere length, essential for sustained cell proliferation. Nucleoside supply further rescued the replication stress-associated DNA damage and micronuclei formation (**Fig. 5d-f**). Altogether, these results emphasize the direct link between nucleotide insufficiency and replication stress, telomere instability and low proliferation capacity, underscoring the importance of nucleotide availability in preserving genome stability and restraining cancer development. Furthermore, LFS cell progression from early and crisis to late immortalized stages recapitulates cancer development from precancerous stages, enabling the study of early cancer hallmarks. A previous work analyzing precancerous cells revealed that from its earliest stages, cancer development is associated with DNA damage and fragile site instability, suggesting the existence of DNA replication stress already in precancerous stages^60^. Here, by directly analyzing the replication dynamics, we detected replication stress and DNA damage already in early-passage LFS cells, supporting that already precursor cells suffer from perturbed replication dynamics and genomic instability. Under such conditions, cells with advantageous adaptations may emerge, favoring the selection of specific clonal subpopulations. Indeed, we found in LFS041 cells that late-passage cells not only rescue the replication stress and DNA damage, but also exhibit traits such as clonal dominance, increased proliferative capacity and survival. We showed that the late-passage cells in LFS041 overcome the chronic replication stress possibly by upregulation of nucleotide biosynthesis through amplification of the oncogene c-*MYC*, which is a master regulator of many nucleotide biosynthesis genes (**Supplementary Information, Fig. S8g, i**). Ultimately, our findings underscore the role of replication stress in shaping clonal evolution and might provide insights into earliest cancer features.

The BJ fibroblast system models the direct acute effects of p53 loss, whereas the LFS system models its long-term chronic effects. Importantly, the proteomic data from both cellular systems revealed key shared pathways highlighting the contribution of p53 deficiency, including upregulation of the ribosome biogenesis pathways (translation) and inflammation and DNA repair pathways and dysregulation of the nucleotide metabolism (**Fig. 4, Supplementary Fig. S7c-e**). The high translation and immune activation pathways, in both cellular systems, likely result from the observed hypertranscription. The proteomic data also showed key differences between the systems that highlight the acute and chronic effects of replication stress and genomic instability due to p53 loss. In BJ cells, there is a downregulation of the PPP metabolism, and upregulation of replication origin proteins and dsRNA (**Fig. 4d and e**). In LFS cells, we observed dysregulation of apoptotic processes, antigen processing and presentation, and metabolic processes, reflecting the chronic effects of p53 loss on the LFS cells (**Fig. 4a-c, Fig. S7c-e**).

Our analysis of the clonal evolution of fibroblasts upon p53 loss indicates a strong selection effect on structural variants, with only a minority of the rearrangements detected at early passages being still present at late passages (**Fig. 2c**). Of note, we focused primarily on the identification of genetic clones and did not study the potential role of transcriptional plasticity here. Even though fibroblasts that bypass the crisis show an altered morphology and selective advantages characteristic of malignant cells, including a high proliferation rate and activation of telomere lengthening mechanisms, this model system does not reflect the appearance of *bona fide* fibrosarcoma cells. Importantly, we cannot tell which genomic rearrangements might have already occurred in the patients in a small fraction of cells before taking the skin biopsy. However, in particular through comparisons with human tumors (**Fig. 7g**), we showed the relevance of this spontaneous model, and identified cellular and molecular changes in culture that lead to the crisis and to the escape from this crisis, which might act as a selection pressure. The impaired p53 function plays a key role in bypassing the crisis, as normal (p53 proficient) fibroblasts do not escape from the crisis but go into senescence. We also identified recurrent events between patients and between cells (e.g. chromothripsis on chromosome 15, telomere stabilization), supporting positive selection processes.

We and others showed that chromothripsis generates highly unstable derivative chromosomes, which evolve and fuel the acquisition of oncogenic alterations, clonal diversification, and intra-tumor heterogeneity^61,62^. Only a minority of chromothriptic events lead to dominant clones, as shown by our longitudinal study in LFS fibroblasts. As compared to bulk sequencing-based approaches, our single-cell resolution analyses allowed the detection of rare events and the precise characterization of the clonal evolution. The high diversity of rearrangements at early stages suggests that the crisis may act as a selection bottleneck, which favors the outgrowth of clones that carry a selective advantage such as *TERT* and *MYC* amplifications, potentially through ecDNAs generated by chromothripsis.

We previously demonstrated that aberrant activation of the Rb-E2F pathway, through overexpression of the HPV16 E6/E7 or cyclin E oncogenes, induces replication stress by promoting premature S-phase entry without a corresponding upregulation of nucleotide biosynthesis genes, leading to insufficient nucleotide supply^49^. Subsequently, several other studies also showed that nucleotide insufficiency is underlying oncogene-induced replication stress^46,48^. The results presented here, showing that loss of p53 affects nucleotide availability, further support the conclusion that nucleotide insufficiency is a central mechanism driving replication stress and genomic instability in cancer.

Finally, this study revealed a novel mechanistic role of hypertranscrption inducing nucleotide insufficiency in driving genomic instability and chromothripsis evolution, which may pave the way to novel translational opportunities to regulate nucleotide metabolism in the context of p53 deficiency. These may include: i) supply of nucleotides or inhibition of hypertranscription to prevent replication-induced genomic instability and by this limit cancer progression, in combination with other therapeutic approaches; ii) inhibition of nucleotide biosynthesis to prevent precancerous cell progression and selectively kill (or eliminate) cancer cells, as they may be more vulnerable to reduced nucleotide levels; iii) as hypertranscription is prevalent in human cancers with p53 dysfunction (**Fig. 7g**) and it is a predictive marker for response to immune checkpoint blockade^54^, the link between hypertranscription and p53 loss found in this study suggests putative translational opportunities to be evaluated.

## ONLINE METHODS

### Cell culture

#### LFS fibroblasts

Skin-derived fibroblasts from Li Fraumeni Syndrome patients (cells were kindly provided by Michael A. Tainsky) were grown in Minimum Essential Medium (MEM) Eagle (Sigma; M5650), supplemented with 10% fetal calf serum, 1% glutamine and 1% penicillin/streptomycin and cultured in 60 mm cell culture dishes (Falcon, #353004) at 37°C with 5% CO2. Cells from two LFS patients were used, namely patients LFS041 and LFS087. On average, early-passage cells were passaged once a week using 0.025% Trypsin/EDTA for trypsinization, post-crisis and late-passage cells twice a week, while crisis-passage cells took several weeks per passage to reach confluency. Fibroblasts from patients LFS8 and LFS9 (healthy skin-derived fibroblasts from the same family as LFS041) were grown in MEM alpha, supplemented with 15% fetal calf serum, 1% glutamine and 1% penicillin/streptomycin. STR profiling was conducted to authenticate the cells using a multiplex PCR-based method provided by Multiplexion. The analysis confirmed the cultures’ uniqueness and the absence of cross-contamination with other cell lines. Furthermore, the cells were confirmed to be mycoplasma-free using a PCR-based assay provided by Eurofins. Cell morphology was imaged using a Zeiss Axio Vert.A1 inverted microscope at 5X magnification. ZEN 3.0 Microscopy Software was used for image processing.

#### BJ fibroblasts

BJ-hTERT normal human fibroblast cells were grown in Dulbecoo’s modified Eagle’s medium (DMEM) supplemented with 10% Fetal bovine serum (FBS), 100,000 U/l penicillin and 100 mg/l streptomycin. The nucleosides supplemented media was prepared by adding 50µM of each nucleoside A, U, C and G from sigma. For p53 downregulation experiment, Bj-hTERT cells were infected by vectors expressing short-hairpin RNA against p53 mRNA as follows: Phoenix retroviral packaging cells were transiently transfected with plasmids, then the BJ-hTERT cells were infected three times with Phoenix cell supernatant containing defective viruses and 10 mg/ml polybrene. Following the infection, the cells were selected using selection reagents Puromycin for up to 7-10 days. The vectors are pRS-shp53-puro (p53 downregulation) and pRS-shRNA-puro (control), kindly received from Professor R. Agami.

### Growth curves for LFS fibroblasts

The time required to reach confluency was used as a proxy for the growth characteristics. IIn the growth curve graphs, the splitting ratio used at each passage is indicated for the corresponding passage number. A shorter time to reach confluency at specific passages suggests increased growth rates, while longer times indicate slower growth, reflecting changes in cell population dynamics. Graphs were generated using Microsoft Excel.

### Population doubling

Primary LFS cells grow in cell culture conditions as mentioned. The initial number of the cells were counted and determined, after several days the cells were trypsinized, counted and passaged by 1:2 ratio. The population doubling (PDL) for each passage calculated as PDL = 3.32 (log (Final cell number) - log (Initial cell number)). The sum of PDLs gives the cumulative population curve.

### Western blot analysis

Sample buffer was used for protein extraction. The extracted protein is loaded in 12% polyacrylamide gel, separated in 100mVolt for 50 min. The gel was transferred to a nitrocellulose membrane and blocked with 8% milk. The antibody hybridization and chemiluminescence (ECL) were performed according to standard procedures. The primary antibodies used are DO-1 (1:5000) from sigma for p53 detection, and mouse anti β-catenin (1:2,500) from BD-Biosciences. HRP-conjugated anti-mouse and anti rabbit was obtained from Jackson ImmunoResearch Laboratories (711-035-152, 1:5000).

### Quantification of micronuclei and chromatin bridges

Cells were seeded on coverslips in 6-well plates and fixed with 4% PFA at pH 6.8 for 15 minutes. Following fixation, the cells were washed twice with 1X PBS for 5 minutes, rinsed in double distilled water and then in absolute ethanol. The coverslips were air-dried at room temperature before being mounted on slides (Thermo Fisher Scientific, #J1800AMNZ) using DAPI Fluoromount-G Mounting Medium (Southern Biotechnology, #0100-20) and left to solidify in the dark at room temperature for 1 hour. Cell imaging and quantifications were performed using Zeiss AXIO imager 2. ZEN 2.3 Microscopy Software was used for data acquisition, while ZEN 3.0 Microscopy Software and ImageJ 1.54f were used for image processing. Micronuclei and chromatin bridges from 750 cells from at least three biological replicates per passage were quantified. GraphPad Prism 8 software was used to create graphs, perform statistical tests and calculate p-values. Statistical significance was assessed using a one-way ANOVA followed by Tukey’s multiple comparisons test, with p-values below 0.05 considered as statistically significant (*p < 0.05, **p < 0.01, ***p < 0.001, ****p < 0.0001).

### Immunofluorescence analysis of Phospho Histone 3 (pH3) and Acetyl-alpha-Tubulin and quantification of mitotic defects

Cells were seeded on coverslips in 6-well plates and allowed to reach 60-70% confluency. Once confluency was achieved, the coverslips were rinsed with cell wash PBS (1X PBS, 1% of 1M MgCl2) and the cells were fixed with 4% PFA at pH 6.8 for 15 minutes. Following fixation, the cells were washed twice with 50mM NH4Cl for 5 minutes each, and twice with 1X PBS with rocking. Next, a freshly prepared blocking buffer (1X PBS, 5% normal goat serum, 0.3% Triton X-100) was added to the coverslips and incubated for 1 hour at room temperature. After removing the blocking buffer, a mixture of primary antibodies – Phospho-Histone H3 (Ser10) (6G3) mouse mAb (Cell Signaling, #9706, Lot#10) diluted at 1:400 and Acetyl-α-Tubulin (Lys40) (D20G3) XP rabbit mAb (Cell Signaling, #5335, Lot #5) diluted at 1:800 – was added to each coverslip in an antibody dilution buffer (1X PBS, 1% BSA, 0.3% Triton X-100). The coverslips were then incubated overnight at 4°C. The next day, the coverslips were washed once with 1X PBS for 45 minutes and twice for 15 minutes. A secondary antibody mix consisting of goat anti-mouse and goat anti-rabbit antibodies, both diluted at 1:1000 in antibody dilution buffer, was applied to each coverslip and incubated in the dark for 90 minutes at room temperature. Subsequently, the coverslips were washed three times for 15 minutes in 1X PBS, rinsed in ddH2O, and then in absolute ethanol. Finally, the coverslips were air-dried at room temperature before being mounted on slides using DAPI Fluoromount-G Mounting Medium and left to solidify in the dark at room temperature for 1 hour. Cell imaging and quantification were performed using a Zeiss AXIO Imager 2 with ZEN 2.3 Microscopy Software. Representative images were captured using a Leica SP8 confocal microscope. Maximum intensity projections were applied using LAS X software, and further image processing was performed with both LAS X software and ImageJ 1.54f. A total of 100 cells per patient and per biological replicate across three independent replicates were quantified. GraphPad Prism 8 software was utilized to create graphs, perform statistical tests, and calculate p-values. Statistical significance was assessed using repeated-measures two-way ANOVA followed by uncorrected Fisher’s LSD for multiple comparisons; p-values below 0.05 were considered statistically significant (n = 3 per group; *p < 0.05, **p < 0.01, ***p < 0.001, ****p < 0.0001).

### Immunofluorescence staining for DNA damage markers

Cells were fixed in 3.7% formaldehyde/PBS for 10 min, permeabilized with 0.5% Triton/PBS for 10 min, and blocked with 3% BSA/PBS for 1 hour. The primary antibodies used were mouse anti-phosphorylated H2AX (Milipore, 1:100), rabbit anti-53BP1 (Bethyl Laboratories, 1:100) and incubated overnight in 4c. The secondary antibodies used were anti-mouse Alexa Fluor 488 (Invitrogen), anti-rabbit Alexa fluor 546 (Abcam). DNA was counterstained with a mounting medium for fluorescence with DAPI (Vector Laboratories). Images were analyzed for foci and micronuclei formation double blindly using Fiji^63^.

### *TERT*-FISH

50,000 cells in supplemented MEM were seeded on a UV-sterilized slide and incubated at 37°C with 5% CO2. Once the cells reached 60-80% confluency, the slides were rinsed with cell wash PBS and fixed with 4% PFA at pH 6.8 for 15 minutes. The slides were then rinsed twice with 1X PBS and dehydrated through a series of ethanol solutions: 70% ethanol for 2 minutes, 90% ethanol for 2 minutes, and 100% ethanol for 4 minutes. Slides were left to air dry at room temperature and subsequently stored at -20°C for short-term storage and at -80°C for long-term storage. Next, the slides were dehydrated again through an ascending ethanol series (70%, 90%, and 100% ethanol, each for 5 minutes at room temperature). Samples were digested in a pre-warmed digestion solution (0.01 M HCl, 0.02 mg/ml pepsin) for 10 minutes at 37°C in a water bath. The samples were then washed in 1X PBS for 5 minutes and post-fixed with 1% PFA on ice for 5 minutes. After washing in 1X PBS for 10 minutes, the slides were dehydrated through an ascending ethanol series and air-dried. Meanwhile, the DNA probe was prepared by mixing 10 µl of DNA *TERT* probe (Clone RP11-117B23, Source BioScience), which was labeled with Digoxigenin-11-dUTP (Roche, #1570013), Cot1-DNA (Life technologies, #15279-011), Herring Sperm DNA (Thermo Fisher Scientific, #15634017) and 1/20th volume ratio of 3 M sodium acetate (NaAc) with 2.5X the volume of 100% ethanol. This mixture was precipitated at -80°C for 30 minutes, centrifuged, and washed with 70% ethanol before air drying at 37°C. 10 µl of deionized formamide were added to the pellet and mixed well by shaking at 1000 rpm for 15 minutes, followed by adding 10 µl of hybridization mix per slide (20% dextran sulfate in 4X SSC pH 7.0) and shaken for at least 15 minutes before further processing. After air drying the slides, the probe was applied onto the slides, covered with a coverslip, and sealed with Fixogum. The slides were denatured on a heating block at 78°C for 10 minutes and incubated overnight in a humidified chamber at 37°C. The next day, the slides were washed with a freshly prepared Wash A solution (50% formamide in 2X SSC pH 7.0) in a water bath at 42°C with shaking for 10 minutes to remove coverslips, followed by three additional 5-minute washes. Next, the slides were washed with Wash B solution (0.5X SSC, pH 7.0, prewarmed at 60°C) at 42°C for three 5-minute washes, with gentle shaking. Blocking involved adding blocking buffer (200 µl of 4X SSC containing 3% BSA, pH 7.3) to the slide, covering them with parafilm, and incubating in a humidified chamber at 37°C for 30 minutes. For detection, 2.8 µl Anti-Digoxigenin-Rhodamine (ROCHE, #11207750910, 400 µg/ml stock concentration) in 200 µl detection buffer (blocking buffer diluted in 1:3 ratio) were added to the slides, which were then covered with parafilm and incubated in the humidified chamber at 37°C for 30 minutes, followed by incubation and washing with Wash C solution (0.1% Tween 20 in 4X SSC, pH 7.3) for three 5-minute washes at 42°C while shaking in the dark. Finally, slides were mounted using VECTASHIELD Vibrance Antifade Mounting Medium with DAPI (Vector Laboratories, #H-1800). Cell imaging and quantifications were performed using Zeiss AXIO imager 2 and ZEN 2.3 Microscopy Software for data acquisition, while ZEN 3.0 Microscopy Software and ImageJ 1.54f were used for image processing. 100 cells from each passage were quantified. GraphPad Prism 8 software was used to create graphs, perform statistical tests, and calculate p-values. Statistical significance was assessed using a nonparametric Kruskal-Wallis test, followed by Dunn’s multiple comparisons test; p-values less below 0.05 were considered statistically significant (*p < 0.05, **p < 0.01, ***p < 0.001, ****p < 0.0001).

### ALT FISH

The ALT-FISH protocol was implemented as previously described^64^, with slight modifications outlined below. A total of 50,000 cells in supplemented MEM were seeded onto UV-sterilized slides and incubated at 37°C with 5% CO2. Later, slides were washed twice with PBS, followed by fixation with 70% ice-cold ethanol for 20 minutes at room temperature. Next, the slides were washed twice with a washing buffer (100 mM Tris–HCl pH 8, 150 mM NaCl, 0.05% Tween-20), and excess buffer was carefully removed. A hybridization mix containing 2X SSC buffer, 8% deionized formamide, and 5 nM fluorescent DNA probe (5′-Atto594-(CCCTAA)5-3′, TelC probe; Eurofins Genomics) was added to the slides, which were incubated for 20 minutes at 37°C. Following hybridization, the slides were rinsed twice in 2X SSC for 5 minutes in dark cuvettes. Afterwards, slides were rinsed in ddH2O, followed by 2 minutes incubation in 70% ethanol then for 2 minutes in 100% ethanol in the dark. After the slides were air-dried, they were mounted using VECTASHIELD Vibrance Antifade Mounting Medium with DAPI. Quantification was performed on 100 cells per patient and per biological replicate across two independent replicates using a Zeiss AXIO Imager 2 microscope and ZEN 2.3 Microscopy Software. Representative images were captured using a Leica SP8 confocal microscope. Maximum intensity projections were applied using LAS X software, and further image processing was performed with both LAS X software and ImageJ 1.54f. GraphPad Prism 8 software was used to create graphs, perform statistical tests, and calculate p-values. All values from both biological replicates are shown in the violin plot (figure 3.D). Statistical significance was assessed using a nonparametric Kruskal-Wallis test, followed by Dunn’s multiple comparisons test; p-values less below 0.05 were considered statistically significant (n =2, *p < 0.05, **p < 0.01, ***p < 0.001, ****p < 0.0001).

### Quantification of ecDNAs and ploidy assessment

Metaphase spreads were prepared by adding 0.04 µg/ml colcemid to dividing cells (∼60% confluent) and cells were incubated for 8 hours at 37°C. Next, the supernatant was collected in a 15 ml Falcon tube, centrifuged and resuspended in 500 µl. 10-15 ml of a hypotonic solution (KCL 0.4% – prewarmed in a water bath) was added in a dropwise manner while gently mixing and incubated for 25 minutes at 37°C. After centrifugation, supernatant was removed, leaving 2-3 ml to resuspend the cell pellet. Cells were then fixated by adding 7-8 ml of ice cold fixative solution (Methanol 3:1 glacial acetic acid) in a dropwise manner while gently mixing, followed by centrifugation and discarding the supernatant. Fixation steps were repeated twice. After discarding the supernatant, cells were resuspended in 200-500 µl of the fixative solution, depending on the size of the pellet. For slide preparation, 20 µl of the cell suspension were dropped onto humidified pre-cleaned slides and allowed to air-dry. Slides were stored at -20°C for short-term storage and at -80°C for long-term storage. VECTASHIELD Vibrance Antifade Mounting Medium with DAPI was used for mounting the slides. Ploidy and ecDNA quantifications were performed using Zeiss AXIO imager 2 and ZEN 2.3 Microscopy Software. For ploidy counts, images were captured and analyzed using graphic tools on ZEN 2.3 Microscopy Software to identify the number of chromosomes per cell. 50 metaphases were quantified for ecDNAs and ploidy. GraphPad Prism 8 software was used to create graphs for ecDNAs, while Microsoft Excel was used to create ploidy graphs. Representative images of ecDNAs were acquired using Leica SP8 confocal microscope and data acquisition was performed using LAS X software. ImageJ 1.54f was used for further image processing.

### Molecular combing for replication dynamic analysis

Newly synthesized DNA from asynchronized cells were labeled with thymidine analog 5-iodo-2’-deoxyuridine (IdU; sigma) followed by thymidine analog 5-chloro-2deoxyurdine (CldU; Sigma) both at 100 μM diluted in medium and incubated for 30 minutes, between the analogs cells were wished three times with warm PBS. After the labelling, cells were harvested and genomic DNA extracted using Fiber-Prep kit (EXTR-001, Genomic vision), combed using the Fiber-Comb, and analyzed as previously described^65^. Fluorescence detection of the IdU and CldU analogies was done by using mouse anti-BrdU (Becton Dickinson) and rat anti-CldU (Novus Biologicals) primary antibodies followed by goat anti-mouse Alexa Fluor 488 (Invitrogen) and goat anti-rat Alexa Fluor 594 (Invitrogen) for secondary antibodies, respectively. The primary antibody for fluorescence detection of ssDNA was mouse anti-ssDNA (Millipore). The secondary antibody was donkey anti-mouse Alexa Fluor 647 (Invitrogen). The length of the replication signals and the distances between origins were measured in micrometers and converted to kilo bases according to a constant and sequence-independent stretching factor (1mM = 2 Kb), as previously reported^43^.

### Telomere length analysis using TeloSizer

Asynchronized cells from LFS041 early, crisis and late passages and from crisis passage treated with long-term nucleosides were washed once by PBS and harvested and embedded into agarose plug (50,000/plug) using Fiber-Prep kit (EXTR-001, Genomic vision), according to the manufacturer’s instructions. Two plugs from each condition were prepared, one used for quality check of the combed DNA using YOYO intercalating dye, and the other plugs were shipped at 4°C to Genomic Vision for telomere length using the TeloSizer® assay (Genomic Vision, France). Briefly, Genomic Vision melted the DNA plugs and then combed the DNA on silanized glass slides according to the EasyComb procedure (Genomic vision), the slides were then hybridized with a telomere-specific probe G-rich PNA probe Cy3-(TTAGGG)3 followed by genomic DNA staining using YOYO. Telomeres at the end of chromosomes were then scanned using a FiberVision (Genomic vision) and automatically detected and measured using the FiberStudio® Easyscan software.

### EU and EdU incorporation assays

For the EU assay, cells were incubated with 1mM of EU for 1 hour. Fixed with 3.7% PFA for 15 min, permeabilized with 0.5% Triton X-100 for 15 min and Click-iT reaction was performed according to the manufacturer’s instructions (Click-iT RNA Alexa Fluor 594 Imaging Kit, Invitrogen). For the EdU assay, cells were incubated with 1mM of EdU for 1 hour. Then, cells were fixed in 3.7% formaldehyde/PBS for 15 min, washed twice with 3% Bovine serum albumin (BSA)/PBS, permeabilized with 0.5% Triton/PBS for 20 min, washed twice with 3% BSA/PBS, incubated for 30 min in EdU detection cocktail (Click-iT EDU from ThermoFisher). For further determination of DNA damage markers, the cells were proceeded to blocking solution in 3% BSA/PBS for 1hr and then processed as Immunofluorescence staining protocol. Images were analyzed using Fiji.

### RNA content analysis using Acridine orange

Cells were analysed for RNA content as previously^66^. Briefly, asynchronized cells were washed in PBS and harvested, and then fixed in 70% ethanol, centrifuged and resuspended in PBS with 2mM MgCl2 (1.0 × 106 cells/ml). To assess the contribution of RNA, the cell solution was incubated with RNaseA (1000 units per 1ml, R6513, Sigma) for 30 minutes at 37 °C, prior to staining with Acridine orange. Then the cell solution incubated in 66% of cold solution A (1% Triton X-100, 8mMHCI, 150mM NaCl in DDW, pH 1.2) for 45 seconds. Next, it was incubated in 66% of 10 mM Acridine orange (A6014, Sigma, diluted in 1mM EDTA, 75 mL of 150mM NaC1, 126mM Na2HPO, 37mM citric acid in DDW, pH 6) and analysed immediately by flow cytometry (BD FACSAria III).

### Real-time PCR

Total RNA was extracted using the Rneasy mini kit extraction kit (QIAGEN). Complementary DNA (cDNA) synthesis was performed using the High Capacity cDNA Reverse Transcription kit (Applied Biosystems). For detecting p53β, PCR was performed using specific primers (Fwd: TGGAAACTACTTCCTGAAAACAACG, Rev: AAGCTGGTCTGGTCCTGAAG) using Platinum SuperFi II Green mix (12369010, Invitrogen) and PCR products were purified using NucleoSpin kit (740609.50, MACHEREY-NAGEL) and analyzed using 2% agarose gel. For real-time PCR, RNA-free and reverse-transcriptase free reactions were used as controls. Real-time PCR was subsequently performed in ABI 7500 using a Power SYBR green PCR master Mix (Applied Biosystems). The expression level was normalized to the transcript levels of GAPDH. Specific primers used in this study are: P53 (Fwd: CAGCACATGACGGAGGTTGT, Rev: TCATCCAAATACTCCACACGC), P21 (Fwd: GGAAGACCATGTGGACCTGT, Rev: GGCGTTTGGAGTGGTAGAAA), RRM2B (Fwd: GGACAGCAGAAGAGGTCGACTTA, Rev: AAGCTTGTTCCAGTGAGGGAGA), GAPDH (Fwd: CTGTTGCTGTAGCCAAATTCGT, Rev: ACCCACTCCTCCACCTTTGA), PSAT1 (Fwd: ACTTCCTGTCCAAGCCAGTGGA, Rev: CTGCACCTTGTATTCCAGGACC). The primers used for amplification of Ribonucleotide Reductase M2 (RRM2), c-myc, adenylosuccinate lyase (ADSL), IMP (inosine monophosphate) dehydrogenase1 (IMPDH1), dihydroorotate dehydrogenase (DHODH) are as described previously^67^.

### Metabolomic analysis of fibroblasts from LFS patients

LFS cells from early passage (p.19–p.20), as well as from healthy skin-derived fibroblasts from non-affected individuals (LFS8 and LFS9; p.14–p.15) were used. To preserve metabolite integrity and ensure measurement accuracy, plates were briefly rinsed with ddH₂O and immediately frozen with liquid nitrogen. Samples were stored at -80°C until further processing. All conditions were prepared in five biological replicates. To ensure comparable cell numbers (∼800,000–1,000,000 cells per sample), sister plates were counted, with an average of two plates per biological replicate.

Frozen plates were processed following an adjusted protocol targeting energy carriers such as NAD/NADH and NADP/NADPH^68^. This method was extended by including polarity switching and additional metabolites of interest. Briefly, cells were scratched and extracted on ice using 250 µl cooled extraction buffer (Acetonitrile : MeOH : 15 mM ammonium acetate in H2O (3:1:1), pH 10). Subsequently, samples were sonicated to ensure complete disruption of all cells using a sonication bath (Transsonic 460, Elma) for 5 min at the highest frequency on ice. Afterwards, samples were centrifuged for 15 min at 4 °C and 13,000g, and the resulting supernatant was transferred to a new LC-MS grade autosampler vial and immediately frozen at -80°C.

For metabolite separation and detection, an ACQUITY I-class PLUS UPLC system (Waters) coupled to a QTRAP 6500+ (AB SCIEX) mass spectrometer with electrospray ionization (ESI) source was used. In detail, metabolites were separated on an ACQUITY Premier BEH Amide Vanguard Fit column (100 mm × 2.1 mm, 1.7 µm, Waters) with constant column temperature of 35 °C. Separation of NAD/NADH, NADP/NADPH and additional energy carriers was achieved by the following LC gradient scheme using mobile phase A (5 mM ammonium acetate in H2O + 0.05% (v/v) ammonium hydroxide, pH 10) and mobile phase B (Acetonitrile + 0.05% (v/v) ammonium hydroxide, pH 10). An overview of multiple-reaction monitoring transitions, retention times, and MS parameters can be found in the table below. Data acquisition was performed using Analyst 1.7.2 (AB SCIEX) and processed using the OS software suite 2.0.0 (AB SCIEX).

Samples were normalized based on the cell number and results from LFS8 and LFS9 were merged. Next, log fold changes (log FC) for the mean levels of detected nucleotide metabolites in LFS041 early-passage cells compared to the mean levels in non-LFS cells were calculated. Samples were excluded from the analysis in case of no value (signal to noise (S/N) <10 or not detected). P-value for each comparison calculated using one-sided t-test (*p < 0.05, **p < 0.01, value <0.1 is indicated).

### Metabolomic analysis of BJ-*hTERT* cells

#### Sample Preparation and extraction

Asynchronized cells were washed twice with cold (4°C) PBS, and cell metabolites immediately extracted by adding 450 μl of ice-cold (-20 °C) methanol:acetonitrile:water (v/v, 5:3:2) per 25ml plate. The lysate was collected and shaked for 10 minutes. Next, the lysate was centrifuged at maximum speed for 20 minutes at 4°C. The resulting supernatant was carefully collected and transferred to HPLC vials for analysis. Quality control (QC) samples were prepared by pooling equal aliquots from each sample group. Additionally, for MS/MS analysis, each group was pooled to create an “ID sample” to facilitate metabolite identification

#### LCMS data acquisition

LCMS metabolomics analysis was performed as described previously^69^. Briefly, Dionex Ultimate 300 high-performance liquid chromatography (UPLC) system coupled to an Orbitrap Q-Exactive plus Mass spectrometer (Thermo Fisher Scientific) with a resolution of 70,000 at 200 mass/charge ratio (m/z), electrospray ionization in the HESI source, and polarity switching mode to enable both positive and negative ions across a mass range of 70 to 1000 m/z, was used. The UPLC setup included a ZIC-pHILIC column (SeQuant; 150 mm × 2.1 mm, 5 μm; Merck) with a Sure-Guard filter (SS frit 0.5 μm). Five µL of the tissue extracts were injected and the compounds were separated with a mobile phase gradient of 15 min, starting at 20% aqueous (20 mM ammonium carbonate adjusted to pH 9.2 with 0.1% of 25% ammonium hydroxide) and 80% organic (acetonitrile) and terminated with 20% acetonitrile. The flow rate and column temperature were maintained at 0.2 mL/min and 45°C, respectively, for a total run time of 26 min. All metabolites were detected using mass accuracy below 5 ppm. Thermo Xcalibur was used for the data acquisition. Data processing and analysis were performed using Compound Discoverer 3.3.2.31 with an in-house library. A total of 194 metabolites were identified based on retention time (RT) and MS2 comparisons. All statistical analyses were performed using appropriate bioinformatics tools and statistical software. Untargeted metabolomics data was processed and analysed using MetaboAnalyst 6.0. Pathway enrichment generated in MetaboAnalyst 6.0, applying a two-sample t-test for statistical comparisons, consistent with the software’s differential analysis module. Pathway impact analysis was conducted using MetaboAnalyst 6.0, with pathway significance assessed using p-value threshold of <0.05, and pathway impact scores determined based on metabolite topological analysis within each pathway.

### Proteomics analysis of LFS fibroblasts

#### Sample preparation for mass spectrometry analysis

Cell pellets from LFS fibroblasts [LFS041 p.19 (early), LFS041 p.27(crisis) and LFS041 p.346 (late)] were trypsinized with 0.25% trypsin, resuspended, and centrifuged. Two replicates were collected from each passage and the cell pellets were stored at -80°C until further processing. Cell pellets were resuspended in 150 µL of 0.1% RapiGest SF Protein Digestion Surfactant (Waters Corporation) in 100mM ammonium bicarbonate (AmBic) in H_2_O containing 10mM chloroacetamide (CAA, Sigma) and 40mM tris(2-carboxyethyl)phosphine (TCEP). The suspension was sonicated at 4°C for 15 cycles (30”ON/30”OFF) using a PicoBioruptor (Diagenode). After quantification using a BCA assay (Thermo Fisher Scientific), Samples were heated to 90°C for 5 minutes and subsequently underwent tryptic digestion (Promega) at 37°C for 18 hours. The pH was adjusted to ∼2 using trifluoroacetic acid (TFA), and samples were incubated at 37°C for 30 minutes before centrifugation at 18,000g for 30 minutes at 4°C. Next, the supernatants were moved to fresh PCR tubes, and the buffer was replaced with 5 mM triethylammonium bicarbonate (TEAB) using the SP3 protein clean-up protocol^70,71^. Peptides were labeled with TMT10plex reagents (Thermo Fisher Scientific) following the manufacturer’s instructions: Sample peptides were combined with TMT labeling reagents and incubated at room temperature for 1 hour, followed by quenching with 5% hydroxylamine for 15 minutes. The labeled peptides were combined into TMT10plex sets and speedvac was used for air-drying the samples. 100 μl TFA 0.1% was used to resuspend the sample, which were then fractionated under high pH conditions using an Agilent 1200 Infinity HPLC system equipped with a Gemini C18 column (3 µm, 110 Å, 100 × 1.0 mm, Phenomenex). Peptides were separated with a 60-minute linear gradient of 0–35% (v/v) acetonitrile in 20 mM ammonium formate (pH 10) at a flow rate of 0.1 ml/min. Peptide elution was monitored at 254 nm using a UV detector with a variable wavelength. A total of forty fractions were collected, which were then pooled into eight fractions.

#### Mass spectrometry data acquisition

0.1% trifluoroacetic acid in H_2_O was used to resuspend the dried fractions. The resuspended samples were initially loaded onto a trap column (PepMap100 C18 Nano-Trap, dimensions: 100 µm x 2 cm). Subsequently, peptide separation was performed using a 25 cm analytical column (Waters nanoEase BEH C18, dimensions: 75 μm x 250 mm, particle size: 1.7 μm, pore size: 130 Å). The chromatographic separation was carried out using a Thermo Easy nLC 1200 system (Thermo Fisher Scientific) coupled to a nanospray source. The HPLC mobile phase composition consisted of solvent A (water with 0.1% formic acid) and solvent B (80% acetonitrile, 0.1% formic acid). The elution gradient was programmed with a linear increase of solvent B: 3% to 8% over 13 minutes, 8% to 16% over 21 minutes, 16% to 50% over 119 minutes, and 50% to 95% over 10 minutes. The gradient was held at 95% for 8 minutes, followed by a decrease to 3% over the final 9 minutes. A Tri-Hybrid Orbitrap Fusion mass spectrometer (Thermo Fisher Scientific) was used to perform peptide analysis, which was operated in positive data-dependent acquisition mode with HCD fragmentation. Both MS1 and MS2 scans were acquired in the Orbitrap analyzer with a 3-second cycle time. MS1 scans were performed at a resolution of 60,000, with an AGC target of 1E6, maximum injection time of 50 ms, and a scan range of 375-1500 m/z. Peptides carrying charge states ranging from 2 to 4 were chosen for fragmentation, with a 60-second exclusion period. MS2 was carried out with a collision energy (CE) of 30%, detected in topN mode, with the first mass set at 110 m/z. The AGC target for MS2 was 2E4, with a maximum injection time of 94 ms.

#### Mass spectrometry data processing analysis and visualization

RAW data were analyzed using MaxQuant software (version 2.0.3.0), which includes the Andromeda search engine^72,73^. Peptide identification was performed by searching against the Homo sapiens Uniprot database, combined with a database of contaminant protein sequences (canonical and isoform). MaxQuant’s default parameters were applied, with the following adjustments: Trypsin/P and LysC as digestion enzymes, methionine oxidation and N-terminal acetylation as variable modifications, and cysteine carbamidomethylation as a fixed modification. The Orbitrap instrument settings included a precursor tolerance of 20 ppm and MS tolerance of 0.5 Da. A false discovery rate (FDR) of 1% was applied at both the protein and peptide levels. The match between runs option was enabled, and Label-Free Quantification (LFQ) and iBAQ values were calculated. Subsequent protein analysis was performed using Perseus software (version 1.6.2.1)^74^. The dataset was filtered to remove potential contaminants, reverse proteins, and proteins identified only by sites. Further analysis included only those proteins that were identified by at least one unique peptide in the biological replicates. Intensity values underwent normalization to correct for sample mixing errors. In cases involving multiple TMT experiments, linear modeling-based batch effect correction was applied to account for batch-induced variation. For volcano plot generation, two-sided t-test statistics were employed based on TMT quantitative information of expressed proteins, with an FDR of 0.05 and S0 constant of 0.1. Protein abundance data were transformed into Z-scores, which were calculated for each protein across the samples, and heatmaps were generated to display standardized expression patterns.

#### Downstream proteomic analysis

Proteomic data were processed by initially converting gene IDs to Ensembl IDs using g:Profiler^75^. In cases where a gene ID was not recognized by g:Profiler, we manually matched it using GeneCards^76^. Gene set enrichment analysis (GSEA) was performed using the clusterProfiler package (version 4.8.3)^77^ in R studio to to conduct GO, KEGG pathway, and Reactome pathway analyses, with gene annotations supported by org.Hs.eg.db (version 3.18.0)^78^. Pathways were considered significant if they had an adjusted p-value (FDR) below 0.05, determined by the Kolmogorov–Smirnov test with Benjamini-Hochberg correction. Plots of GSEA results were created using ggplot2 (version 3.5.1)^79^ for visualizing enriched pathways. Heatmaps illustrating protein expression patterns were generated using the pheatmap package (version 1.0.12)^80^, selecting gene sets based on specific biological processes identified in GSEA-GO results. Protein expression data were visualized by clustering genes and samples based on their expression patterns using hierarchical clustering with Euclidean distance.

### Proteomic analysis of BJ-hTERT cells

Quantitative analysis was performed with Stable Isotope Labeling with Amino Acids in Cell Culture (SILAC) labeled cells as an internal standard^81^. Labeling was performed by culturing BJ-hTERT cells in SILAC-DMEM (DMEM without the natural lysine and arginine) and supplemented with ^13^C_6_ 15N_2_-lysine, ^13^C_615_N_4_-arginine (Cambridge Isotope Laboratories) and with dialyzed FBS and antibiotics. Cells were cultured for more than 10 doublings in the SILAC medium to achieve full labeling, and the labeling incorporation was tested in a separate LC MS/MS analysis. Super-SILAC mix lysate^82^ consisted of equal amounts of labeled shp53 and control was combined with non-labeled BJ-hTERT lysate in a 1:1 ratio as a reference for quantification.

Cultured cells were lysed in a buffer including 4% SDS, 100 mM Tris HCl pH 7.6 and 100 mM DTT. Lysates were incubated for 10 min at 95 °C followed by short sonication. Each cell line (shp53 or control) was analyzed in three biological replicates and each biological replicate was processed in two different methods. One method included protein digestion using the FASP protocol^83^, followed by fractionation to 6 fractions using strong anion exchange (SAX) chromatography^84^. The second method included methanol:chloroform protein precipitation, solubilization of the proteins with 6 M urea, 2 M thiourea in 50 mM ammonium bicarbonate, in-solution digestion, followed by strong cation exchange (SCX) fractionation into 6 fractions. Finally, all peptide samples were purified on C18 stageTip^85^. The results of the two methods for each biological replicate were averaged and analyzed as a single repeat.

LC-MS/MS analysis was performed on the EASY-nLC1000 UHPLC coupled to the Q-Exactive or QExactive Plus mass spectrometers (Thermo Scientific)^86^ using EASY-Spray ionization source. Peptides were separated on a 50 cm EASY-spray PepMap column (Thermo Scientific) using a 220 min gradient of water:acetonitrile. MS analysis was performed in a data-dependent mode using a top-10 method. MS spectra were acquired at 70,000 resolution, m/z range of 300-1700 Th, a 21 target value of 3E+06 ions and a maximal injection time of 20 ms. MS/MS spectra were acquired at 17,500 resolution, a target value of 1E+05 ions, and maximal injection time of 100 ms. Raw MS files were analyzed with MaxQuant (Cox and Mann, 2008) version 1.5.0.6 and the Andromeda search engine^87^ integrated into the same version. MS/MS spectra were searched against the UniprotKB database. Database results were filtered to have a maximal FDR of 0.01 on both the peptide and the protein levels. Proteomic results were analyzed based on the normalized SILAC ratio light to heavy (L/H) after normalization by subtraction of the most frequent value in each sample. Statistical tests and calculations were done using the Perseus program. Median of technical repeats were calculated prior to analysis. Data were filtered to retain only proteins with numerical values in at least 2 of 3 biological repeats of either groups (shp53 or control). For the comparison Welch’s t-test was performed with the same parameters of permutation-based FDR=0.05 and S0=0.15^88^. Functional enrichment analysis of significantly DE proteins (Log2 fold change >1 and < -1, pvalue < 0.05) was performed against the human genome (Ensembl Genomes 58) by g:Profiler using a cumulative hypergeometric test. The g:SCS method was used for computing multiple testing with p-value<0.05 (http://biit.cs.ut.ee/gprofiler). Protein-protein interaction networks of significantly DE proteins were identified by the Cytoscape stringAP plugin (version 2.2.0) using a confidence score of 0.7. Networks were annotated with AutoAnnotate (version 1.5.2) and visualized in Cytoscape (version 3.10.3). Overlaps were adjusted using the yFiles Layout Algorithms (version 1.1.4).

### Bulk RNA-sequencing and quantification of gene fusions

Cell pellets from two LFS patients [LFS041 p.19 (early), LFS041 p.29 (crisis), LFS041 p.65 (post-crisis), LFS041 p.346 (late), LFS087 p.20 (early), LFS087 p.47 (post-crisis) and LFS087 p.195 (late)] were prepared after trypsinization in 0.25% trypsin, cell resuspension and centrifugation. The cell pellets were kept at -80°C until performing RNA extraction using AllPrep DNA/RNA/Protein Mini Kit (QIAGEN, Cat. No.: 80004), following the manufacturer’s instructions. RNA concentration was measured using NanoDrop, while a Bioanalyzer (Agilent RNA 6000 Pico Kit, Cat. No. 5067-1513) was used to measure the quality of the RNA. Sequencing libraries were prepared with the Illumina TruSeq mRNA stranded Kit following the manufacturer’s instructions. Briefly, mRNA was purified from 500 ng of total RNA using oligo(dT) beads. Poly(A)+ RNA was fragmented to 150 bp and converted to cDNA. The cDNA fragments were then end-repaired, adenylated on the 3′ end, adapter ligated and amplified (15 cycles of PCR). The final libraries were validated using the Qubit® RNA BR Assay Kit (Thermo Fisher Scientific, Cat.No. Q10211) and the TapeStation RNA ScreenTape assay (Agilent Technologies, Cat. No. 5067-5576, 5067-5577). 2x 100 bp paired-end sequencing was performed on the Illumina HiSeq 4000 following the manufacturer’s protocol. All passages from one given patient-derived culture were processed and sequenced together.

#### Data Processing and Alignment

DKFZ/ODCF RNAseq workflow (version 1.3.0) was used to analyze RNA sequencing data^89^. The specific versions of the components used include the alignment and quality control workflows (version 1.2.73-3)^90^, Roddy default plugin (version 1.2.2)^91^, Roddy base plugin (version 1.2.1)^92^, and Roddy (version 3.5.9)^91^.

In brief, a two-pass alignment approach with the STAR aligner was used to align FASTQ reads for individual samples (version 2.7.10a)^93^. The alignment was performed to a STAR index created from the 1000 Genomes assembly and gencode version 19 gene models, utilizing sjdbOverhang of 200. Duplicate reads in the main alignment file were marked using Sambamba (version 0.6.5)^94^. Samtools was used to sort the Chimeric file (version 1.6)^95^, followed by duplicate marking and generating of BAM files with Sambamba. Quality control was carried out using the Samtools flagstat command and the RNA-SeQC tool (version 1.1.8)^96^. Gene-specific read counting over exon features was conducted using Subreads (version 1.6.5)^97^, based on the Gencode v19 models. A custom script was used to compute RPKM and TPM expression values. Gene fusion detection was performed using Arriba^98^. We focused on the high-confidence fusion events.

#### Differential gene expression analysis

Differential gene expression analysis was performed using the NOISeq package (version 2.46.0)^99,100^ within R studio (version 4.3.0.) (https://rstudio. com), in order to compare between passages within individual patients. We used the following key parameters: no normalization (norm = ‘n’), a probability threshold for non-replicability (pnr = 0.2), a noise threshold (v = 0.02), and a minimum expression level (lc = 1), to ensure robust detection of significant expression changes.

GSEA GO was performed as described above. Heatmaps illustrating expression patterns were generated using pheatmap package in R studio^80^, by selecting gene sets based on specific biological processes identified in GSEA-GO results or those hypothesized to play a pivotal role in the mechanism, even if not significantly enriched in GSEA. RNA expression data were visualized by clustering genes and samples based on their expression patterns using hierarchical clustering with Euclidean distance.

### Sanger sequencing of *TP53* mutation sites in LFS

Genomic DNA was extracted from LFS cells using GenElute (G1N70-1KT, Sigma). The mutation site of the *TP53* gene was amplified by PCR with specific primers (Fwd: TACTCCCCTGCCCTCAACAA, Rev: ACAAACACGCACCTCAAAGC) using Platinum SuperFi II Green mix (12369010, Invitrogen). PCR products were purified using the NucleoSpin kit (740609.50, MACHEREY-NAGEL). Sanger sequencing was performed using inner PCR primers (Fwd: TGTGCAGCTGTGGGTTGATT, Rev: TGTTCCGTCCCAGTAGATTACC) at Hylab or Genomic Technologies Facility, Hebrew university Jerusalem. Sequencing data were analysed using FinchTV.

### Bulk whole-genome sequencing

Cell pellets from LFS fibroblasts (LFS041 p.63, post-crisis and LFS087 p.195, late) were prepared after trypsinization in 0.25% trypsin, cell resuspension and centrifugation. Cell pellets were kept at -80°C until performing DNA extraction using DNeasy Blood & Tissue Kit (QIAGEN; Cat. No.: 69504), following manufacturer’s instructions. stranded DNA assay Qubit dsDNA HS Assay Kit-100 assays (Life Technologies Q32851) was used to quantify the DNA, while a Bioanalyzer (Agilent High Sensitivity DNA 5067-4626) was used to measure the quality of the DNA. Sequencing libraries were prepared using the Illumina TruSeq HT Library Prep Kit following the manufacturer’s instructions. Briefly, 100 ng of genomic DNA was fragmented to ∼350 bp using a Covaris ultrasonicator (Covaris, Inc.). The fragmented DNA was then end-repaired, size-selected using magnetic beads, extended with an ‘A’ base on the 3′ end and ligated with TruSeq paired-end indexing adapters. The adapter-ligated fragment libraries were enriched using 8 cycles of PCR and purified 1-2 times using magnetic beads. The generated libraries were validated using the Qubit® dsDNA BR Assay Kit (Thermo Fisher Scientific, Cat.No. Q32853) and using the TapeStation Genomic DNA ScreenTape assay (Agilent Technologies, Cat. No. 5067-5365, 5067-5366). Whole genome sequencing was done using the Illumina NovaSeq 6K paired-end 150 SP platform.

#### Analysis of bulk whole-genome sequencing data

Whole-genome sequencing data were processed by the DKFZ OTP pipeline^101,102^. Briefly, reads were aligned to the 1000 Genome project version of the GRCh37 (hg19) reference genome^103^ using BWA-MEM (version 0.7.15)^104^. Duplicates were marked with Picard (version 2.25.1)^105^ and later removed with SAMtools^106^ together with low quality reads. The resultant BAM files were used as input to the analyses described below.

### OncoAnalyser

Bulk WGS derived BAM files were then run through the nf-core OncoAnalyzer nextflow pipeline (v0.5.0)^107^ based on tools developed by the Hartwig Medical Foundation, running in tumour-only mode. In brief, this pipeline calls SNVs using the HMF developed mutation-caller SAGE^108^, structural variants using GRIDSS^109^, haplotype resolved copy number variation using PURPLE^110^, and then clusters and annotates structural variant clusters using LINX^110^.

### AmpliconSuite

AmpliconSuite^111^ (v1.2.2) was run on the BAM files generated by the DKFZ OTP (described above) to identify putative highly amplified extra-chromosomal DNA fragments. CNVkit^112^ (v0.9.10) was first used to segment read counts across the genome and detect copy number changes using the circular binary segmentation algorithm^113^. Copy number altered segments with copy state of at least 4.5 and minimum length of 50 kb were then used as seeds to AmpliconArchitect^114^ to reconstruct the structures of these focally amplified regions, followed by AmpliconClassifier to obtain a list of possible circular and ecDNA structures and BFB cycles.

### Chromothripsis detection

To identify chromothriptic chromosomes, CNV calls from OncoAnalyser were first filtered for events larger than 500 bases and then used as input for the ShatterSeek R package (version 1.1)^115^ together with the corresponding somatic SVs. This method first identifies the largest SV event cluster by forming an SV-graph where individual events are connected when their breakpoints are interleaved, and then selecting the largest connected component per chromosome. The distributions of SV breakpoints and end-joining orientations are then tested against a series of criteria, looking for evidence for or against the random distribution expected in a chromothripsis event. The number of CNV events overlapping the regions affected by this SV cluster is also calculated, and this information is combined to determine a label for chromothripsis in each chromosome, as described in more detail in Cortés-Ciriano et al.^115^. For our application of this method, we used a CNV cutoff of 7 oscillating events (between 2 or 3 states), and an FDR of 0.2 for determining statistical significance. The oncoanalyser SVs and CNVs, together with the statistical criteria calculated by ShatterSeek were then visualized using the ReConPlot R package (version 0.2)^116^.

### Single-cell DNA sequencing by HIPSD-seq and HIPSD&R-seq^32^

#### Tn5 loading

A high-activity Tn5 transposase^117^ was used instead of the Tn5 transposase provided in the 10X Multiome kit. Annealing buffer (50 mM NaCl, 40 mM Tris, pH 8) was utilized to resuspend lyophilized adapter oligonucleotides at a concentration of 100 μM. The adapters were pre-annealed on a thermocycler by heating at 85°C for 2 minutes, followed by a gradual cooling to 20°C at a rate of 1°C per minute. For HIPSD-seq^32^, Read1 (5′-TCGTCGGCAGCGTCAGATGTGTATAAGAGACAG) was annealed with blocked-phos-ME (5’-[Phos]C*T*G*T*C*T*C*T*T*A*T*A*C*A*[23ddC]), while Read2 (5′-GTCTCGTGGGCTCGGAGATGTGTATAAGAGACAG) was annealed with blocked-phos-ME. For HIPSD&R-seq^32^, phos-Read1 (5′-[Phos]TCGTCGGCAGCGTCAGATGTGTATAAGAGACAG) was annealed with phos-ME (5’-[Phos]C*T*G*T*C*T*C*T*T*A*T*A*C*A*C*A*T*C*T) – phos: 5’ phosphorylation, *: PTO modification, and phos-Read2 (5′-[phos]GTCTCGTGGGCTCGGAGATGTGTATAAGAGACAG) with phos-ME. Adapters were mixed with an equal volume of 100% glycerol and stored at -20°C. Tn5 assembly involved mixing the transposase with the annealed primers in equal volumes, followed by incubation at room temperature for 30 minutes. The assembled Tn5 was subsequently diluted to a final concentration of 83 µg/ml in dilution buffer (50 mM Tris pH 7.5, 100 mM NaCl, 0.1 mM EDTA, 1 mM DTT, 0.1% NP-40, 50% glycerol) for HIPSD-seq.

#### Sample preparation

For HIPSD-seq, cells from LFS041 p.27 (crisis) and LFS041 p.62 (post-crisis) were harvested after trypsinizing the cells with 0.25% trypsin and resuspending them in PBS. For HIPSD&R-seq, cells from LFS041 p.22 (early) and LFS087 p.196 (late) were harvested in the same way, and resuspended in PBS with 1% BSA.

#### Nuclei extraction and nucleosome depletion

For HIPSD-seq, fixation was performed by incubating 1 × 10⁶ cells in 1 ml of 1.5% methanol-free formaldehyde (FA; Thermo Fisher Scientific, #28906) in PBS for 10 minutes at room temperature while gently shaking. 200 mM glycine was added for fixation neutralization, followed by ice incubation for 5 minutes. The cells were then centrifuged at 550 g for 5 minutes at 4°C and washed with ice-cold PBS. To isolate nuclei, the cells were resuspended in 1 ml of ice-cold NIB buffer (10 mM Tris-HCl; pH 7.4, 10 mM NaCl, 3 mM MgCl2, 0.1% Igepal and 1X protease inhibitor cocktail, #5871S, Cell Signaling Technology), followed by incubation on ice with gentle mixing for 20 minutes. The nuclei were centrifuged at 500 g for 5 minutes at 4°C and washed once with 1X NEBuffer 2.1 (NEB, #B7202). For the nucleosome depletion step, nuclei were resuspended in 1X NEBuffer 2.1 containing 0.3% SDS (Serva, #20767), followed by incubation at 42°C for 15 minutes with shaking. The SDS was quenched by adding 2% Triton X-100 (Sigma-Aldrich, #93443) and incubating at 42°C with shaking for 15 minutes. The nuclei were then centrifuged at 500 g for 5 minutes at 4°C and resuspended in 1X Nuclei Buffer (10X Genomics). Luna-FL™ cell counter (Logos Biosystems) was used to count the nuclei, which were then diluted to a final concentration of 2000–5000 nuclei/μl.

For HIPSD&R-seq, the protocols for fixation with 1.5% FA, nuclei isolation with NIB buffer, and nucleosome depletion using 0.3% SDS were carried out as described above for HIPSD-seq. However, all buffers used during and after nuclei isolation (i.e. to NIB buffer and 1X NEBuffer 2.1) were supplemented with 1 U/μL RNAse inhibitor (Takara Bio, #2313A). After nucleosome depletion quenching with 2% Triton X-100, nuclei were centrifuged at 500 g for 5 minutes at 4°C and resuspended in 1X Nuclei Buffer (10X Genomics) containing 1 U/μl RNAse inhibitor. Finally, the nuclei were counted using a Luna-FL™ cell counter and diluted to a concentration of 2000–5000 nuclei/μl.

#### HIPSD-seq

1.67 x 10^6^ cells from LFS041 p.27 and 1.1 x 10^6^ cells from LFS041 p.62 were used for nuclei isolation and nucleosome depletion as described earlier. The nucleosome-depleted nuclei were processed following the 10X ATAC protocol described in the Chromium Single Cell ATAC Reagent Kits User Guide v1.1 Chemistry (CG000209, 10X Genomics), with the modification that the provided transposase was substituted with a highly active in-house Tn5 (83 µg/ml). For transposition, 10,000 nuclei were loaded onto the Chromium Next GEM Chip H (PN-1000161). Single-cell DNA libraries were prepared using the standard reagents of the Chromium Next GEM Single Cell ATAC Library & Gel Bead Kit v1.1 (PN-1000176), with unique indexing applied via the Single Index Kit N Set A (PN-1000212). Qubit 3.0 Fluorometer (#Q33216, Invitrogen) and the 4200 Tapestation system (Agilent Technologies) were used for quality control and to perform molarity calculations of the final libraries. Sequencing of the libraries was performed on the NovaSeq 6000 platform with paired-end 100 SP reads: 51 cycles for read 1, 51 cycles for read 2, 8 cycles for i7, and 16 cycles for index 2, with 1% PhiX spike-in.

#### HIPSD&R-seq from fibroblasts

A total of 4.3 x 10^5^ cells from LFS041 p.22, while 5 x 10^5^ cells each from LFS041 p.67 and LFS087 p.196 were pooled (1:1 ratio)^32^ for the mixed patient experiment for HIPSD&R-seq were used for nuclei isolation and nucleosome depletion as described above. 10X Multiome protocol, according to the Chromium Next GEM Single Cell Multiome ATAC + Gene Expression Reagent Kits User Guide (CG000338, 10X Genomics) was used to process the nucleosome-depleted nuclei. For transposition, 10,000 nuclei were loaded onto the Chromium Next GEM Chip J (PN-1000230), with the 10X Genomics transposase replaced by a highly active in-house Tn5. Both scDNA and scRNA libraries were prepared using the standard reagents provided in the Chromium Next GEM Single Cell Multiome ATAC + Gene Expression Reagent Bundle (PN-1000285). The scDNA libraries were indexed with the Single Index Kit N Set A (PN-1000212), whereas the scRNA libraries were indexed with the Dual Index Kit TT Set A (PN-1000215). Qubit 3.0 Fluorometer (#Q33216, Invitrogen) and the 4200 Tapestation system (Agilent Technologies) were used for quality control and to perform molarity calculations of the final libraries. Sequencing of scDNA libraries was performed on the NovaSeq 6000 platform (200 cycles, S4) with 101 cycles for read 1, 101 cycles for read 2, 8 cycles for i7, and 24 cycles for i5. scRNA libraries were sequenced using the NovaSeq 6000 platform (paired-end 100 SP), with 28 cycles for read 1, 90 cycles for read 2, and 10 cycles each for i7 and i5. All libraries were sequenced using 1% PhiX.

#### Computational analysis

The initial preprocessing of HIPSD-seq (crisis and late passage) and HIPSD&R-seq (early passage and the mixed experiment) data was performed with Cell Ranger ATAC pipeline (version 2.1.0), with a reference provided by 10x Genomics (Cell Ranger Arc, GRCh38, version 2020-A). In the case of HIPSD&R-seq, only the DNA component was analysed. Because the default cell definition from Cell Ranger is not appropriate for samples with depleted nucleosomes, the cells were defined based on the barcode rank plot (log(fragments) vs log(rank)). Only cells above the highest gradient were used for further processing.

The initial BAM output from Cell Ranger was then split into single-cell BAM files with a command ‘bamsplice’ from Cell Ranger DNA pipeline (version 1.1.0). Next, single-cell BAM files were filtered to retain only high-quality reads with SAMtools^106^. To extract the number of counts per 1MB bin size, hmmcopy_utils was used^118^. Only cells with more than 90% of non-empty bins and 60,000 counts were kept for copy number calling with HMMcopy (version 1.42.0)^119^. When running HMMcopy, the e value was set to 0.9999. Because we previously observed a tendency of HMMcopy to overcall homozygous deletions in our data, we encoded both, homozygous and heterozygous deletions as a loss with 0-1. Finally, CNAs were plotted with ComplexHeatmap^120^, only using genomic bins that are annotated as ideal by HMMcopy.

For the mixed-patient (LFS041 p.67 : LFS087 p.196 = 1:1) HIPSD-seq experiment, the data was processed as described above. However, the next approach was used to select only the cells from the patient LFS087 p.196. Single-cell CNA profiles of all the cells were correlated (Pearson’s correlation) to bulk CNA profiles for both patients. In order for a cell to be classified as LFS087 p.196, the cell needed to have higher correlation to the bulk CNA profile of the patient LFS087 than to the CNA profile of the patient LFS041.

### Strand-seq analysis

Strand-seq libraries were prepared following the protocol detailed previously^30^ with minor modifications. The key steps of the procedure are summarized below. In brief, BrdU incorporation was carried out using growing cells from three distinct passages from LFS041 (p.22 (early), p.63 (post-crisis) and p.343 (late)). Cells were labeled with 40 µM BrdU (Sigma, B5002) for a single round of cell division. This step is crucial as cells with incomplete BrdU incorporation or those that have undergone multiple DNA synthesis phases with BrdU present will yield unreliable strand-specific sequencing data. Cells were frozen and stored at -80°C until further processing. Upon thawing in supplemented MEM medium, cells were centrifuged and resuspended in Nuclei Staining Buffer A to a final concentration of 1 x 10⁶ cells/ml. The buffer composition was as follows: 100 mM Tris–HCl (pH 7.4), 154 mM NaCl, 1 mM CaCl2, 0.5 mM MgCl2, 0.2% BSA, 0.1% NP40 (Sigma-Aldrich, Cat. No. 74385), 10 µg/ml Hoechst 33258 (Enzo Life Sciences, Cat. No. ENZ-52402) in ultra-pure water. The cell suspension was filtered through a cell strainer and incubated on ice for approximately 30 min. Single cells were then sorted into individual wells of a 96-well plate using fluorescence-activated cell sorting (FACS), each containing 5 µl of TheraPEAK ProFreezeTM Freezing Medium (Lonza, Cat, No. BEBP12-769E) and the plate was stored at -80°C until further use. Following thawing, DNA MNase fragmentation was carried out by incubating the samples with 0.5 U (Unit) Micrococcal Nuclease (NEB, Cat. No. M0247S) in MNase buffer supplemented with 1.5 mM DTT (Sigma-Aldrich, Cat. No. 43816) and 5% PEG 6000 (Calbiochem, Cat. No. 528877) for 8 min at room temperature, in a total reaction volume of 15 µl/well. The enzymatic reaction was terminated by adding 10 mM EDTA. DNA purification was performed using AMPure XP beads (Beckman Coulter, Cat. No. A63881) at a 1.0X ratio, with subsequent elution in 10 µl EB Buffer, utilizing a Biomek FXp liquid handling robotic system to facilitate large-scale library preparation. End-repair was carried out by incubating the samples for 30 min at room temperature with a mixture containing 0.3 U/μl T4 DNA polymerase (NEB, Cat. No. M0203S), 0.1 U/μl Klenow DNA polymerase (NEB, Cat. No. M0210S), and 1 U/μl T4 polynucleotide kinase (NEB, Cat. No. M0201S), 1x T4 ligase buffer (NEB, Cat. No. B0202S) and 2 mM dNTP mix (NEB, cat. no. N0447S), whereas 2.5 µl of the mixture was added to each well. Following this step, the DNA again underwent clean up using AMPure XP beads at a 1.8X ratio, employing the Biomek FXP liquid handling robotic system to maintain consistency and efficiency. Next, A-tailing was performed by adding 1.5 μl of A-tailing Master Mix to each well, which contained 1.67 U/μl Klenow Fragment (3′→5′ exo−; NEB, Cat. No. M0212S) and 1.67 mM dATP (NEB, Cat. No. N0440S). The samples were incubated at 37°C for 30 min, then purified with AMPure XP beads at a 1.8X ratio. Subsequently, adapter ligation was performed by incubating DNA fragments with forked Illumina adaptors in a master mix prepared using 0.0893 µM PE adaptors (Illumina PE adaptor-1: 5’-[Phos]GATCGGAAGAGCGGTTCAGCAGGAATGCCGAG-3′, Illumina PE adaptor-2: 5′-ACACTCTTTCCCTACACGACGCTCTTCCGATC*T-3′ – *: phosphorothioate bond between the last two nucleotides), 2.67x T4 Quick Ligation Reaction Buffer, and 0.144 U/μl Quick Ligase (NEB, Cat. No. M2200L; with the included buffer), diluted in Ultrapure H2O to a final volume of 7.5 µl per well. After adding the master mix to the DNA fragments, the reaction volume reached 20 µl, resulting in a final adaptor concentration of 33.5 nM per cell.

The ligated DNA was purified once more using AMPure XP beads at a 1.6X ratio and eluted in a volume of 9.5 µl. Subsequently, the DNA was incubated with 10 µg/ml Hoechst 33258 for 15 min, followed by UV irradiation. The irradiation was carried out using a crosslinker equipped with five 365-nm longwave UV bulbs for 15 min, delivering a total dose of 2.7 x 10³ J/m². The nicked DNA was amplified using a combination of 96-well-custom multiplexing Primer PE 2.0 (5′-CAAGCAGAAGACGGC ATACGAGATNNNNNNCGGTCTCGGCATTCCTGCTGAACCGCTCTTCCGATCT-3′), which includes a combination of oligonucleotides each containing a 6-bp (hexamer) multiplexing barcodes (Sigma-Aldrich), and Primer PE 1.0 (Illumina) (5′-AATGATACGGCGACCACCGAGATCTA­CACTCTTTCCCTACACGACGCTCTTCCGATCT-3′). First, 1 µl of Primer PE 2.0 was added per well, followed by 12.5 µl of Phusion HF PCR Master Mix (NEB, Cat. No. M0531L) and 1 µl of Primer PE 1.0, bringing the total reaction volume to 25 µL. Thermal cycling conditions were as follows: initial denaturation at 98°C for 30 seconds, followed by 18 cycles of denaturation at 98°C for 10 sec, annealing at 65°C for 30 sec, and extension at 72°C for 30 sec. A final extension step was carried out at 72°C for 5 min. Post-amplification, the DNA from all wells was pooled and subjected to purification and size selection using AMPure XP beads at a 0.8X ratio. This step served to remove free primers and adapter dimers. The resulting libraries were sequenced on a NextSeq500 platform using a MID-mode, 75bp paired-end protocol. Following sequencing, the data was demultiplexed, aligned to the hg38 reference assembly using BWA mem (0.7.17-r1188)^104^, and duplicate reads were mapped using sambamba (v1.0)^92^.

#### Strand-seq data processing

Strand-seq data was processed using the MosaicCatcher pipeline (v2.0.1)^30^. Mapping quality filtered (MAPQ > 10) and non-duplicated reads were used to generate read-counts per 200kb bin for Watson and Crick strands separately in each cell. Read counts per bin were then visualized per chromosome and cell. Prior to downstream analysis, cells were manually filtered based on their visualized read count distributions to exclude noisy cells (those exhibiting spiky peaks, high background or uneven coverage), cells with incomplete BrdU incorporation, cells with fewer than 200,000 reads, and those displaying all-Watson or all-Crick chromosomes or extremely skewed Watson/Crick ratios (>80% Watson or >80% Crick). For the remaining cells, the read counts per strand and chromosome were segmented, haplotype-resolved using StrandPhaseR and fed into MosaiClassifier to create initial SV calls per cell.

Ideograms generated using Mosaicatcher were used to profile structural variations, including deletions, duplications, inverted duplications, inversions, and complex rearrangements, in single cells across different passages using Microsoft Excel. Each structural variant was mapped to its genomic position in megabase resolution for individual chromosomal arms after detailed examination of each cell. This comprehensive approach facilitated variant calling and enabled us to create a matrix of all possible structural variations for each cell.

#### Clustering of Single Cells Based on Structural Variants

Next, we took two approaches to cluster single cells based on structural variant annotations per-cell detected from StrandSeq (as described above). The first approach was event-based. For this approach, unique breakpoints detected across all cells were used to segment the genome. For each cell, the presence or absence and the type of structural variant at each segment was recorded. The second approach was bin-based. Here the genome was binned into approximately 2 MB bins, with equal bin-sizes per chromosome derived from the tile function as implemented by the GenomicRanges R package (v1.52.0)^121^. For each cell and segment derived from binning, a structural variant was recorded if it was detected by Strand-seq to overlap that segment within that cell. In the case of two or more structural variants overlapping the same segment, the SV overlapping the greater number of bases within the segment was assigned. In both cases, a virtual diploid cell which is unmodified at each genomic segment was added for subsequent rooting of phylogenetic trees. For both genome segmentation approaches, distance between cells was then calculated using a modified Hamming distance, where for each pair of cells, each segment contributed a distance of 1 if the event recorded for that segment did not match, except for a comparison between unmodified and complex segments, which contributed a distance of 2. This was chosen to account for the fact that the “complex” structural variant annotation as described above can arise from the overlap of multiple simple SVs in the same genomic location. The cells were then clustered using the neighbor joining method as implemented by the ape R package (v5.7-1). For visualization purposes, the ape package was used to plot the resulting clustering as phylogenetic trees, rooted at the virtual diploid cell added to the data. Finally, main structural variance differences and shared events between main clusters were annotated manually based on the matrices created earlier using Excel.

#### Fold-back inversions calling (BFB patterns) in strand-seq data

We analyzed the strand-seq data to identify fold-back inversions in single cells, which are hallmark features of BFB events^122^. Fold-back inversions were quantified by searching for characteristic patterns on all chromosomal arms, defined as small amplifications involving either the H1 (haplotype 1), H2 (haplotype 2) strand or both, followed by a deletion. The classification of fold-back inversions and their connection to BFB cycles was guided by previously defined structural rearrangement footprints^122^. GraphPad Prism 8 software was used to create graphs, perform statistical tests and calculate p-values. Fisher’s exact test was performed for pairwise comparisons between each of the three passages. To account for the number of passages (n = 3), results were normalized accordingly, and p-values were adjusted as comparisons were made across the three passages.

### Analysis of hypertranscription in a public dataset

We re-analyzed data from a study on hypertranscription, which encompasses RNA output levels from 7494 patients from The Cancer Genome Atlas (TCGA) across 31 cancer types^54^. The authors of this study calculated the hypertranscription scores using the RNAmp method, which compares the allelic ratio of somatic mutations in RNA to their corresponding ratio in DNA, leveraging the increased representation of transcribed alleles in RNA compared to DNA. The patient IDs of the same 7494 patients were used to classify them into two groups based on their respective *TP53* and *MDM2* status from the publicly available TCGA dataset through the Genomic Data Commons (GDC) portal. The first group included tumours with WT *TP53* and *MDM2* (WT *TP53/MDM2*), while the second group consisted of tumours with *TP53* loss, *MDM2* gain, or both (*TP53^-^/MDM2^+^/TP53^-^MDM2^+^*), which are referred to as *TP53^-^/MDM2^+^* in the main text. Both heterozygous and homozygous CNAs were considered. After filtering, the dataset contained 6,305 tumour samples, split into 4032 *TP53^-^/MDM2^+^* and 2273 WT *TP53/MDM2* tumours.

The fold increase in RNA output, namely the hypertranscription fold change (hytx_fch) was used to compare the hypertranscription status between the two groups. The comparison was performed based on the total number of patients, regardless of the tumor type, as well as for each tumor type separately. For comparisons within each cancer type, a minimum sample size of 50 patients per group was required to ensure statistical robustness. Tumor types that did not meet this criterion were excluded from the analysis. A subset of selected tumour types is presented in (**Supplementary Information, Fig. S10g**). GraphPad Prism 8 software was used to create graphs, perform statistical tests and calculate p-values. Statistical significance was assessed using a non-parametric t-test (Mann-Whitney test) to compare two independent groups, as the data were not normally distributed. P-values less than 0.05 were considered statistically significant (*p < 0.05, **p < 0.01, ***p < 0.001, **p < 0.0001).

### Statistics and reproducibility

Statistical analyses employed tests described in the respective methods sections and specified in the figure legends or the main text as appropriate. Statistical significance was defined as a p-value less than 0.05, and multiple testing corrections were applied when needed. Sample sizes were not predetermined using statistical methods, and all collected data were included in the analyses. Detailed information on biological replicates and experimental repetitions is provided in the respective figure legends.

Graphs, statistical analyses, and p-value calculations were conducted using GraphPad Prism 8 or 10 software, unless otherwise stated. All values are reported as mean ± standard error of the mean (SEM) or mean ± standard deviation of the mean (SD) as indicated.

## Data availability

Source data are provided with this paper (EGAS00001008074). All other data supporting the findings of this study are available from the corresponding author on reasonable request.

## Acknowledgements

The authors thank Maayan Roniger for generating some of the pilot data for this project., Michalea Hergt and Michele Vousten for excellent technical support, Anne-Sophie Aleixo for support with the ALT-FISH, Axel Benner for advice on the statistical analyses, Gernot Poschet and Glynis Klinke for support with the metabolomics analyses and the entire Kerem and Ernst research groups for fruitful discussions and ideas. W.Z. acknowledges the support by the Neubauer Foundation for the Ph.D. fellowship. This research was partially supported by: the cooperation program in Cancer Research of the Deutsches Krebsforschungszentrum (DKFZ) and Israel’s Ministry of Science and Technology (MOST) to BK and AE, the Israel Cancer Association to BK, the Israel Science Foundation (grants No. 1284/18 and 1616/24) to BK, the DFG to AE (grant No. 460595631).

