## Supplementary Information for "P53 deficiency triggers hypertranscription, inducing nucleotide insufficiency causing replication stress, and genomic instability"

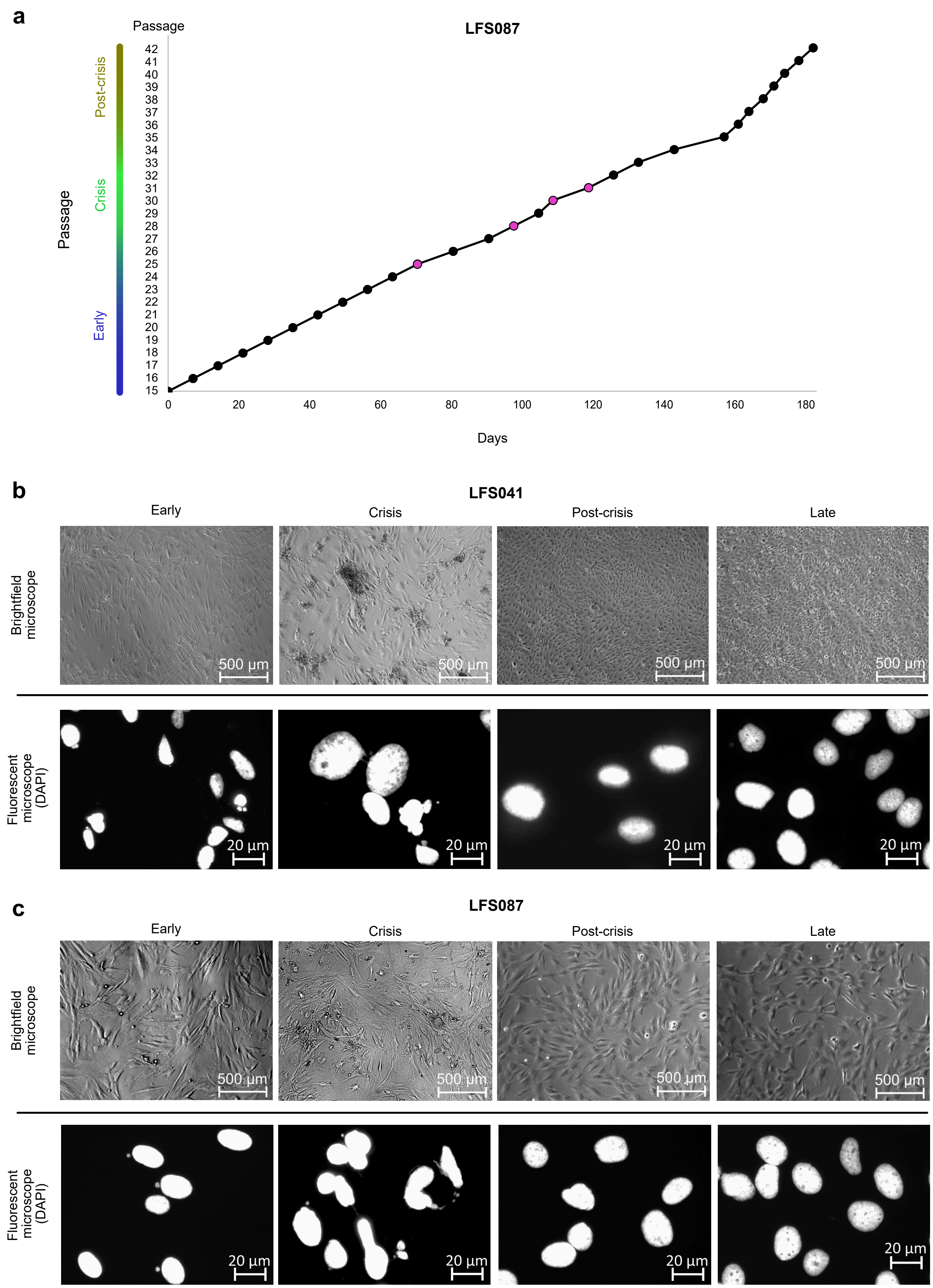
a
LFS087
Passage
Post-crisis
Crisis
Early
42
41
40
39
38
37
36
35
34
33
32
31
30
29
28
27
26
25
24
23
22
21
20
19
18
17
16
15
0
20
40
60
80
100
120
140
160
180
Days
b
LFS041
Early
Crisis
Post-crisis
Late
Brightfield microscope
500 µm
Fluorescent microscope (DAPI)
20 µm
c
LFS087
Early
Crisis
Post-crisis
Late
Brightfield microscope
500 µm
Fluorescent microscope (DAPI)
20 µm

Figure S1 a-c

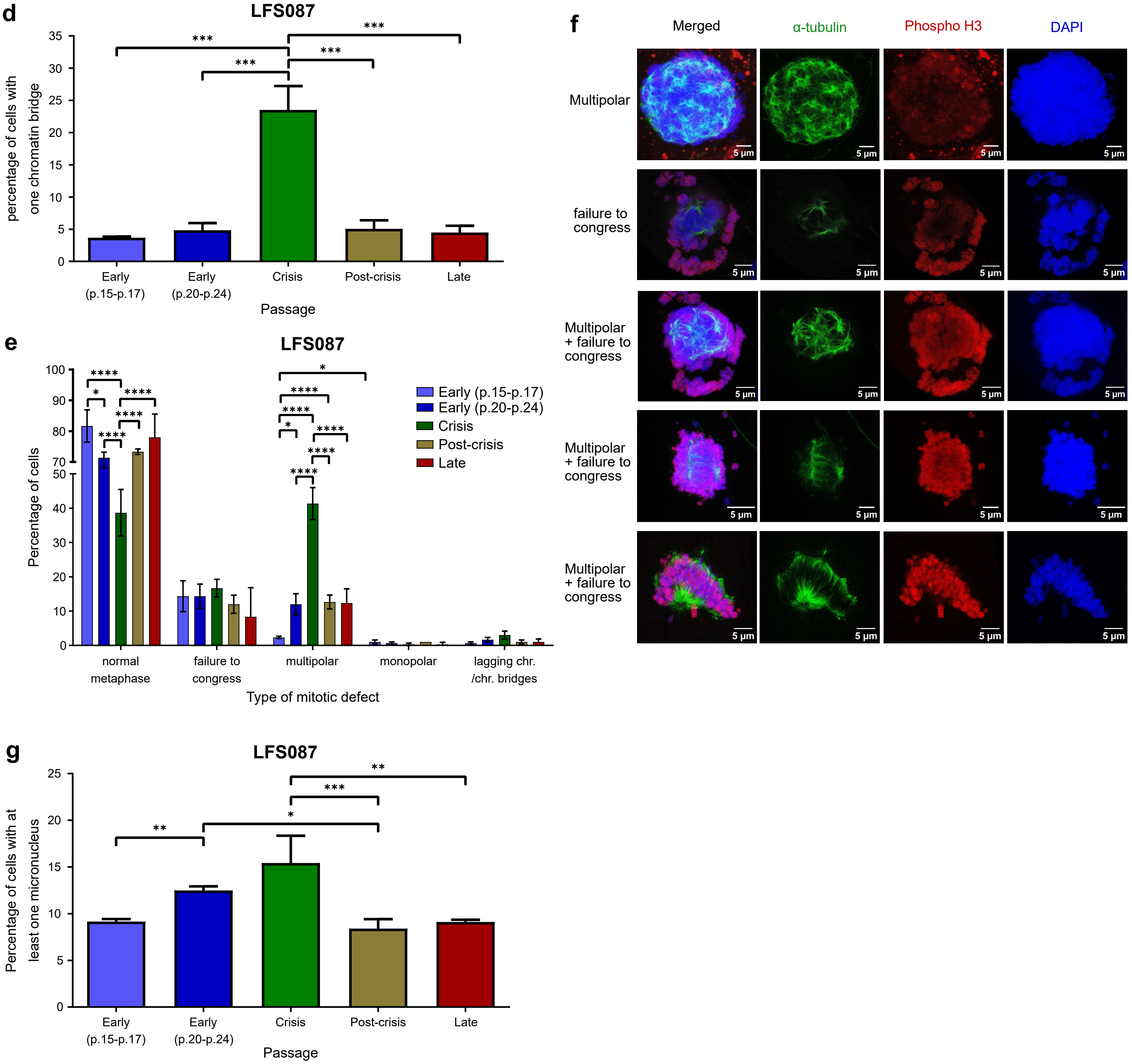
d
LFS087
percentage of cells with one chromatin bridge
35
30
25
20
15
10
5
0
***
***
***
***
Early (p.15-p.17)
Early (p.20-p.24)
Crisis
Post-crisis
Late
Passage
e
LFS087
Percentage of cells
100
90
80
70
50
40
30
20
10
0
****
*
****
****
****
*
****
****
*
****
****
****
Early (p.15-p.17)
Early (p.20-p.24)
Crisis
Post-crisis
Late
normal metaphase
failure to congress
multipolar
monopolar
lagging chr. /chr. bridges
Type of mitotic defect
f
Merged
α-tubulin
Phospho H3
DAPI
Multipolar
failure to congress
Multipolar + failure to congress
Multipolar + failure to congress
Multipolar + failure to congress
5 µm
g
LFS087
Percentage of cells with at least one micronucleus
25
20
15
10
5
0
**
***
*
**
Early (p.15-p.17)
Early (p.20-p.24)
Crisis
Post-crisis
Late
Passage

Figure S1 d-g

**Figure S1. Phenotypic characteristics of LFS skin-derived fibroblasts.**

**a.** Growth curve for fibroblasts from patient LFS087. The growth curve depicts the time required for cells to reach confluency (x-axis) from passage 15 (early passage), passing through the growth crisis, and continuing to passage 42 (post-crisis) (y-axis) for patient LFS087. Black circles: splitting ratio 1:3; purple circles: splitting ratio 1:2. **b.** and **c.** Representative images showing the cellular morphology (upper panels) as well as DAPI staining (lower panels) from different passages (early, crisis, post-crisis and late) for two LFS patients, LFS041 and LFS087, respectively. Brightfield images were acquired using a 5x magnification objective on an inverted microscope (scale bar: 500 µm), while immunofluorescent images were taken at 63x magnification objective (scale bar: 20 µm). **d.** Quantification of chromatin bridges for patient LFS087. Three independent biological replicates were performed for each condition: early (p.15-p.17) (mean: 3.7 ±0.2), early (p.20-p.24) (mean: 4.8 ±1.1), crisis (mean: 23.5 ±3.7), post-crisis (mean: 5.1 ±1.3), and late (mean: 4.5 ±1.1). For each replicate, 750 cells were quantified. **e.** Quantification of micronuclei for patient LFS087. Three independent biological replicates were performed for each condition: early (mean: 9.156 ±0.1602), early (mean: 12.49 ±0.2475), crisis (mean: 15.42 ±1.691), post-crisis (mean: 8.400 ±0.5812), and late (mean: 9.1 ±0.2). For each replicate, 750 cells were quantified. **f.** Representative immunofluorescence pictures showing examples of mitotic defects. Scale bar: 5 µm. **g.** Quantification of mitotic defects for patient LFS087. Three independent biological replicates were performed for each condition. For each replicate, 100 cells were quantified. Lagging chr./chr. bridges: lagging chromosomes/ chromatin bridges. In **d** and **e,** data are presented as mean ± SEM. Statistical significance was assessed using a one-way ANOVA followed by Tukey's multiple comparisons test. In **g,** data are presented as mean values. Statistical significance was assessed using repeated-measures two-way ANOVA followed by uncorrected Fisher’s LSD for multiple comparisons. P-values below 0.05 were considered statistically significant ($n = 3$ per group, *$p < 0.05$, **$p < 0.01$, ***$p < 0.001$, ****$p < 0.0001$).

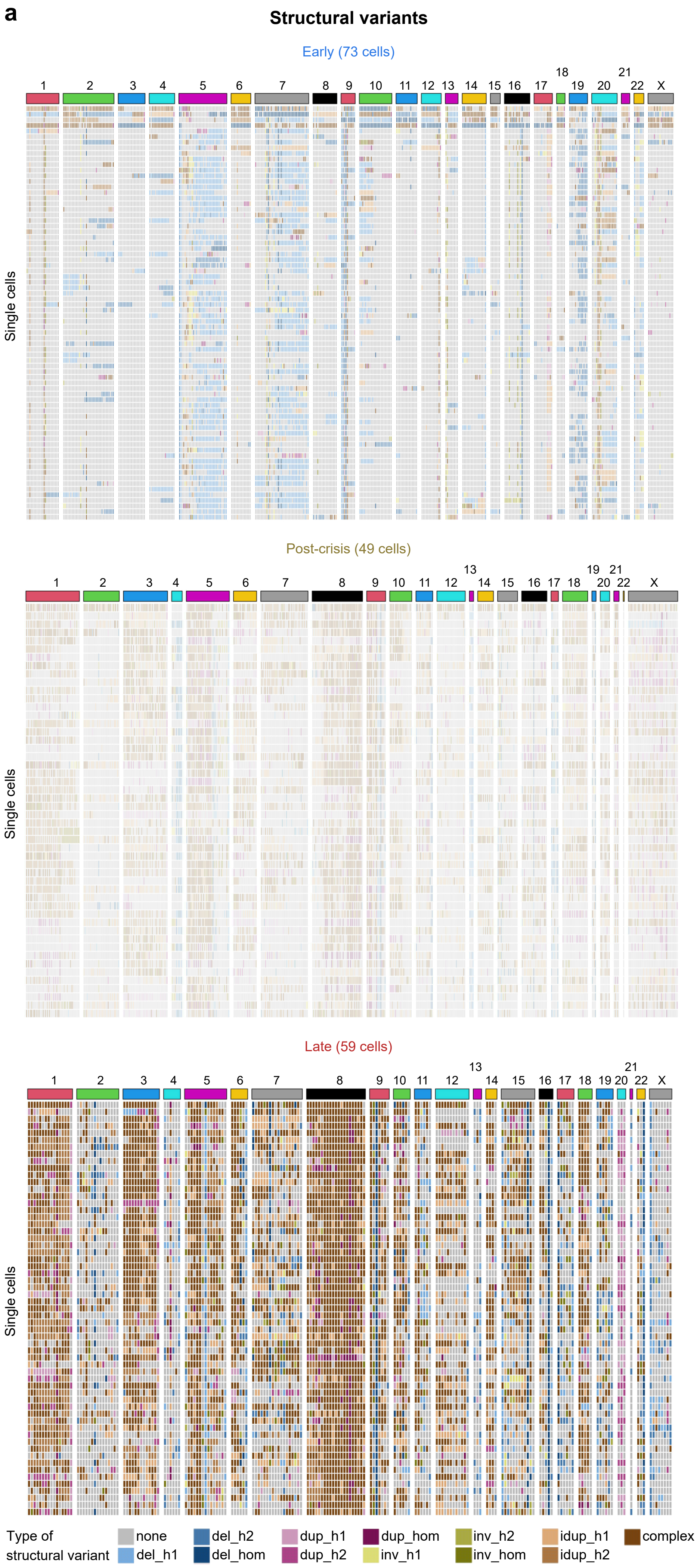

a
Structural variants
Early (73 cells)
Post-crisis (49 cells)
Late (59 cells)
Single cells
1 2 3 4 5 6 7 8 9 10 11 12 13 14 15 16 17 18 19 20 21 22 X
Type of structural variant
none
del_h1
del_h2
del_hom
dup_h1
dup_h2
dup_hom
inv_h1
inv_h2
inv_hom
idup_h1
idup_h2
complex


Figure S2 a

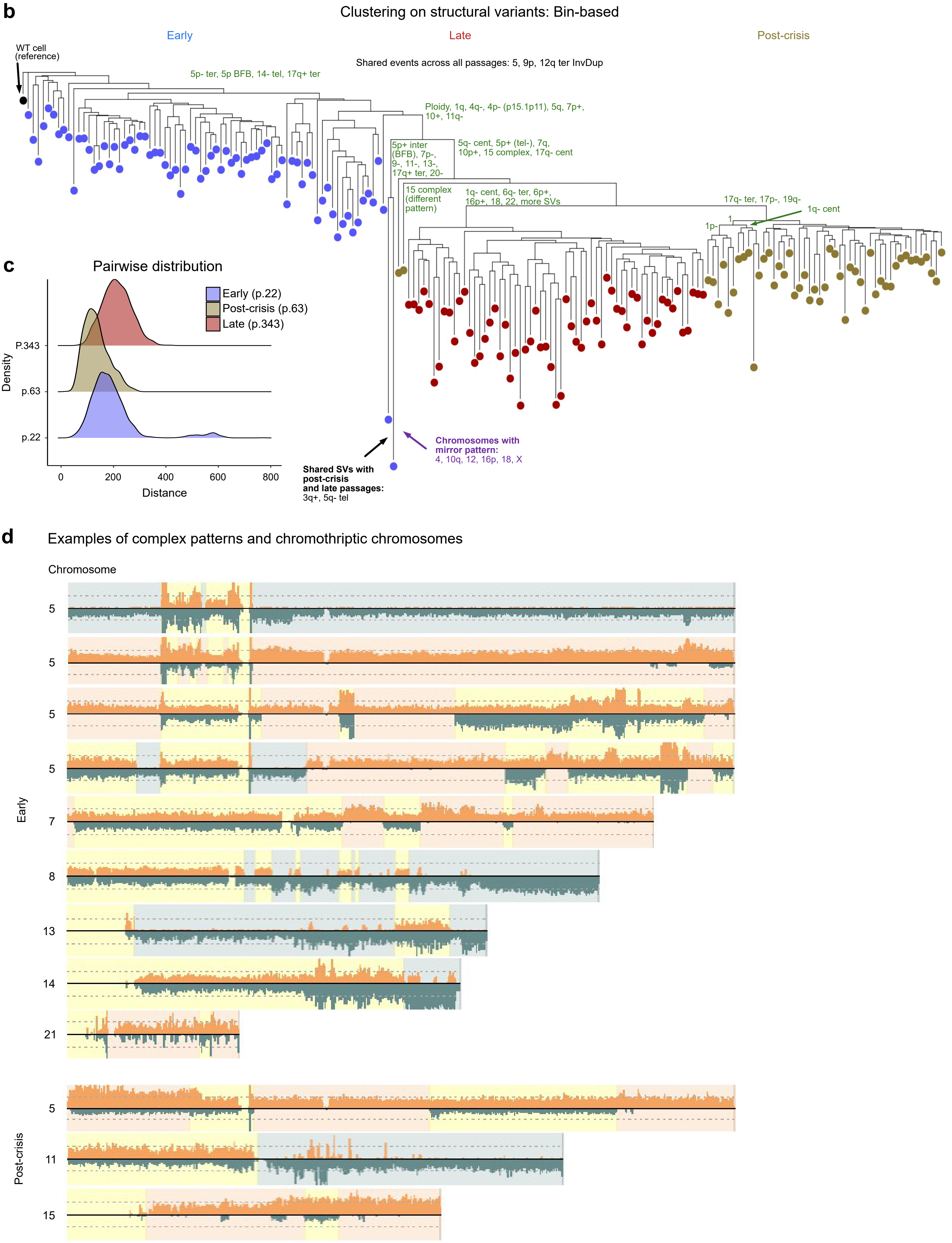
b
Clustering on structural variants: Bin-based
Early
Late
Post-crisis
WT cell (reference)
Shared events across all passages: 5, 9p, 12q ter InvDup
5p- ter, 5p BFB, 14- tel, 17q+ ter
Ploidy, 1q, 4q-, 4p- (p15.1p11), 5q, 7p+, 10+, 11q-
5p+ inter (BFB), 7p-, 9-, 11-, 13-, 17q+ ter, 20-
5q- cent, 5p+ (tel-), 7q, 10p+, 15 complex, 17q- cent
15 complex (different pattern)
1q- cent, 6q- ter, 6p+, 16p+, 18, 22, more SVs
17q- ter, 17p-, 19q-
1q- cent
1p-
1
Shared SVs with post-crisis and late passages: 3q+, 5q- tel
Chromosomes with mirror pattern: 4, 10q, 12, 16p, 18, X
c
Pairwise distribution
Early (p.22)
Post-crisis (p.63)
Late (p.343)
Density
P.343
p.63
p.22
0
200
400
600
800
Distance
d
Examples of complex patterns and chromothriptic chromosomes
Chromosome
Early
5
5
5
5
7
8
13
14
21
Post-crisis
5
11
15

Figure S2 b-d

**e** Strand-seq data with complex patterns similar to post-crisis and late passages

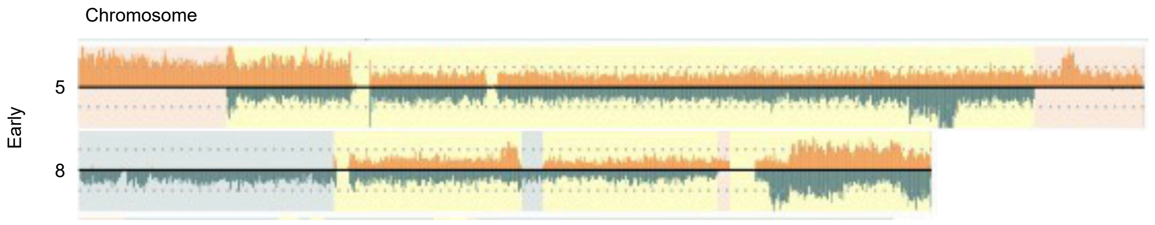


**f** **HIPSD-seq – LFS087**

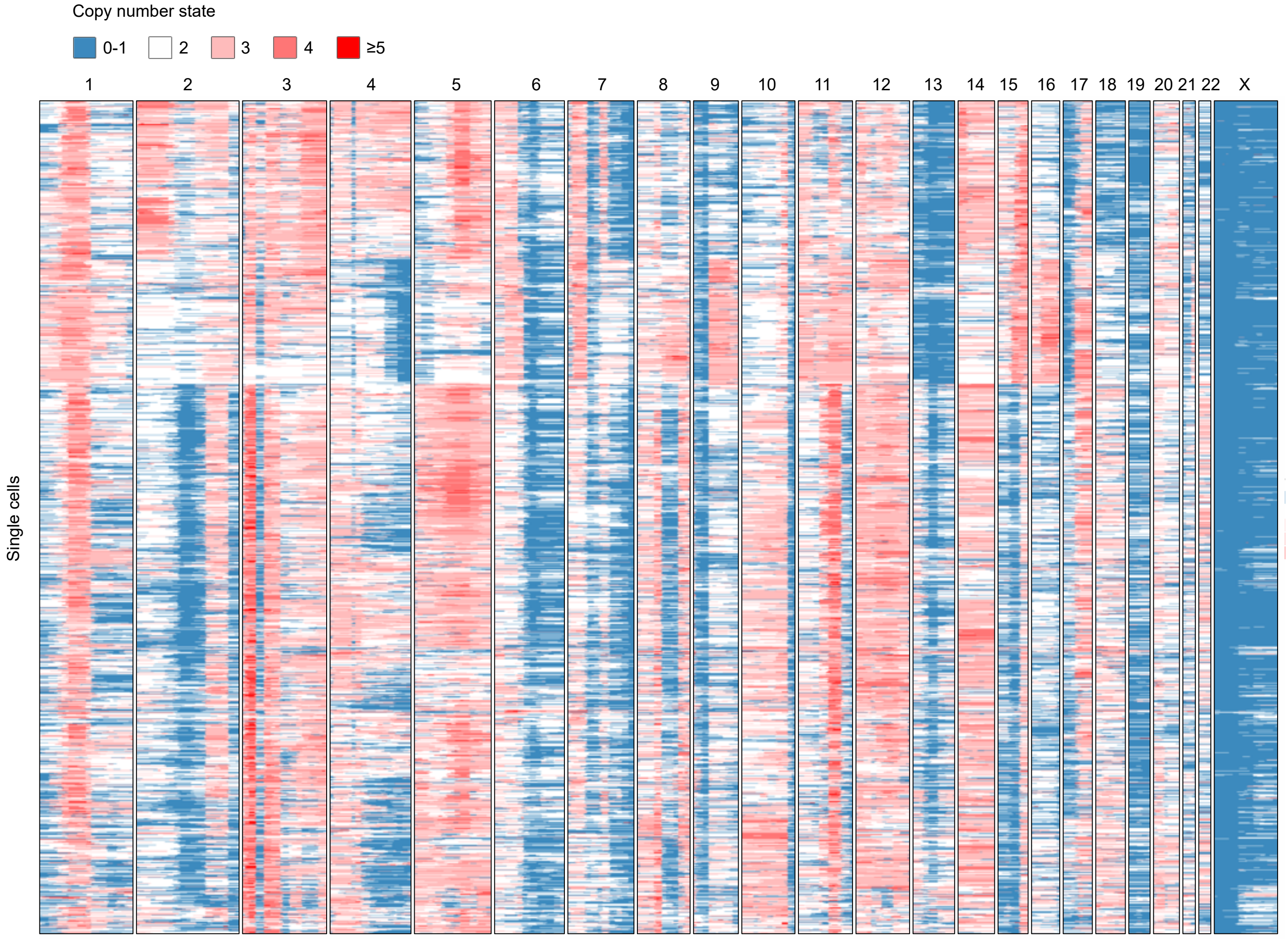


Figure S2 e-f

**<u>Figure S2.</u> LFS fibroblasts show complex rearrangements already at early passages.**

**a.** Heatmaps showing structural variants per chromosome for each passage (strand-seq data). Each row shows one cell. Deletions are depicted in blue, amplifications and complex rearrangements in orange and brown, respectively. Del: deletion, dup: duplication, inv: inversion, idup: inverted duplication, h1: homolog 1, h2: homolog 2, hom: homologous. **b.** Examples of complex rearrangements and chromothriptic chromosomes from early and post-crisis-passage cells of patient LFS041. **c.** Clustering on structural variants detected in strand-seq data using modified Hamming distances on 2MB bins and neighbour joining hierarchical clustering (**Methods**). Structural variants separating the main cell clusters are shown on the nodes. Shared events are annotated in black, while differences between the main clusters are annotated in green. Chromosomes with mirror patterns between cells are described in purple. **d**. Density plot illustrating the distribution of pairwise distances among fibroblast cells from early (p.22), post-crisis (p.63), and late (p343) passages. The x-axis represents pairwise genetic distances, while the y-axis indicates the density of cells exhibiting a given distance value. **e.** Representative examples of plots from Strand-seq data for chromosomes 5 and 8 from early passage cells (LFS041 p.22) that show similar complex rearrangements to those detected as clonal in post-crisis and late passages (see Figure 2C). **f.** HIPSD-seq heatmap (bin size of 1000 kb) of LFS087 p.196 (late, n=1869 cells) showing copy-number variation. Each row represents one cell.

**a** Bulk WGS – LFS041 p.63 (post-crisis) – Circos plots

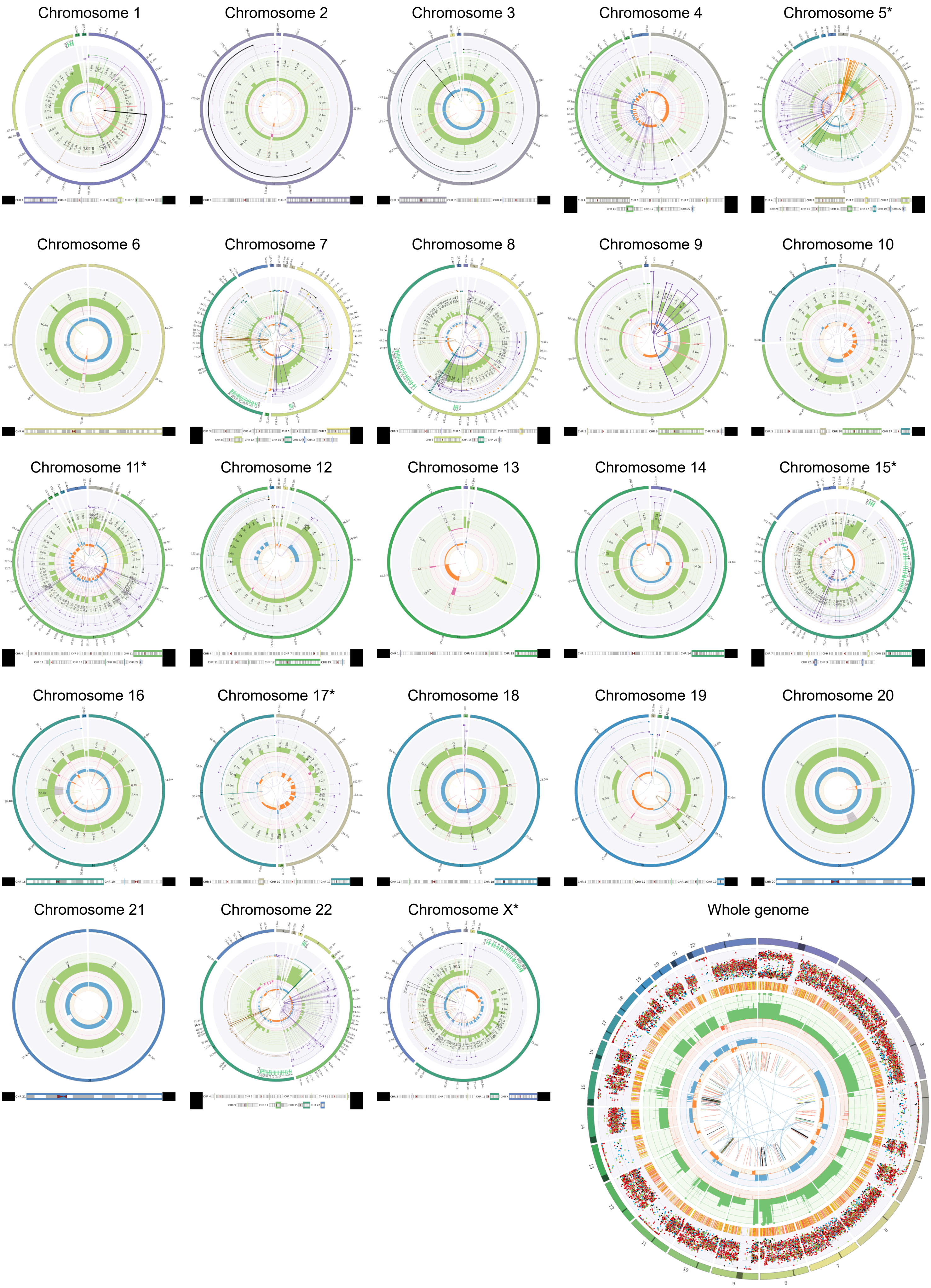


Figure S3 a

**b** Bulk WGS – LFS087 p.195 (late) – Circos plots

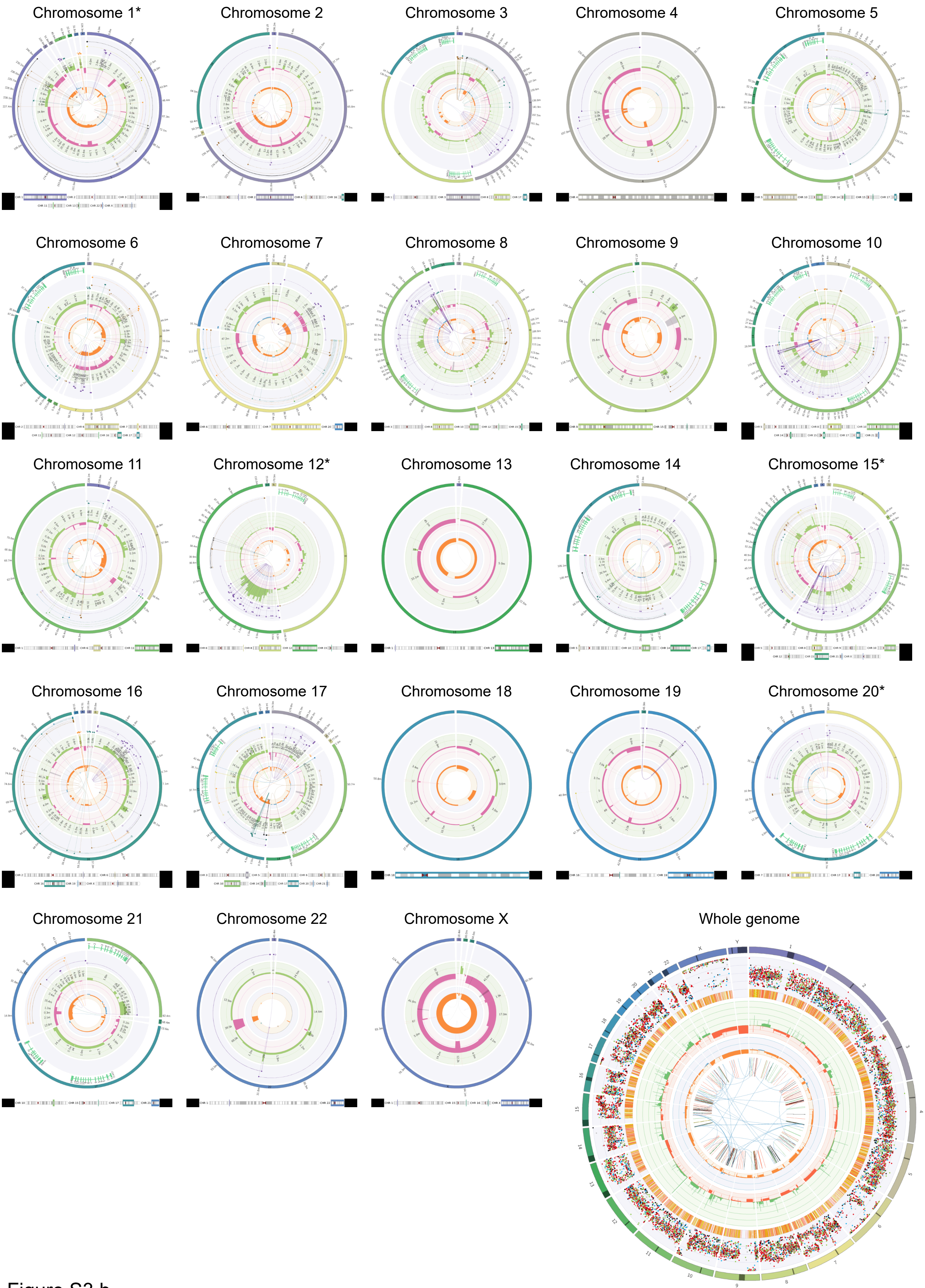


Figure S3 b

**c** Bulk WGS – LFS041 p.63 (post-crisis) – ReConPlot (copy number plots)

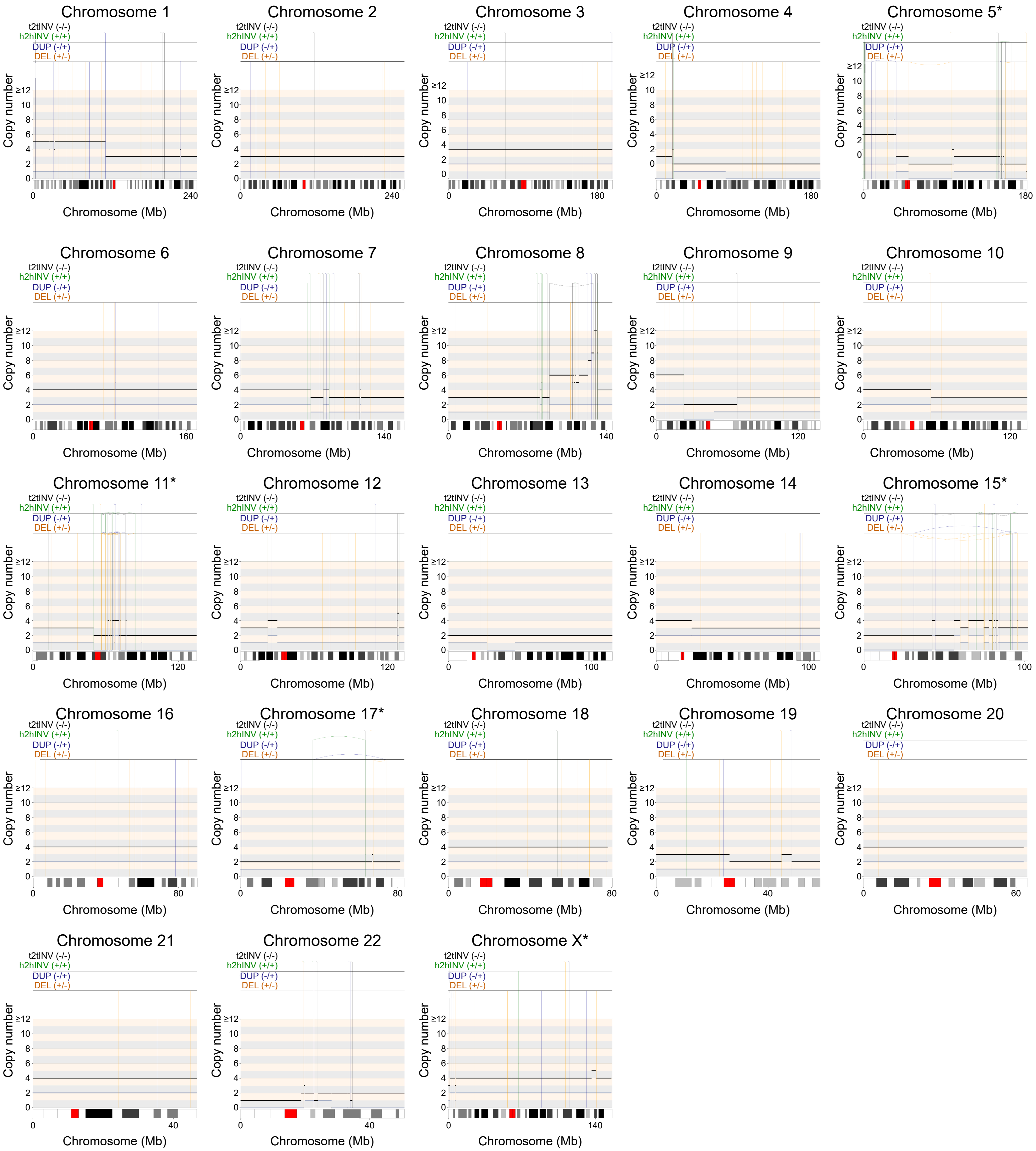


Figure S3 c

**d** Bulk WGS – LFS087 p.195 (late) – ReConPlot (copy number plots)

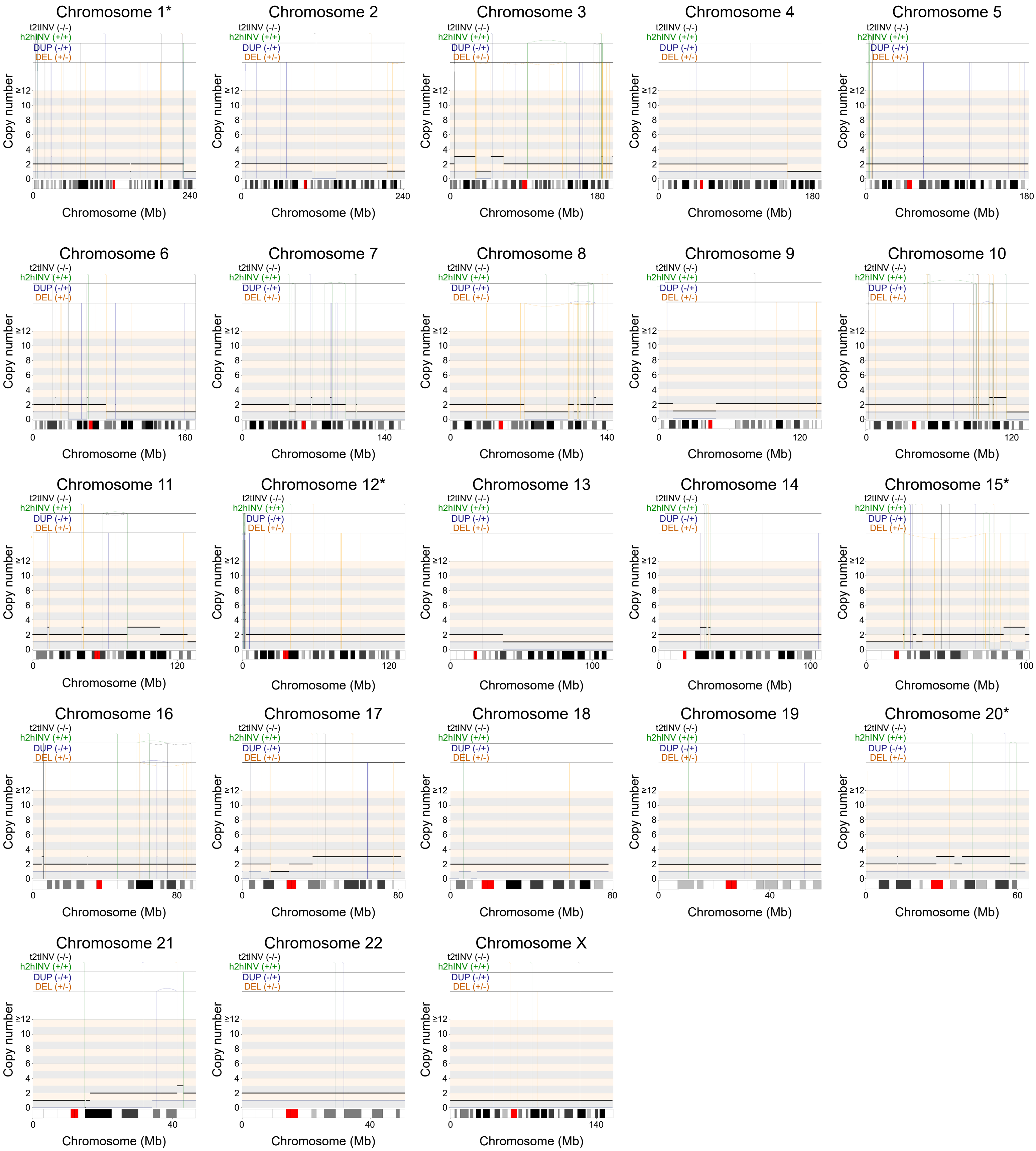


Figure S3 d

**Figure S3.** **Bulk WGS data for LFS041 and LFS087.**

**a, b.** Circos plots depicting somatic alterations identified from whole-genome sequencing (WGS) data of LFS041 p.63 and LFS087 p.195, respectively using OncoAnalyser. Each plot illustrates the distribution of somatic mutations, copy number variations and structural variants of each chromosome and across the whole genome. In the whole genome Circos plot, the outer track shows the chromosomes, followed by somatic variants (SNP/allele frequency), somatic variants (short indel locations), copy number changes and the inner circle exhibits minor allele copy numbers, while the inner connections indicate relationships between chromosomal regions affected by structural variants. **c, d.** ReConPlots showing the copy number variants (CNVs) identified from WGS data for patient LFS041 p.63 and LFS087 p.195, respectively. Each plot represents the distribution of CNVs along each chromosome, with color coding indicating sites of deletions (orange), duplications (blue), head-to-head inversions (green) and tail-to-tail inversions (black). Total chromosome length is shown in megabase (Mb). Chromosomes annotated with an asterisk (*) indicate chromothriptic chromosomes. t2tINV: tail-to-tail inversion, h2hINV: head-to-head inversion, DUP: duplication and DEL: deletion.

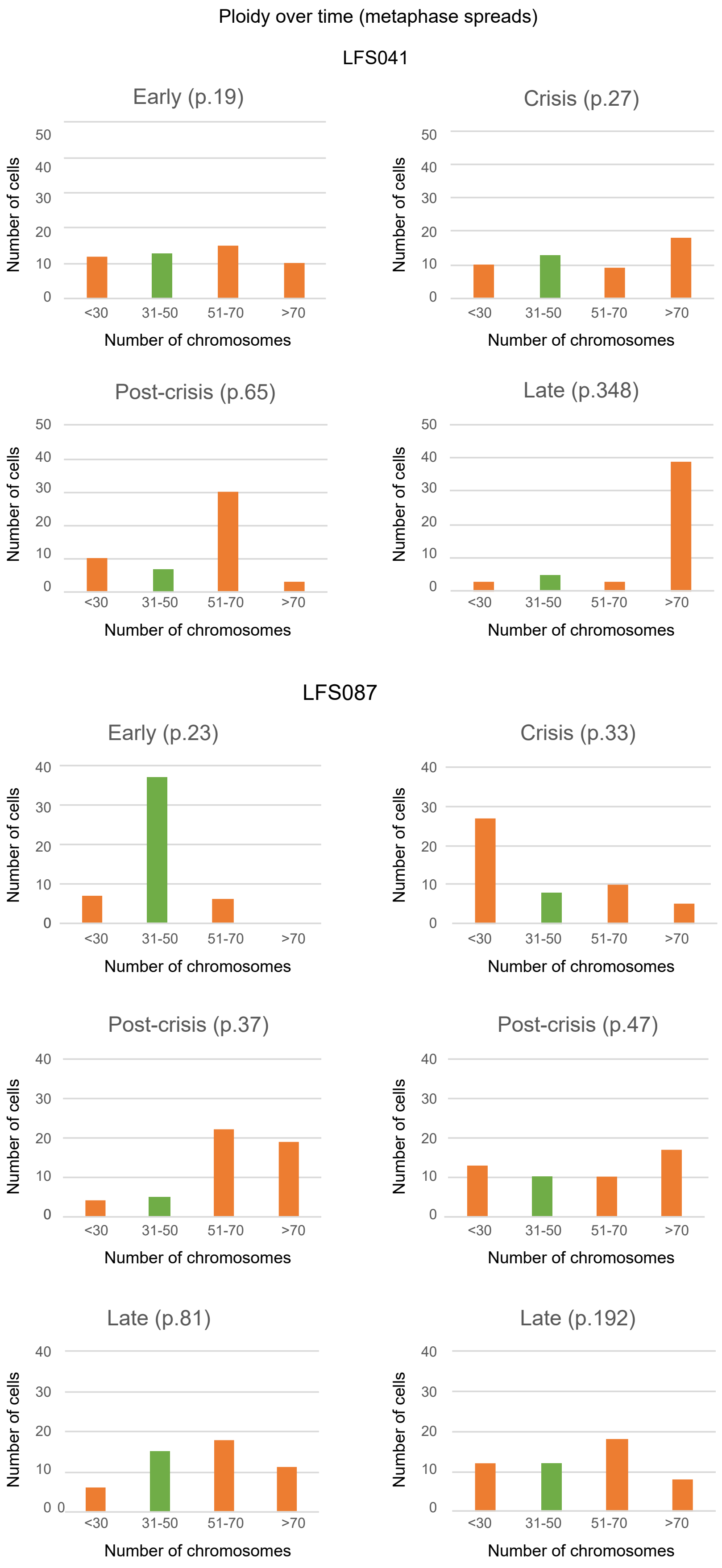


**Figure S4.** **Ploidy assessed by counting chromosomes on metaphase spreads for patients LFS041 and LFS087 shows higher numbers of polyploid cells in post-crisis and late passages as compared to early passage cells.** Chromosomal counts were obtained from 50 cells per replicate for patients LFS041 and LFS087. Bar graphs display the distribution of chromosome numbers per cell. Green bars represent near-diploid cells (31-50 chromosomes), while orange bars indicate non-diploid populations (<30, 51-70, and >70 chromosomes).

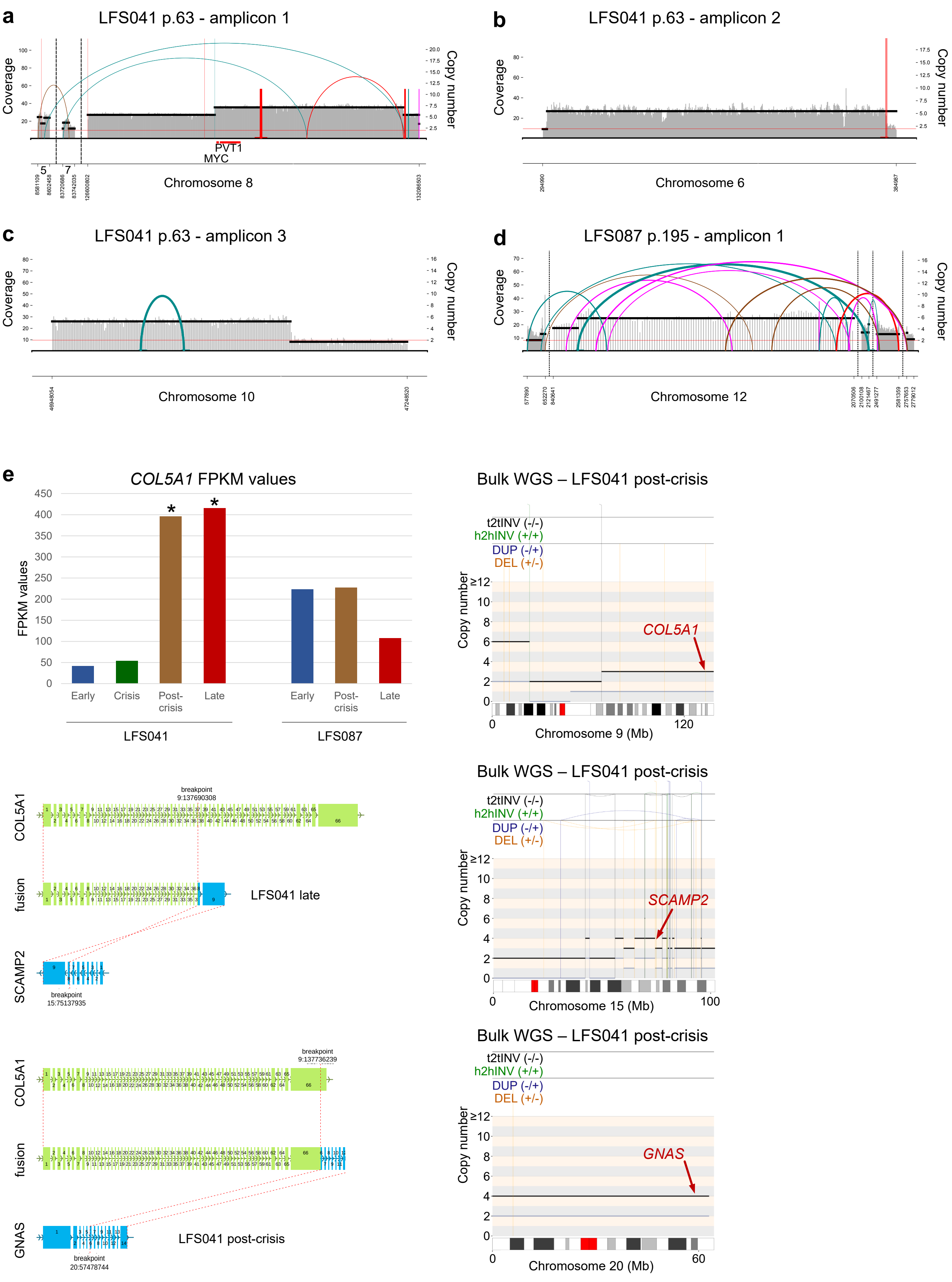
a
LFS041 p.63 - amplicon 1
Coverage
Copy number
PVT1
MYC
Chromosome 8
b
LFS041 p.63 - amplicon 2
Chromosome 6
c
LFS041 p.63 - amplicon 3
Chromosome 10
d
LFS087 p.195 - amplicon 1
Chromosome 12
e
COL5A1 FPKM values
FPKM values
Early
Crisis
Post-crisis
Late
LFS041
LFS087
Bulk WGS – LFS041 post-crisis
t2tINV (-/-)
h2hINV (+/+)
DUP (-/+)
DEL (+/-)
COL5A1
Chromosome 9 (Mb)
SCAMP2
Chromosome 15 (Mb)
GNAS
Chromosome 20 (Mb)
breakpoint
9:137690308
fusion
LFS041 late
breakpoint
15:75137935
breakpoint
9:137736239
LFS041 post-crisis
breakpoint
20:57478744

Figure S5 a-e

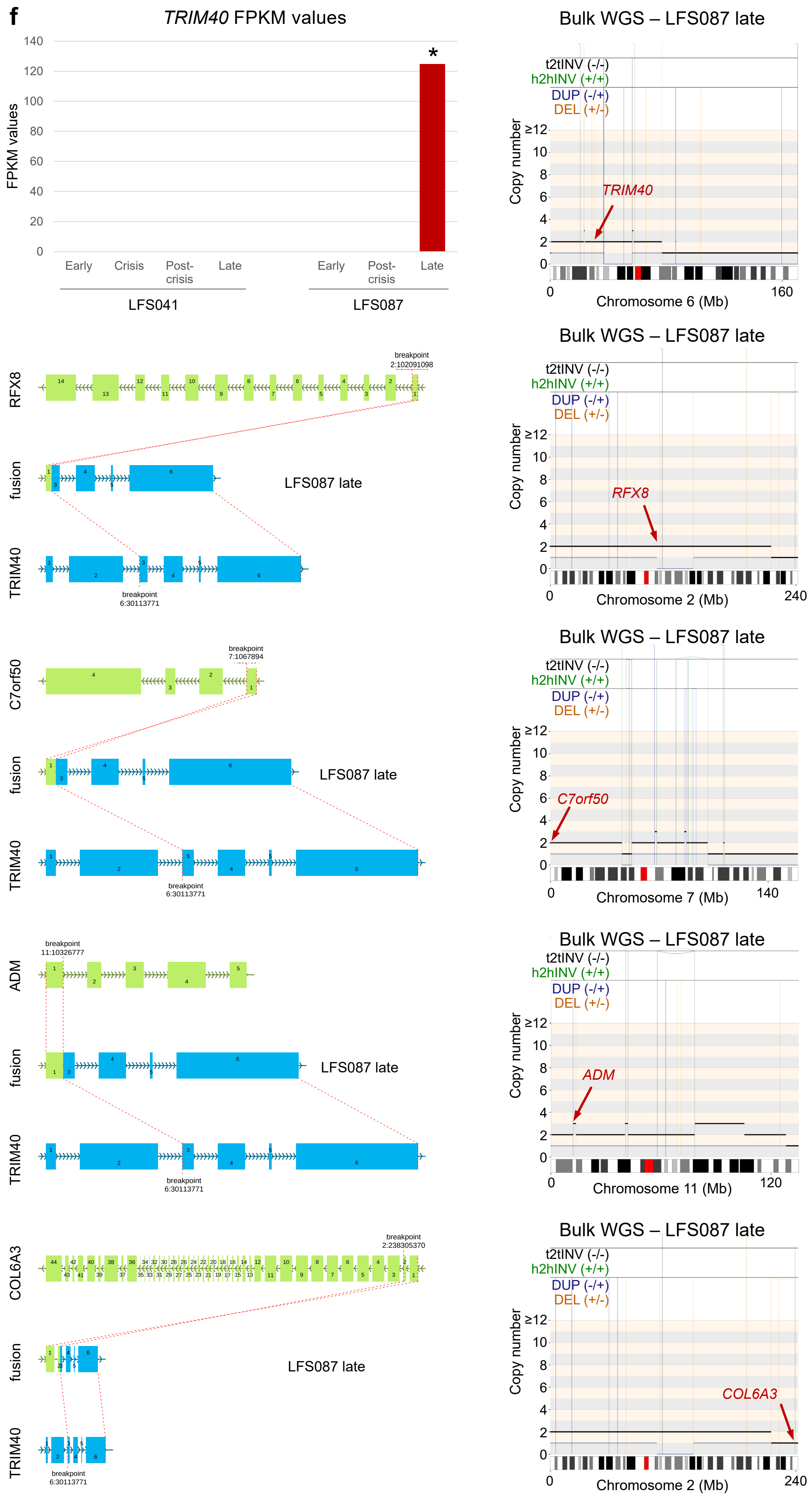
f
TRIM40 FPKM values
FPKM values
140
120
100
80
60
40
20
0
*
Early
Crisis
Post-crisis
Late
Early
Post-crisis
Late
LFS041
LFS087
Bulk WGS – LFS087 late
t2tINV (-/-)
h2hINV (+/+)
DUP (-/+)
DEL (+/-)
Copy number
≥12
TRIM40
Chromosome 6 (Mb)
0
160
RFX8
Chromosome 2 (Mb)
240
C7orf50
Chromosome 7 (Mb)
140
ADM
Chromosome 11 (Mb)
120
COL6A3
fusion
TRIM40
LFS087 late
breakpoint
2:102091098
6:30113771
7:1067894
11:10326777
2:238305370

Figure S5 f

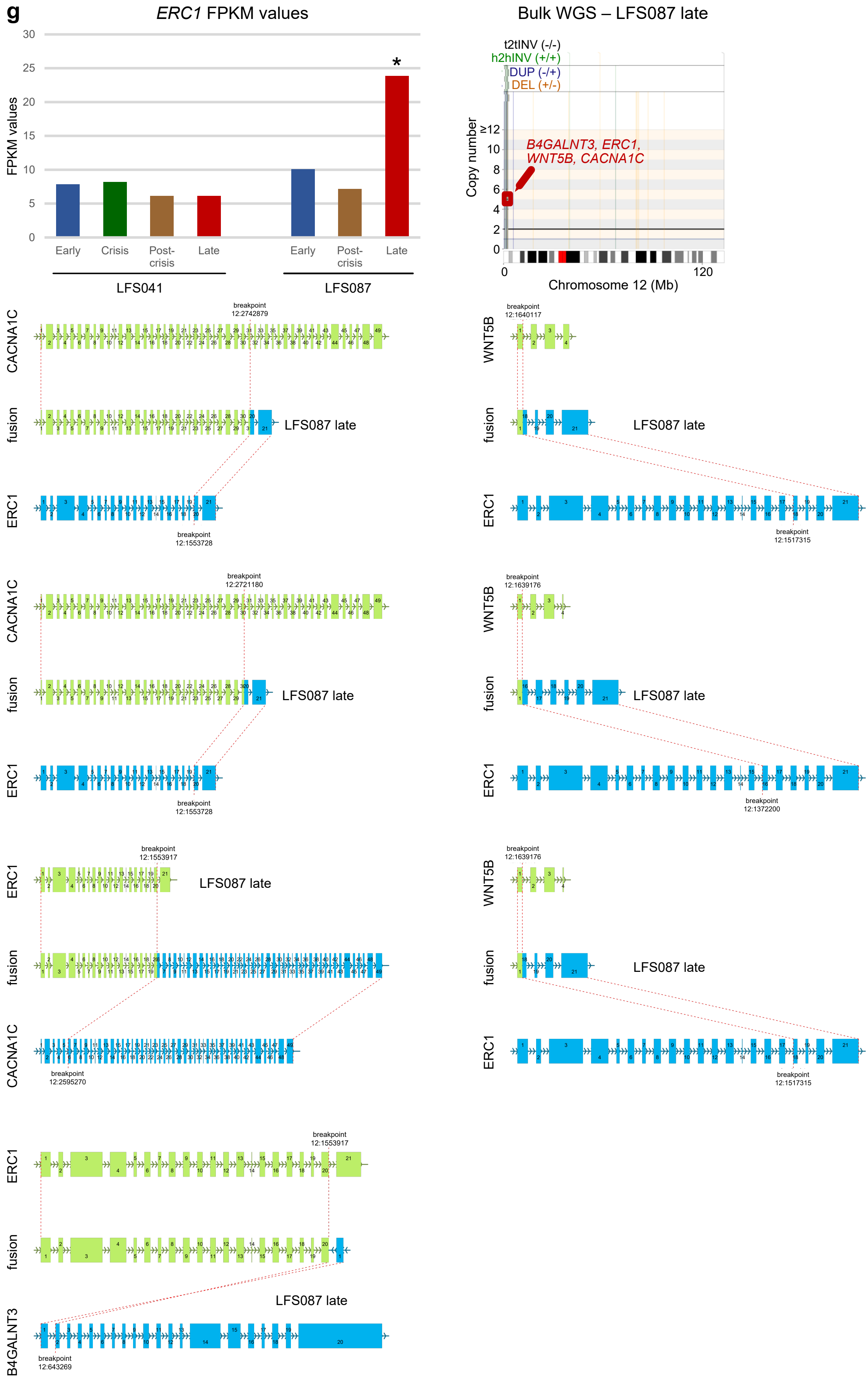
g
ERC1 FPKM values
FPKM values
30
25
20
15
10
5
0
*
Early
Crisis
Post-crisis
Late
Early
Post-crisis
Late
LFS041
LFS087
Bulk WGS – LFS087 late
t2tINV (-/-)
h2hINV (+/+)
DUP (-/+)
DEL (+/-)
Copy number
≥12
10
8
6
4
2
0
B4GALNT3, ERC1, WNT5B, CACNA1C
0
120
Chromosome 12 (Mb)
CACNA1C
fusion
ERC1
WNT5B
B4GALNT3
LFS087 late
breakpoint 12:2742879
breakpoint 12:1553728
breakpoint 12:1640117
breakpoint 12:1517315
breakpoint 12:2721180
breakpoint 12:1639176
breakpoint 12:1372200
breakpoint 12:1553917
breakpoint 12:2595270
breakpoint 12:643269

Figure S5 g

**<u>Figure S5.</u> Winning subclones show ecDNAs and gene fusions.**

**a, b** and **c.** display ecDNA structures identified in LFS041 p63 using AmpliconArchitect, based on bulk WGS data. *MYC* is highlighted in panel **a**, which corresponds to ecDNA shown in Figure 3H. **d.** shows an example of ecDNA structure identified for LFS087 p.195. **e.** Bar plot showing FPKM values for *COL5A1* in different passages of LFS041 and LFS087 cells, one of the fusion partners in the *COL5A1-SCMAP2* fusion (detected in LFS041 p.346 - late) and in the *COL5A1-GNAS* fusion (detected in LFS041 p.65 - post-crisis) detected by Arriba analysis from bulk RNA-seq. The asterisks indicate the patient and the passage where the fusion was detected. Arriba plots show the splice site of each gene fusion, along with the corresponding patient and passage. Right, ReConPlot from bulk WGS of LFS041 p.63 (post-crisis) shows the gene locus on the corresponding chromosome, as well as the different types of structural variations, with color coding indicating sites of deletions (orange), duplications (blue), head-to-head inversions (green) and tail-to-tail inversions (black). Total chromosome length is shown in megabase (Mb). Chromosomes annotated with an asterisk (*) indicate chromothriptic chromosomes. t2tINV: tail-to-tail inversion, h2hINV: head-to-head inversion, DUP: duplication and DEL: deletion. **f.** exhibits similar results to **e.** but for *TRIM40* and its different fusion partners, which includes the following gene fusions, *RFX8-TRIM40*, *C7orf50-RIM40*, *ADM-TRIM40* and *COL6A3-TRIM40*, all detected in LFS087 p.195 (late) by bulk RNA sequencing. **g.** exhibits similar results to **e.** but for *ERC1* and its fusion partners, which includes the following gene fusions, *CACNA1C-ERC1*, *WNT5B-ERC1* and *ERC1-B4GALNT3*, all detected in LFS087 p.195 (late) by bulk RNA sequencing. Stars on the bar plots show passages for which the gene fusion is detected in bulk RNA sequencing.

**a**

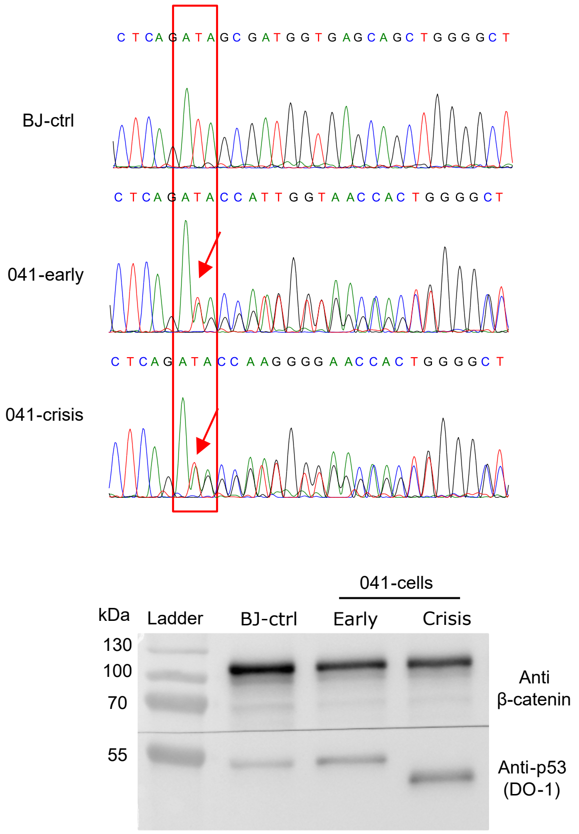


**b**

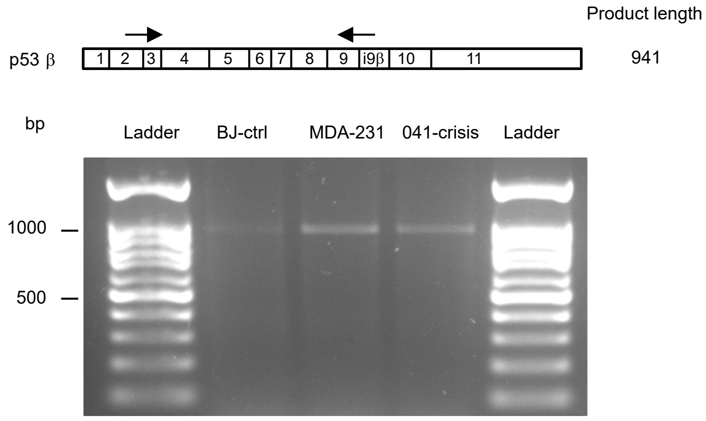


**c**

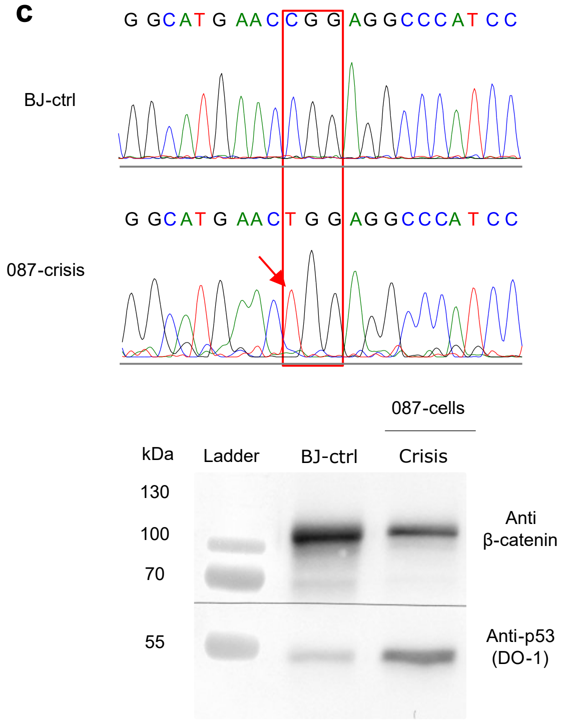


**d**

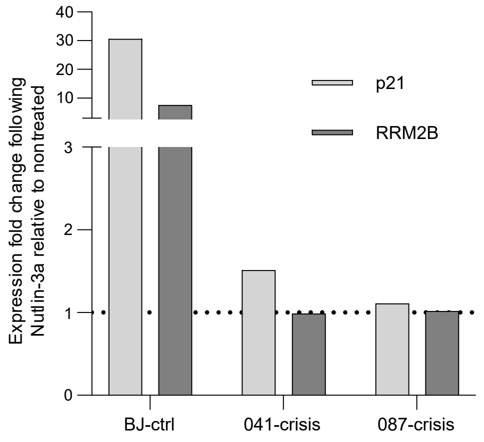


**e**

| LFS cell patient | 041-cells | 087-cells |
|---|---|---|
| **Heterozygous TP53 germline mutation** | Deletion in **Codon 184** GAT -> GAA | Missense in **Codon 248** CGG-> TGG |
| **Second p53 hit** | Functional loss of WT p53 size at protein level | Loss of WT Tp53 allele |

Figure S6 a-e

f

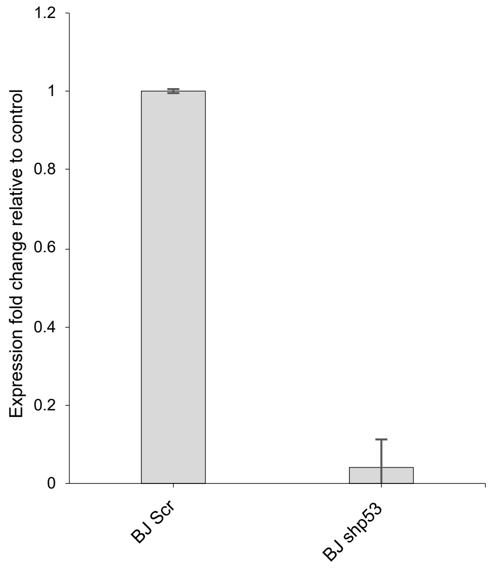


g

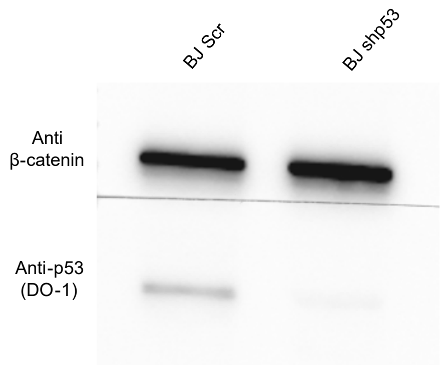


<u>Figure S6.</u> **Loss of WT p53 at the crisis passage in LFS cells and in BJ cells following p53 downregulation.**

**a. Upper panel -** DNA sequencing of the mutation site (red box) in 041-cells at early and crisis passages, red arrows point towards the deleted base. **Lower panel -** Western blot of p53 protein, detected by DO-1 antibody, in BJ control cells and 041-cells at early and crisis passages, β-catenin as loading control. **b. Upper panel -** Schema of p53β mRNA and the location of primers specific for p53β mRNA. **Lower panel** – The RT-PCR products were run on an ethidium bromide-stained agarose gel. BJ cells (negative control), MDA-321 (positive control cells expressing p53β) and 041-crisis passage cells. **c. Upper panel -** DNA sequencing of the mutation site (red box) in 087-cells at crisis passage, red arrow points towards the mutated base. **Lower panel -** Western blot of p53 protein, detected by DO-1 antibody, in crisis 087-cells, β-catenin as loading control. **d.** Fold change in expression analyzed by real-time PCR of p53 target genes (p21 and RRM2B) in BJ cells and in crisis LFS cells (041-crisis and 087-crisis) following p53 activation using 10μM nutlin-3a for 48 hours. **e.** Summary of the two LFS patient cells harboring a heterozygous germline mutation and the second hit detected at the crisis leading to loss of the WT p53. **f.** Fold changes in expression analyzed by real-time PCR of p53 mRNA measured in p53 downregulated cells relative to the expression in scr cells. GAPDH mRNA level used for normalization. The results are the summary of three independent experiments**. g.** Western blot of p53 protein in BJ control cells and following shp53. β-catenin as loading control.

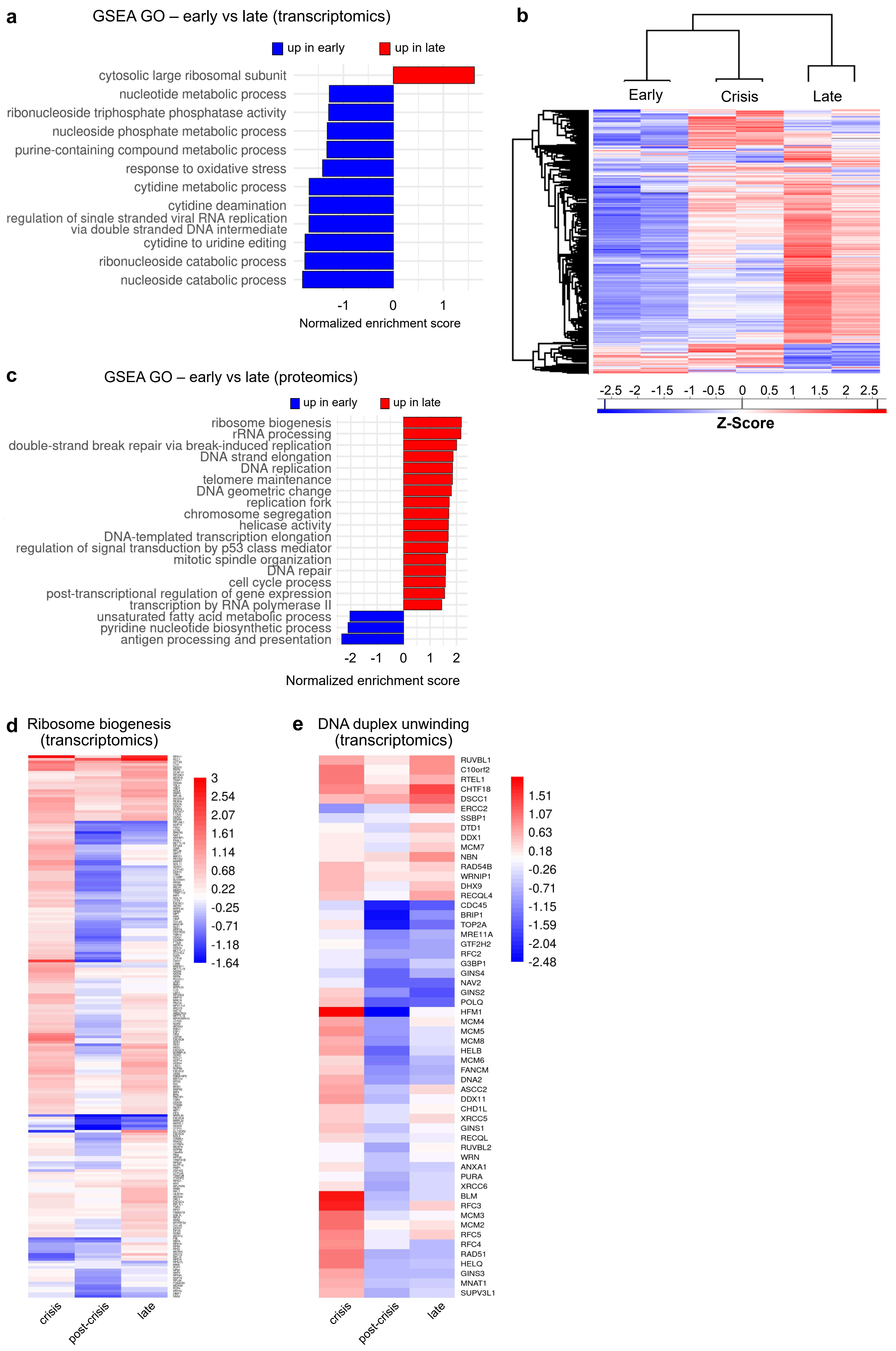
a
GSEA GO – early vs late (transcriptomics)
up in early
up in late
cytosolic large ribosomal subunit
nucleotide metabolic process
ribonucleoside triphosphate phosphatase activity
nucleoside phosphate metabolic process
purine-containing compound metabolic process
response to oxidative stress
cytidine metabolic process
cytidine deamination
regulation of single stranded viral RNA replication via double stranded DNA intermediate
cytidine to uridine editing
ribonucleoside catabolic process
nucleoside catabolic process
-1
0
1
Normalized enrichment score
b
Early
Crisis
Late
-2.5 -2 -1.5 -1 -0.5 0 0.5 1 1.5 2 2.5
Z-Score
c
GSEA GO – early vs late (proteomics)
up in early
up in late
ribosome biogenesis
rRNA processing
double-strand break repair via break-induced replication
DNA strand elongation
DNA replication
telomere maintenance
DNA geometric change
replication fork
chromosome segregation
helicase activity
DNA-templated transcription elongation
regulation of signal transduction by p53 class mediator
mitotic spindle organization
DNA repair
cell cycle process
post-transcriptional regulation of gene expression
transcription by RNA polymerase II
unsaturated fatty acid metabolic process
pyridine nucleotide biosynthetic process
antigen processing and presentation
-2
-1
0
1
2
Normalized enrichment score
d
Ribosome biogenesis (transcriptomics)
3
2.54
2.07
1.61
1.14
0.68
0.22
-0.25
-0.71
-1.18
-1.64
crisis
post-crisis
late
e
DNA duplex unwinding (transcriptomics)
RUVBL1
C10orf2
RTEL1
CHTF18
DSCC1
ERCC2
SSBP1
DTD1
DDX1
MCM7
NBN
RAD54B
WRNIP1
DHX9
RECQL4
CDC45
BRIP1
TOP2A
MRE11A
GTF2H2
RFC2
G3BP1
GINS4
NAV2
GINS2
POLQ
HFM1
MCM4
MCM5
MCM8
HELB
MCM6
FANCM
DNA2
ASCC2
DDX11
CHD1L
XRCC5
GINS1
RECQL
RUVBL2
WRN
ANXA1
PURA
XRCC6
BLM
RFC3
MCM3
MCM2
RFC5
RFC4
RAD51
HELQ
GINS3
MNAT1
SUPV3L1
1.51
1.07
0.63
0.18
-0.26
-0.71
-1.15
-1.59
-2.04
-2.48
crisis
post-crisis
late

Figure S7

**<u>Figure S7.</u> Proteomics and transcriptomics analyses identify critical pathways that drive chromothripsis in LFS fibroblasts.**

**a.** GSEA GO analysis showing key significantly enriched processes in LFS041 fibroblasts, comparing early (p.19) and late (p.346) passages using bulk RNA sequencing. **b.** Hierarchical clustering of differentially regulated proteins across early (p.19), crisis (p.27), and late (p.346) passages in patient LFS041. Clustering is based on Z-scores derived from mass spectrometry-based proteomics, highlighting distinct expression patterns. Proteins are grouped into clusters based on their upregulation (red) or downregulation (blue) across the three passages. **c.** GSEA GO analysis showing key significantly enriched processes in LFS041 fibroblasts, comparing early (p.19) and late (p.346) passages using mass spectrometry-based proteomics. **d, e.** Differential gene expression analysis of ribosome biogenesis **(d)** and DNA duplex unwinding **(e)** based on bulk RNAseq analysis in LFS041**.** Heatmaps display differentially expressed genes across early (p.19), post-crisis (p.65), and late (p.346) passages, with comparisons relative to early passage (p.19). The colour gradient represents the level of gene expression changes derived from NOIseq, with red indicating upregulation (in crisis, post-crisis, or late passages) and blue indicating downregulation as compared to early passage.

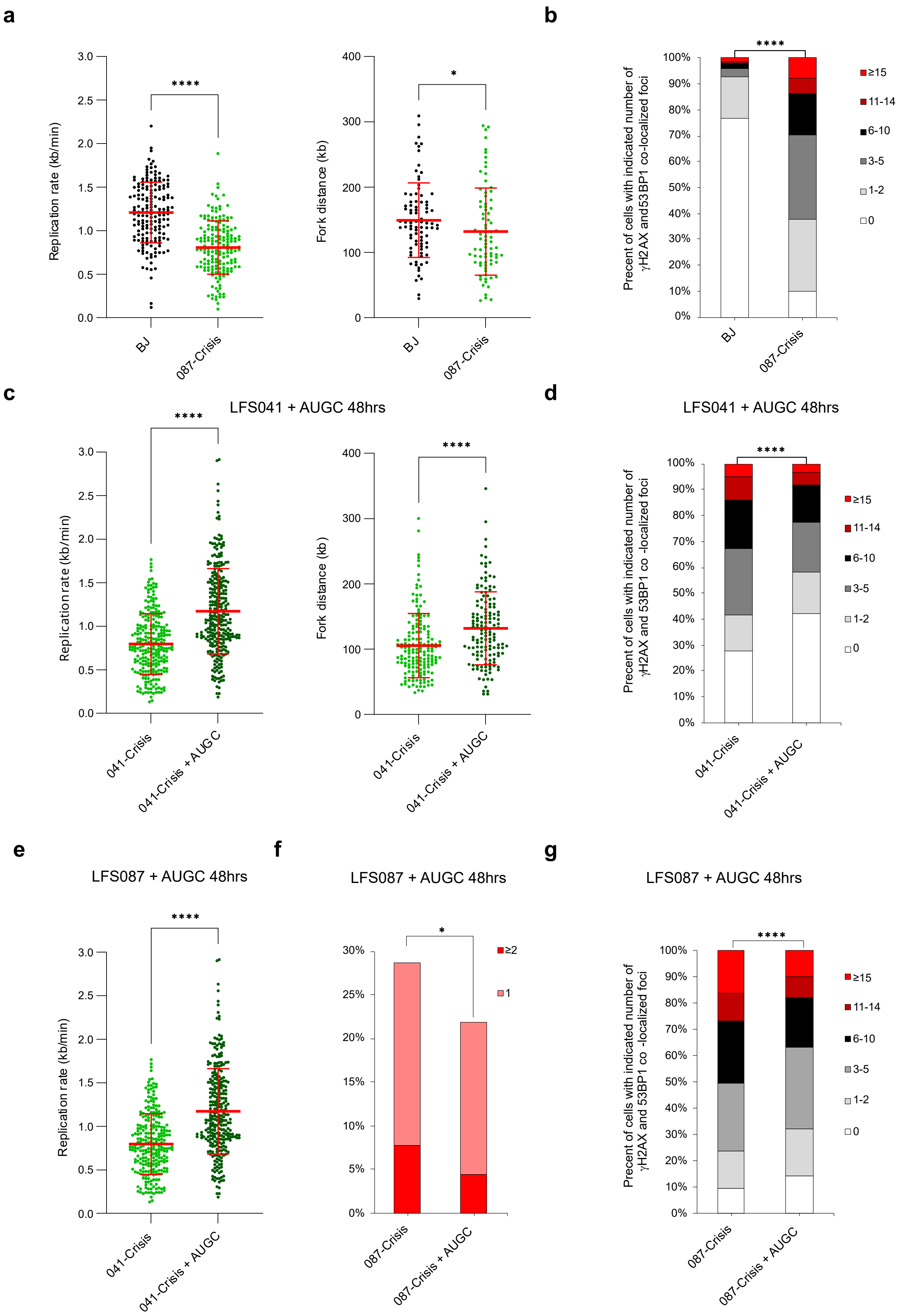
a
b
c
LFS041 + AUGC 48hrs
d
LFS041 + AUGC 48hrs
e
LFS087 + AUGC 48hrs
f
LFS087 + AUGC 48hrs
g
LFS087 + AUGC 48hrs
Replication rate (kb/min)
Fork distance (kb)
Precent of cells with indicated number of γH2AX and53BP1 co-localized foci
BJ
087-Crisis
041-Crisis
041-Crisis + AUGC
087-Crisis + AUGC
****
*
≥15
11-14
6-10
3-5
1-2
0
≥2
1

Figure S8 a-g

**h**

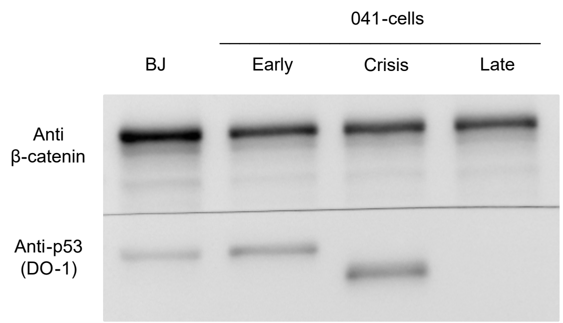


**i**

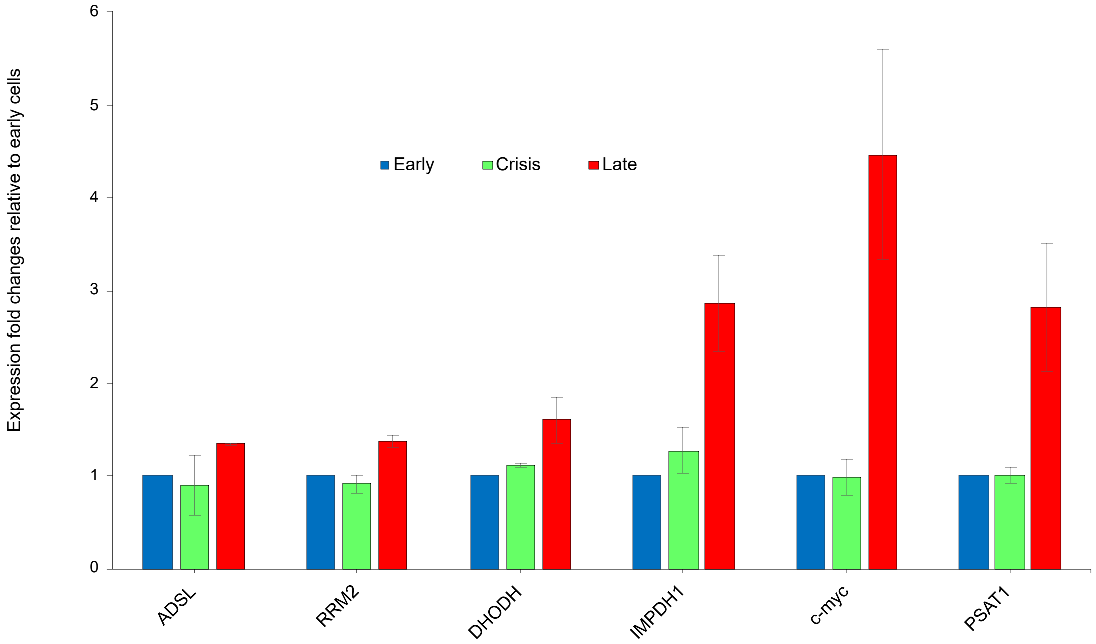


**j**

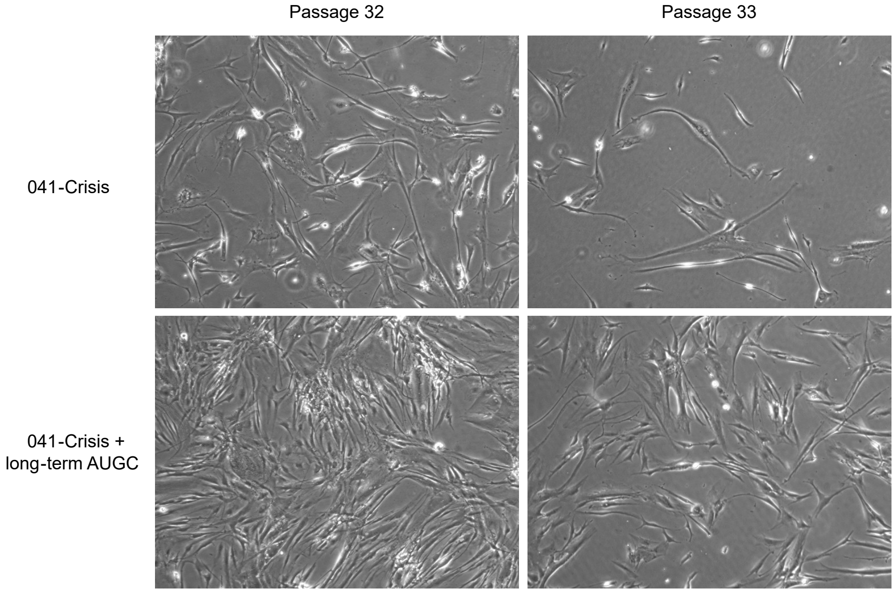


Figure S6 h-j

**<u>Figure S8.</u> Replication stress due to insufficient nucleotides leads to genomic instability in crisis LFS cells.**

**a. Left -** Fork rate (kb/min) distribution. Red lines indicating the mean±SD fork rate values for BJ control cells (1.2±0.3; n=171) and crisis passage cells of LFS087 (0.8±0.3; n=171). **Right -** Fork distance (kb) distribution. Red lines are the mean±SD fork distance values for BJ control cells (149±57; n=86) and crisis passage cells of LFS087 (132±67; n=80). The data are representative of two independent experiments with similar results. Statistical analysis was performed using Mann Whitney rank-sum test. **b.** Co-localization of γH2AX and 53BP1 foci for BJ cells (mean foci per cells 0.9; n=243), crisis passage cells of 087-cells (5.2, n=229). The data are representative of two independent experiments with similar results. Statistical analysis was performed using Mann Whitney rank-sum test done on the distribution. **c. Left -** fork rate (kb/min) distribution. Red lines indicating the mean±SD fork rate values of 041-crisis cells (0.8±0.4; n=255), 041-crisis cells supplemented with nucleosides for 48 hours (1.2±0.5; n=293). **Right -** Fork distance (kb) distribution. Red lines are the mean±SD fork distance values of 041-crisis cells (106±49; n=164), 041-crisis cells supplemented with nucleoside for 48 hours (132±56; n=131). The data are a summary of three independent experiments. Statistical analysis was performed using unpaired t-test.

**d.** Co-localization of γH2AX and 53BP1 foci for 041-crisis cells (mean foci per cells 4.9; n=325), 041-crisis cells supplemented with nucleosides for 48 hours (3.4, n=250). The data are representative of two independent experiments with similar results. Statistical analysis was performed using Mann Whitney rank-sum test done on the distribution. **e.** Fork rate (kb/min) distribution. Red lines indicating the mean±SD fork rate values for 087-crisis cells (0.8±0.3; n=293) and crisis 087-cells supplemented with nucleosides for 48 hours (1.1, n=112). The results are from one experiment. **f.** Percentage of cells with the indicated number of micronuclei in 087-crisis cells (n=542), 087-crisis cells supplemented with nucleosides for 48 hours (n=500). Chi-Square test of Independence was performed to compare the distribution of cells with micronuclei between the conditions. The data are a summary of three independent experiments. **g.** Co-localization of γH2AX and 53BP1 foci for 087-crisis (mean foci per cells 8.7; n=348), 087-crisis cells supplemented with nucleosides for 48 hours (6.3, n=348). The data are a summary of three independent experiments. Statistical analysis was performed using Mann Whitney rank-sum test done on the distribution.

**h.** Western blot of p53 protein, detected by DO-1 antibody, in BJ control cells and 041-cells at early, crisis and late passages. β-catenin used as loading control. **i.** Fold changes in expression analysed by real-time PCR of c-myc and its target genes, involved in de-novo nucleotide biosynthesis, in early, crisis and late passages. Bar values indicate the mean. GAPDH mRNA level used for normalization. The results are the summary of two independent experiments. **j.** Representative images showing the cellular morphology in 041-crisis cells and 041-crisis cells supplemented with long-term nucleotides. *P<0.05, **P<0.01, ***P<0.001 and **** P<0.0001.

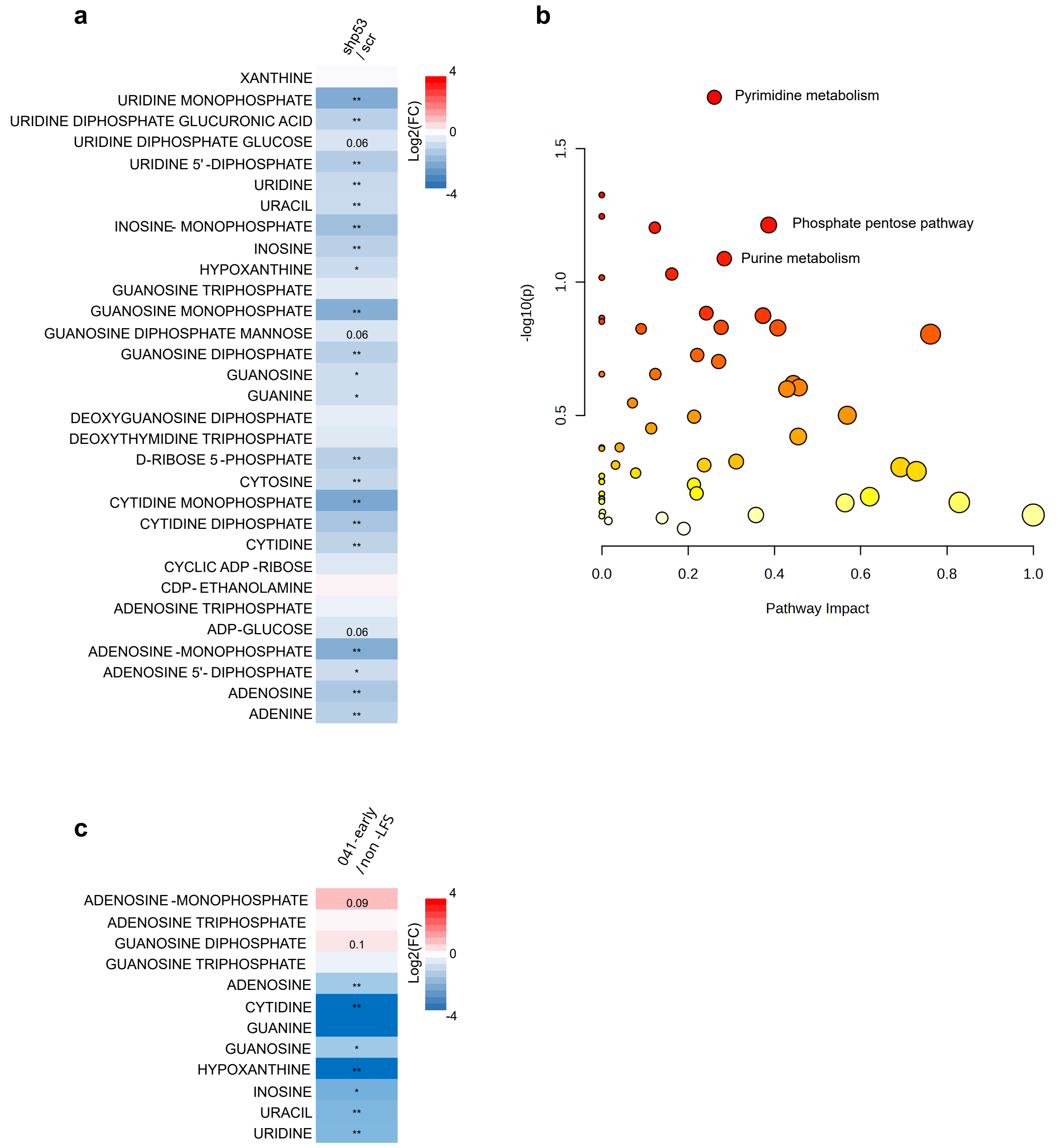
a
shp53 / scr
XANTHINE
URIDINE MONOPHOSPHATE
URIDINE DIPHOSPHATE GLUCURONIC ACID
URIDINE DIPHOSPHATE GLUCOSE
URIDINE 5' -DIPHOSPHATE
URIDINE
URACIL
INOSINE- MONOPHOSPHATE
INOSINE
HYPOXANTHINE
GUANOSINE TRIPHOSPHATE
GUANOSINE MONOPHOSPHATE
GUANOSINE DIPHOSPHATE MANNOSE
GUANOSINE DIPHOSPHATE
GUANOSINE
GUANINE
DEOXYGUANOSINE DIPHOSPHATE
DEOXYTHYMIDINE TRIPHOSPHATE
D-RIBOSE 5 -PHOSPHATE
CYTOSINE
CYTIDINE MONOPHOSPHATE
CYTIDINE DIPHOSPHATE
CYTIDINE
CYCLIC ADP -RIBOSE
CDP- ETHANOLAMINE
ADENOSINE TRIPHOSPHATE
ADP-GLUCOSE
ADENOSINE -MONOPHOSPHATE
ADENOSINE 5'- DIPHOSPHATE
ADENOSINE
ADENINE
**
**
0.06
**
**
**
**
**
*
**
0.06
**
*
*
**
**
**
**
**
0.06
**
*
**
**
Log2(FC)
4
0
-4
b
Pyrimidine metabolism
Phosphate pentose pathway
Purine metabolism
-log10(p)
1.5
1.0
0.5
0.0
0.2
0.4
0.6
0.8
1.0
Pathway Impact
c
041-early / non -LFS
ADENOSINE -MONOPHOSPHATE
ADENOSINE TRIPHOSPHATE
GUANOSINE DIPHOSPHATE
GUANOSINE TRIPHOSPHATE
ADENOSINE
CYTIDINE
GUANINE
GUANOSINE
HYPOXANTHINE
INOSINE
URACIL
URIDINE
0.09
0.1
**
**
*
**
*
**
**
Log2(FC)
4
0
-4

Figure S9

**<u>Figure S9.</u> Loss of WT p53 in BJ and in early passage LFS cells significantly reduces nucleotide metabolites.**

**a.** Heatmap representation of log fold changes (log FC) in metabolite levels, analyzed using LC-MS, in shp53 cells compared to the level in scr cells. The comparisons include six biological repeats. Red - increased metabolite levels compared to the indicated condition. blue - decreased metabolite levels compared to the indicated condition. P-value for each comparison calculated using one-sided t-test and indicated in each box (*$p < 0.05$, **$p < 0.01$, the value $<0.1$ is indicated). **b.** Pathway analysis using MetaboAnalyst for untargeted metabolomics data in shp53 cells compared to scr cells. The x-axis represents the pathway impact value, the y-axis shows the -log10(p-value). Each dot corresponds to a metabolic pathway, with its size proportional to the pathway impact. P-value represented by the color and transition from yellow to red reflects a decreased P-value. **c.** Heatmap representation of log fold changes (log FC) in metabolite levels in LFS041 early cells compared to non-LFS cells. At least 4 biological repeats from each group. P-value for each comparison calculated using one-sided t-test and indicated in each box (*$p < 0.05$, **$p < 0.01$, the value $<0.1$ is indicated).

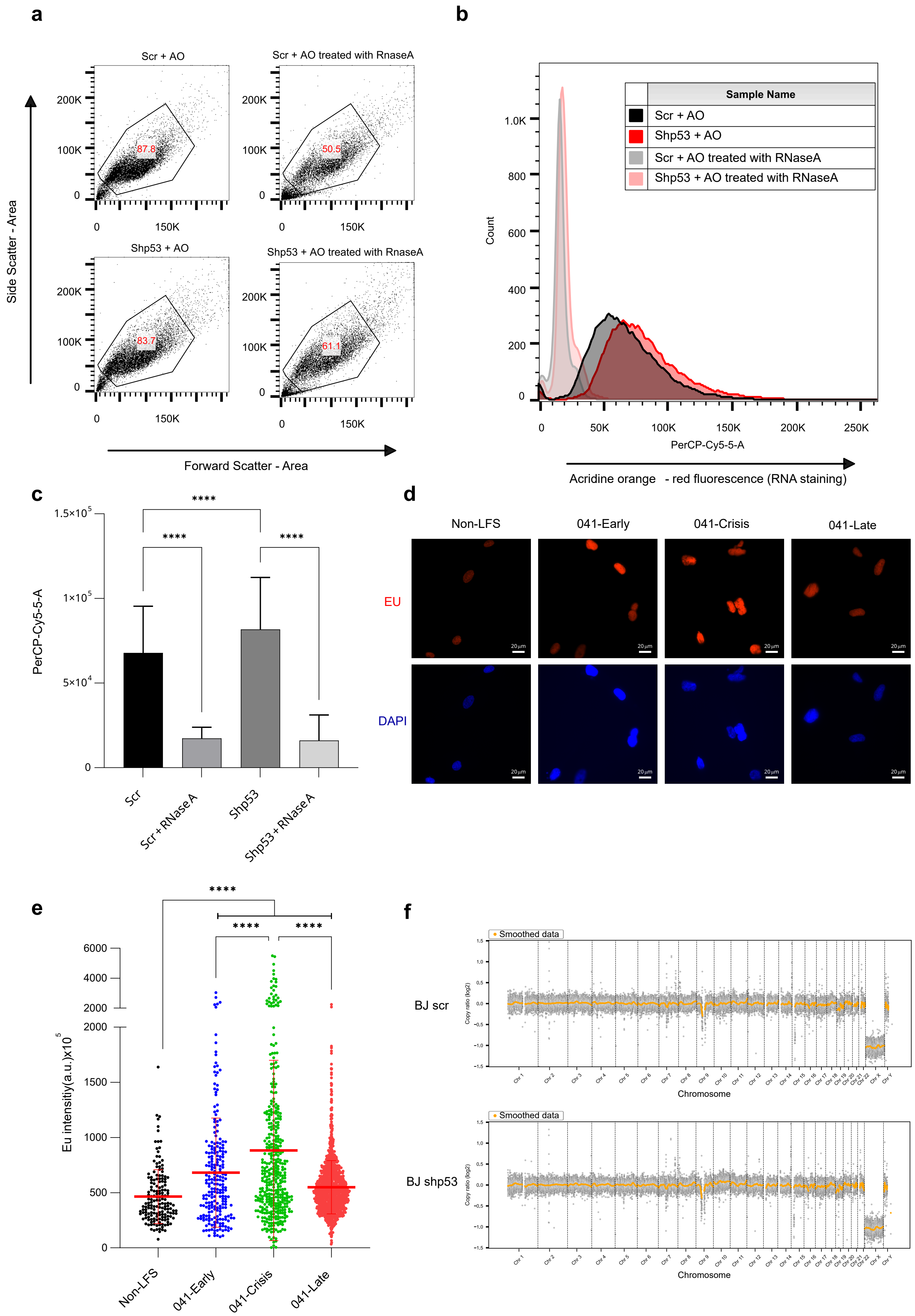


Figure S10 a-f

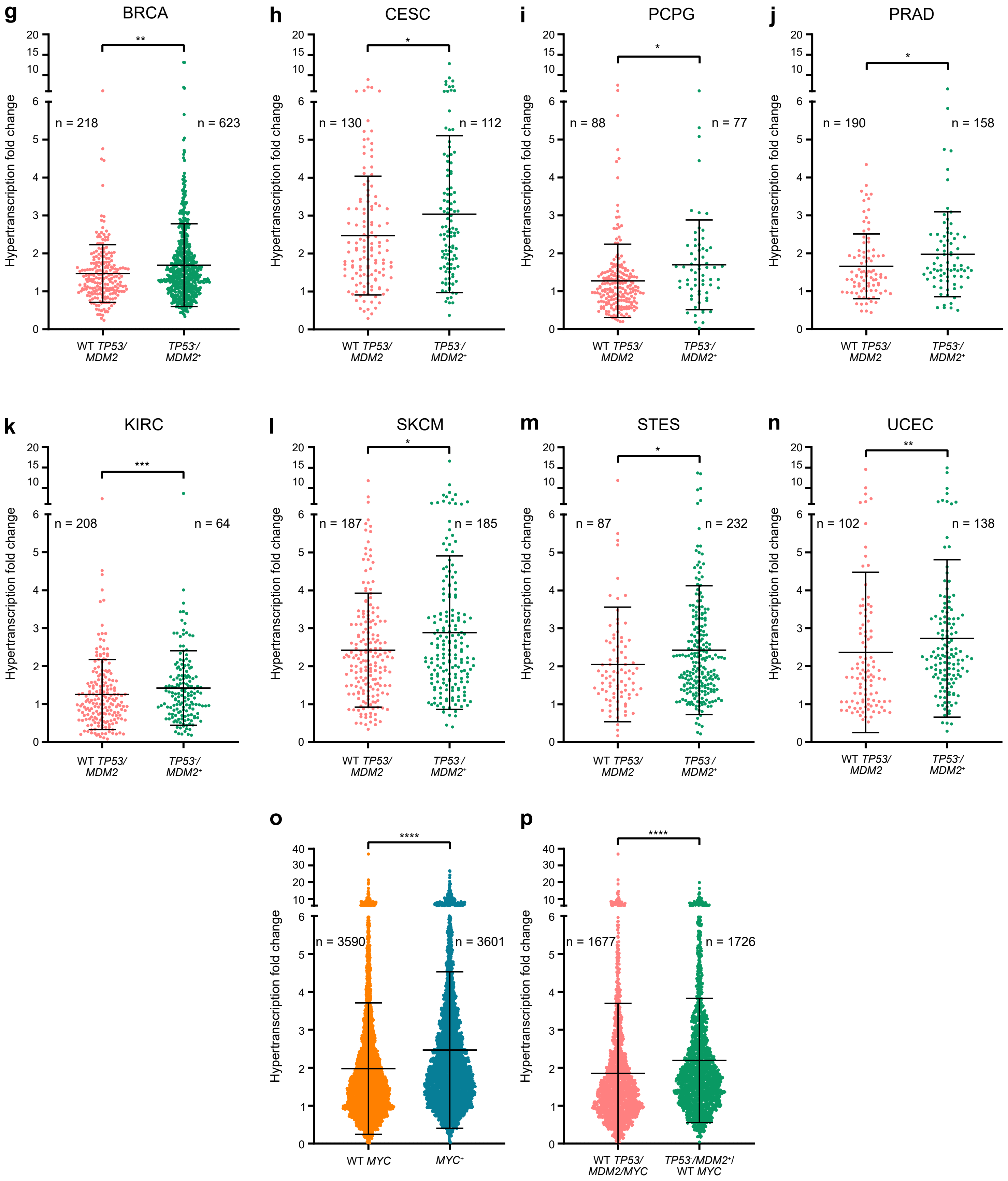


Figure S10 g-p

**<u>Figure S10.</u> p53 loss leads to increased transcription.**

**a.** Flow cytometry gating strategy based on side scatter area (SSC-A) and forward scatter area (FSC-A) for scr and shp53 cells stained with Acridine orange (AO) (**left**), Cell were also stained with AO and treated with RNaseA to validate the RNA specificity of the red AO fluorescence (**right**). The percent of gated cells is indicated in red.

**b**. Flow cytometry analysis of red fluorescence, emitted from AO, in the same conditions shown in a. **c.** Quantification of red fluorescence data shown in a and b for scr (mean 67753, n=17607), scr + RNaseA (17288, n=10092), shp53 (81638, n=16739) and shp53 + RNaseA (16078, n=20000). **** P<0.0001 calculated by one-Way Anova test. **d.** Representative images EU stanning (red) in non-LFS and in 041-cells at early, crisis and late passages. **e**. Quantification of nuclear EU intensity for non-LFS cells (mean intensity 464±245; n=164), 041-early cells (682±494; n=226), 041-crisis cells (883±816, n=406) and 041-late cells (550±241, n=1513). The data are a summary of two independent experiments. Statistical analysis was performed using one-Way Anova test. **f.** P53 downregulation did not result in aneuploidy. Copy number fold change across the genome in scr (upper panel) and shp53 cells (lower panel) analyzed using low-pass whole genome sequencing. **g-n.** Tumor type-specific hypertranscription fold-change score, showing significant differences between *TP53/MDM2* WT vs *TP53-/MDM2+* tumors. BRCA: breast invasive carcinoma, CESC: cervical squamous cell carcinoma and endocervical adenocarcinoma, KIRC: kidney renal clear cell carcinoma, PCPG: pheochromocytoma and paraganglioma, PRAD: prostate adenocarcinoma, SKCM: skin cutaneous melanoma, STES: stomach and esophageal carcinoma, UCEC: uterine corpus endometrial carcinoma. **o.** Hypertranscription fold-change in the pan-cancer cohort segregated by *MYC* status: WT *MYC* (mean ± SEM: 1.98 ± 0.03, n=3590) and *MYC+* (mean ± SEM: 2.47 ± 0.03, n=3601).**p.** Hypertranscription fold-change for all tumors without gain of *MYC*, comparing *TP53-/MDM2+*/WT *MYC* (mean ± SEM: 2.2 ± 0.04, n=1726) to WT *TP53/MDM2/MYC* (mean ± SEM: 1.85 ± 0.05, n=1677) to test for the effect of p53 in the context of balanced *MYC*. Each circle represents a tumor sample, with the total number of samples per group indicated on the plot. Data are presented as mean ± SEM. Statistical significance was assessed using a non-parametric t-test (Mann-Whitney test) to compare two independent groups, as the data were not normally distributed. P-values lower than 0.05 were considered statistically significant (*p < 0.05, **p < 0.01, ***p < 0.001, **p < 0.0001).
